# Large RNA polymerase II condensates are promoter-centric assemblies associated with early stages of transcription at all expressed genes

**DOI:** 10.64898/2026.05.25.727590

**Authors:** Jelena V. Bogdanović, Jonah Galeota-Sprung, Amoldeep S. Kainth, Filmon Medhanie, Alisha Budhathoki, Victoria Pappas, Salman F. Banani, Jan-Hendrik Spille, Alexander J. Ruthenburg

## Abstract

Transcriptional condensates concentrate the machinery required for RNA polymerase II mediated transcription. These structures range from numerous small, short-lived species, to a handful of larger, stable assemblages. Large condensates have been implicated in driving potent transcription of several super-enhancer regulated genes, yet the underlying mechanisms and the range of their client genes remain unclear. Here, we developed a biochemical approach which combines density gradient centrifugation and affinity purification to partially purify large transcriptional condensates from nuclei, allowing systematic characterization of their nucleic acid components. We find that transcriptional condensate isolates engage thousands of gene promoters and harbor the nascent transcriptome, but do not stably co-purify with distal enhancers. Binding patterns of RNA polymerase II within condensates suggest these structures could facilitate promoter escape and promoter-proximal pause release. Together, our work supports a promoter-centric condensate organization and paves the way towards understanding the functional link between condensate architecture and nascent transcription.

## INTRODUCTION

Transcription is not uniformly distributed within eukaryotic nuclei. Instead, it localizes to discrete nuclear bodies with high concentrations of the transcriptional machinery. These nuclear bodies have been described as transcription factories,^1,2^ hubs,^3–5^ and transcriptional condensates, as liquid-liquid-phase separation properties of their key constituents suggested a potential mechanism for their formation.^6–10^ The term ‘transcriptional condensate’ has since acquired wide usage agnostic of the precise physical properties of their formation,^11–14^ which nevertheless remain very active avenues of inquiry.^15–20^ Transcriptional condensates are membraneless nuclear bodies enriched for RNA polymerase II (Pol II), the transcriptional coactivators Mediator and Brd4, several general transcription factors, and pause release factors.^7,10,16,17,20,21^ They have been proposed to promote efficient gene expression.^7,10,21–24^ Despite intensive interest, the range of genes subject to this form of regulation and precise mechanisms of transcriptional activation promoted by condensates remain poorly understood. The prevailing model entails that condensates nucleate on super-enhancer DNA and form transient contacts with promoters of target genes, thereby supporting transcriptional output at select lineage-defining genes.^7,8,22,23,25–28^ Other studies have implicated transcriptional condensates in housekeeping gene regulation,^29^ and earlier work on RNA polymerase I-III transcription factories suggested a broad role for these structures in genome-wide nascent transcription.^30^ Thus, it remains unresolved whether transcriptional condensates primarily engage super-enhancers or other DNA elements; whether they are specialized for select gene classes or employed broadly across the active genome; and whether their main transcriptional role is initiation, pause release, early or general elongation.

A major obstacle to addressing these questions is that transcriptional condensates have primarily been studied *in situ.* Live-cell microscopy has revealed two classes of transcriptional condensates: numerous transient and small clusters (<100 nm in diameter), versus a more rare, stable population of large condensates (>200 or 300 nm, depending on the reference).^10,23^ Imaging has been essential for defining their dynamics,^7,10,23,31^ molecular composition,^7,17,20^ and demonstrating causal relationships between condensate formation and transcriptional output at specific genes.^4,10,21,23,24,32,33^ However, imaging alone requires a candidate-based approach and provides limited access to the DNA and RNA associated with transcriptional condensates.

Recent *in situ* proteomics and soluble nuclear extract preparations using coacervation properties of Brd4 or Med1 intrinsically disordered regions (IDRs) have begun to define condensate protein constituents.^17,20,34^ However, due to the inability to biochemically enrich the comparatively rare larger transcriptional condensates from the multitude of smaller ones, it has remained impossible to systematically determine the genomic elements and genes engaged by these structures, and the RNA species they contain.

Given the high stability of a subset of larger transcriptional condensates,^10^ which can persist for hours,^35^ we sought to biochemically isolate them. Previously, larger and similarly sized cytosolic and nuclear bodies have been biochemically isolated.^36–39^ Earlier isolation of much smaller transcription factories, with average sizes of 80-100 nm, suggested that these assemblies can survive mild nuclease- and protease-based preparations.^30,40^ We therefore sought to exploit the increased stability, size, and density of larger transcriptional condensates for biochemical enrichment from native mouse embryonic stem cell (mESC) nuclei. To that end, we developed POCEMON (<u>Po</u>l II <u>c</u>ondensate <u>e</u>nrichment and <u>m</u>apping <u>o</u>f <u>n</u>ucleic acids), a combined size- and density-based separation of sonicated nuclear material with affinity purification of Pol II- or Mediator subunit Med1-containing nuclear particles. Using multiple orthogonal approaches, we find that POCEMON-enriched transcriptional condensate isolates (TCIs) appear representative of larger endogenous transcriptional condensates. Systematic mapping of large TCI-associated DNA reveals promoter-centric structures which broadly engage all classes of genes in direct proportion to their nascent expression, with little evidence of stable distal enhancer engagement. This is distinct from the expectation of previous models that proposed an integral scaffolding role of distal enhancers in transcriptional condensates.^7,8,10,22,23,25,26,28^ Distributions of Mediator, polymerase, and RNA within TCIs are consistent with a model in which core Mediator recruits promoters to transcriptional condensates, and polymerase rapidly clears the promoter, entering early phases of elongation, with the nascent transcripts remaining physically tethered. The limited nuclear volume that these condensates occupy, and the wide range of promoters recovered in our preparations suggest marked heterogeneity in the set of genes transcribed by the set of large transcriptional condensates in one cell at a given time. Despite this heterogeneity, established three-dimensional juxtaposition of DNA elements is correlated with TCI co-occupancy. An accompanying manuscript (Pandey et al.) further supports our observations by super-resolution imaging, showing that transcriptional condensates contain multiple dense chromatin nanodomains with promoter-specific chromatin marks, whose number per large condensate corresponds to the number of TCI DNA elements establishing multiway interactions. Moreover, Pandey et al. observe both initiation- and elongation-associated phosphoisoforms of Rbp1 within large condensates, consistent with our observation of TCI Pol II entry into productive elongation. Together, our work provides insights into genomic engagement of transcriptional condensates and enables finer understanding of mechanisms by which condensates contribute to transcription.

## RESULTS

### Partial purification of medium and large transcriptional condensates from mESC nuclei

To identify genomic DNA and RNA associated with transcriptional condensates, we developed POCEMON, a combined gradient fractionation- and affinity purification-based approach designed to enrich material from the larger regime of the transcriptional condensate distribution (>200 nm) (**Figure 1A-C**). POCEMON utilizes brief sonication to fragment purified nuclei into a continuum of particle sizes, which can then be separated by differential centrifugation steps to enrich nuclear bodies of the desired size regime (**Figure 1C-D**). Previous native preparations of other sub-nuclear bodies indicate some structures have a surprising resilience to this form of nuclear disruption.^36–38,40^ We noted no difference in the apparent stability and overall composition of transcriptional condensates with crosslinking (**Figure 1E, S1A-B**) and thus proceeded with native preparations to reduce background. During sonication and in a brief incubation afterward, nuclear material is subjected to HaloTag-based biotinylation of Rpb1, largest subunit of Pol II, or separately Mediator complex subunit Med1, for downstream affinity purification (**Figure 1C, S1C**). Following HaloTag biotinylation, sonicated nuclear isolate is subjected to centrifugation through a sucrose cushion to deplete nucleoli and remove large nuclear debris (**Figure 1C**). The resulting nucleoli depleted fraction (NdF) is then subjected to further size- and density-based separation using Percoll gradient fractionation (Size/density gradient 1, **Figure 1C**). The densest fraction, fraction 10 (F10), contains larger condensates, whereas smaller/less dense species are distributed through earlier fractions (**Figure 1D-F**). This distribution also roughly corresponds to isolated DNA fragment length (**Figure S1D**).

**Figure 1.**
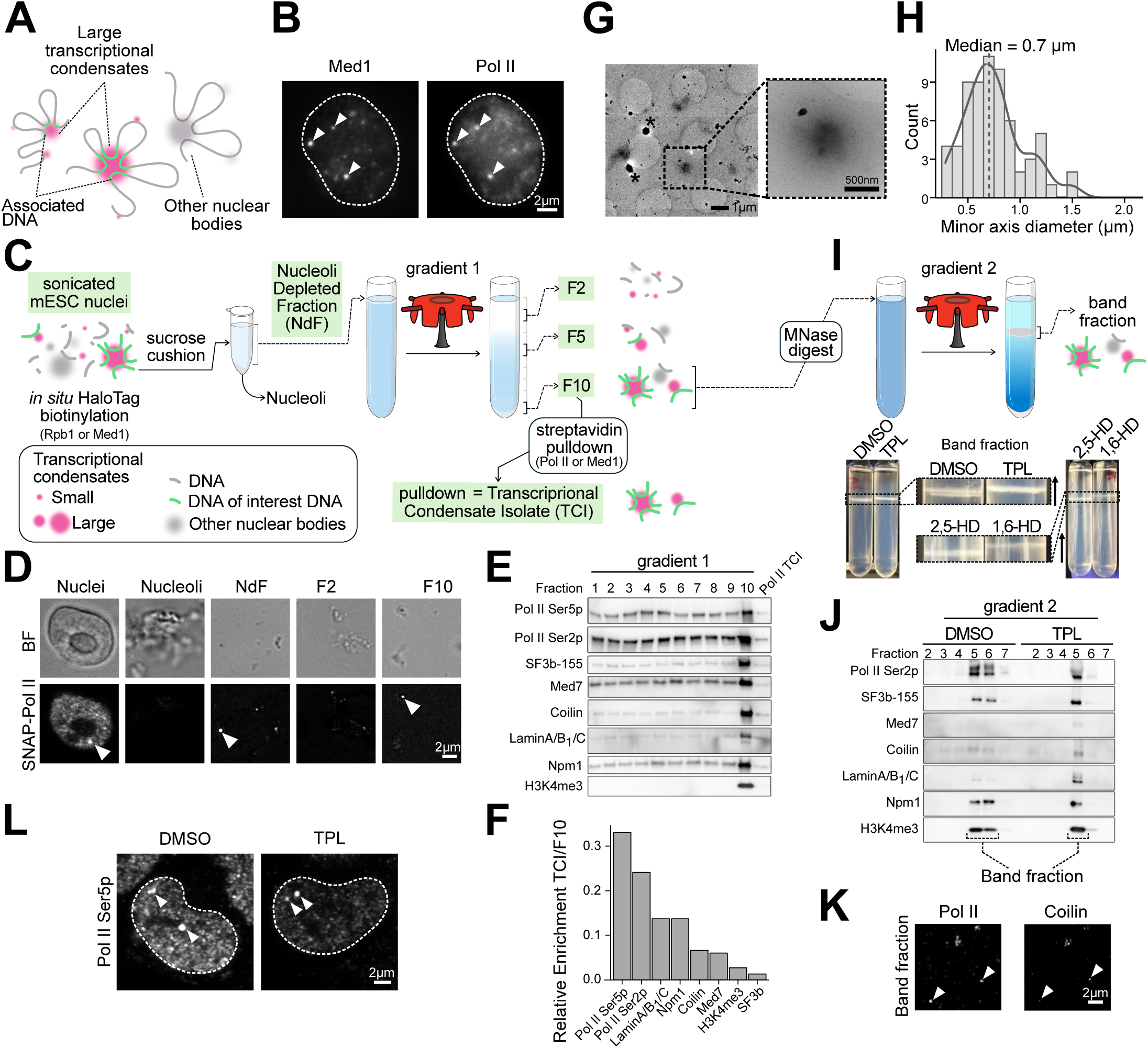
POCEMON enriches large transcriptional condensates from mESC nuclei. **(A)** Conceptual model of large transcriptional condensates containing Pol II/Mediator and associated genomic DNA, shown alongside other nuclear bodies. **(B)** Representative image of Halo-Med1 and SNAP-Pol II transcriptional condensates in the mESC nucleus (dashed white line). **(C)** Schematic of the POCEMON biochemical approach. Sonicated mESC nuclei carrying HaloTag-labeled Rpb1 or Med1 are subjected to biotinylation and sucrose cushion to generate a nucleoli-depleted fraction (NdF). This material is then separated by a primary Percoll size/density gradient, and ten volumetric fractions are collected. F10 is then subjected to streptavidin pulldown to furnish the transcriptional condensate isolate (TCI). **(D)** Brightfield (BF, upper panels) and confocal SNAP-Pol II fluorescence images (lower panels) across the POCEMON protocol. Arrowheads indicate transcriptional condensate-like structures. **(E)** Immunoblot analysis of fractions from primary gradient and of the final TCI preparation. **(F)** Densitometry of the TCI fraction relative to F10 from primary gradient, indicating relative enrichment of different proteins/marks in the final Pol II TCI preparation. **(G)** Representative cryo-electron micrograph of F10 showing rounded, electron-dense structures. Inset shows a higher-magnification view of a representative particle. Asterisks denote ice. **(H)** Size distribution of spheroids measured from segmented electron tomograms and plotted as minor-axis diameter. **(I)** Top, schematic of secondary density-gradient assay to assess perturbation-dependent changes in transcriptional condensate sedimentation. The F10 material was treated with MNase and resolved by a secondary higher-density Percoll gradient to separate a discrete band fraction. Bottom, images of representative secondary gradient preparations following indicated treatments: 10 μM triptolide (TPL) or the equivalent volume of DMSO added to culture medium for 60 minutes prior to nuclear isolation; 10% (w/v) 1,6-hexanediol (1,6-HD) or 2,5-hexanediol (2,5-HD) for 12 minutes prior to nuclear isolation. **(J)** Immunoblot analysis of fractions from secondary gradient under DMSO and TPL conditions. The band fraction is highlighted and probed for the indicated factors. **(K)** Immunofluorescence of the recovered band fraction stained for Pol II and Coilin. Arrowheads indicate condensate-like puncta. **(L)** Representative images of Pol II-CTD Ser5p immunofluorescence in control and triptolide-treated cells. Arrowheads indicate condensate-like puncta.

Rather than purification to homogeneity, this fractionation step affords enrichment of large transcriptional condensates alongside other nuclear bodies of similar size and density, including Cajal bodies^36^. Brightfield and confocal imaging of fractions throughout the preparation show that large SNAP-Pol II-positive puncta remain detectable after fractionation and are particularly evident in F10, consistent with retention of larger condensate material (**Figure 1D**). Immunoblot analysis (**Figure 1E**) shows that initiating- and elongating-Pol II phosphoisoforms (Ser5p and Ser2p, respectively)^41^ display a wide distribution across the gradient, with enrichment toward the terminal fraction, as do Med1 and residual Npm1, a factor enriched in the nucleolus but also present in the nucleoplasm^42,43^. In contrast, Cajal body marker Coilin^44,45^ and lamins A, B_1_, and C remain associated primarily with larger structures, as evidenced by their predominant sedimentation into F10. This is consistent with the more uniform size of Cajal bodies, ranging from 0.5-1 μm in diameter^46^, which corresponds to the size of the largest transcriptional condensates. We note that nuclear speckle marker SRRM1^47,48^, Coilin, and Npm1, were relatively depleted in the NdF prior to the gradient (**Figure S1E**). All proteins we assayed for were detectable in the terminal fraction (**Figure 1E, S1G**), consistent with the interpretation that this fraction contains a mixed population of large/dense nuclear assemblies rather than exclusively transcriptional condensates. Thus, the first gradient resolves transcriptional condensates of differing sizes/density, but the material in F10 remains compositionally heterogeneous.

To further enrich for transcriptional condensates from the pool of large nuclear bodies, F10 is subjected to affinity purification using streptavidin pulldown from Pol II or Med1 tagged lines—this is the version of the preparation we refer to as POCEMON (**Figure 1C**). We refer to the POCEMON-isolated material as *Transcriptional Condensate Isolate (TCI)*, since ratiometric immunoblot analysis indicates relative enrichment of known transcriptional condensate factors over other subnuclear body factors (**Figure 1E-F**). Notably, we observed greatest enrichment for the initiation-associated Pol II (Ser5p), consistent with live-cell microscopy observations of this phosphoisoform in large condensates.^49^

To confirm enrichment of expected nuclear body sizes and visualize their ultrastructure, we performed cryo-electron tomography on F10-derived samples. Search maps identified rounded, electron-dense structures, and micrographs revealed spheroid condensate-like assemblies on the submicron scale (**Figure 1G**). These observations provide orthogonal structural support that the terminal fraction contains discrete mesoscale assemblies rather than dispersed macromolecular material. Quantification of segmented tomograms further defined the size distribution of structures recovered in F10, revealing assemblies spanning ∼300–1500 nm with a median diameter of ∼700 nm (**Figure 1H**). This size range is consistent with capture of the desired >200 nm-sized transcriptional condensates, including >500 nm large histone locus bodies^50,51^ (HLBs, overlapping largest transcriptional condensates) and Cajal bodies^10,23,35^. Absence of structures smaller than 300 nm and larger than 1500 nm further validates the efficacy of our nucleoli removal and gradient 1 separation (**Figure 1C**), thereby confirming the size regime in the late stage of our preparations.

For further validation that our preparations retain features of transcriptional condensates present in living cells, we sought to perturb condensates prior to their biochemical isolation. To assess whether physical properties of the crude transcriptional condensates change upon perturbation, we introduced a second, higher-density Percoll gradient as an analytical step (**Figure 1I**), where bulk condensate material forms a visible band whose position could be compared between conditions. Prior to application to this second gradient, we lightly digested F10 with micrococcal nuclease (MNase), following a prior Cajal body preparation’s use of nuclease.^36^ This approach is intended to report perturbation-dependent changes in the effective size and/or density of transcriptional condensate-associated material, as these parameters determine migration. Immunoblotting of fractions from this second gradient showed that the band fraction was of similar composition to F10 (**Figure 1J**). Imaging of the band fraction revealed Pol II-positive assemblages (**Figure 1K**), whereas Coilin was comparatively sparse and restricted to a subset of Pol II puncta, perhaps suggesting physical linkage consistent with spatial proximity observed in nuclei (**Figure S1F**).^35,52^ We reasoned that 1,6-hexanediol treatment of cells, which can disrupt certain condensates,^7,9,10,53,54^ would lead to an upward shift of the band fraction. Consistent with this expectation, the entire band fraction, including Pol II, was shifted upon 1,6-hexanediol treatment, relative to the 2,5-hexanediol control (**Figure 1I, S1G**). As a more specific cellular perturbation, we used triptolide (TPL), which acutely blocks transcription initiation through covalent inhibition of TFIIH promoter melting activity,^55^ while also inducing proteasomal degradation of Rpb1, the largest subunit of the Pol II complex, on an hour timescale.^55,56^ We therefore expected prolonged TPL treatment of the living cells before isolation to alter the composition of transcription-associated Pol II assemblies. Immunofluorescence imaging and immunoblot analysis confirmed that global nuclear Pol II levels are reduced after TPL treatment relative to the DMSO control, although large condensates are still detectable (**Figure 1L, S1E,G**). To assess TPL-induced changes in transcriptional condensate material, we applied the two-gradient approach. Surprisingly, TPL-associated shift of the band fraction resembled the 1,6-hexanediol shift in its magnitude and with similar behavior in both native and crosslinked preparations (**Figure 1I-J, S1A-B**). This result is consistent with altered physical properties of the recovered transcriptional condensate-associated material upon transcriptional inhibition and suggests that Pol II-containing material is either a major component of F10, or that other co-purifying nuclear bodies may be physically associated with transcriptional condensates.

Together, these data establish POCEMON as a biochemical approach that enriches endogenous large transcriptional condensates from mESCs, while retaining their gross nuclear properties. Compared to their cellular counterparts, their size distribution is similar by both light microscopy and cryo-ET, they are similarly most enriched for Pol II Ser5p, and cellular treatments with a pleiotropic condensate disrupting agent or a transcriptional initiation inhibitor cause measurable sedimentation changes of crude transcriptional condensates. The resulting condensate-enriched fraction after streptavidin capture (TCI) is suitable for downstream nucleic-acid sequencing to define their composition.

### POCEMON identifies genomic regions associated with endogenous transcriptional condensates

To define the composition of the biochemically isolated transcriptional condensates, we sequenced their DNA and RNA, alongside additional fractions for comparison (denoted in green, **Figure 1C**). Although each replicate’s fractions were derived from an independent preparation from cultured cells, corresponding fractions clustered together across replicates, indicating high reproducibility of fractionation (**Figure S2A-D**). As an initial test of whether POCEMON recovers genomic regions associated with transcriptional condensates, we assessed for transcriptionally active chromatin features. Genomic elements reproducibly enriched in Pol II or Med1 TCI DNA relative to the sonicated nuclei starting material were defined as respective TCI regions. Meta-analyses showed that both Pol II and Med1 TCI regions are strongly enriched for Pol II and Med1 ChIP-seq signal (**Figure 2A**), indicating that TCI-associated DNA identified biochemically corresponds to regions occupied by core transcriptional machinery. Although TCI regions comprise a subset of the most enriched Pol II-and Med1-bound loci, they also capture a set of distinct regions not encompassed by ChIP-seq peaks, particularly in Pol II TCI DNA (**Figure 2B**). These results argue that POCEMON does not simply recover global transcriptionally engaged chromatin but rather enriches a more restricted class of Pol II/Med1-occupied regions.

**Figure 2.**
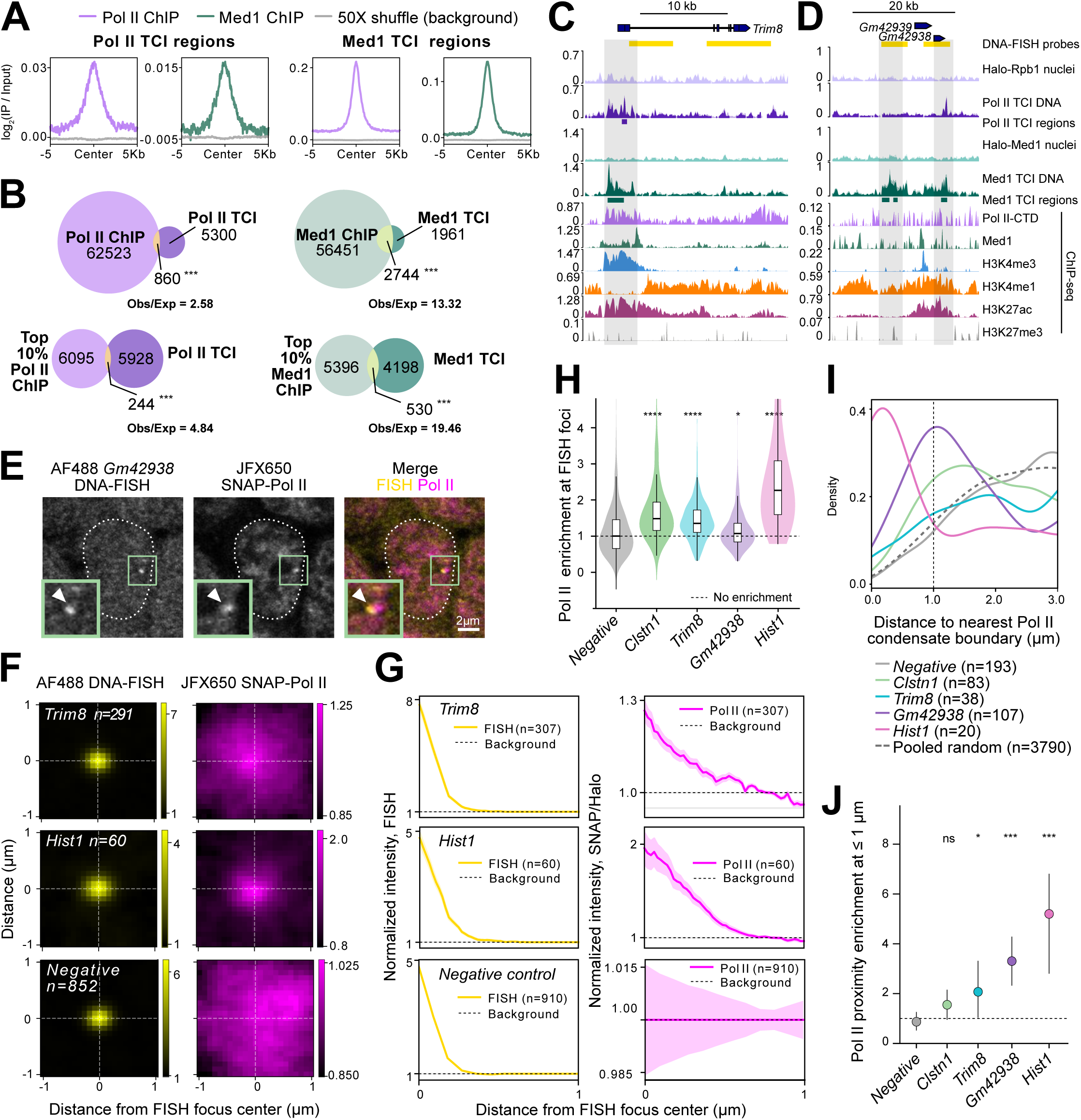
POCEMON identifies a subset of highly Pol II/Med1-occupied genomic regions that can localize near endogenous transcriptional condensates. **(A)** Metaplots showing mean Pol II and Med1 ChIP-seq log_2_(IP/Input) from Narita *et al.*^104^ contoured over Pol II and Med1 TCI DNA regions or a 50X-shuffled background. **(B)** Overlap of TCI DNA regions with Pol II- and Med1-bound genomic regions defined by called peaks from ChIP-seq. Top, all ChIP-seq peaks. Bottom, highest 10% ChIP-seq peaks, ranked by mean log_2_(IP/Input). **(C)** Representative Pol II TCI DNA region at the *Trim8* promoter. Tracks of input nuclear DNA, Pol II and Med1 TCI DNA, shown as replicate-median; and indicated ChIP-seq profiles from Narita *et al.*^104^ (Pol II, Med1, H3K4me1, H3K27ac), Kubo *et al.*^105^ (H3K4me3), and Illingworth *et al.*^106^ (H3K27me3). Yellow bars denote the DNA-FISH probe region. **(D)** Genome-browser track as in panel C, showing two long non-coding RNA loci, *Gm42938* and *Gm42939*. **(E)** Representative image of DNA-FISH at the lncRNA *Gm42938* locus together with tagged Pol II signal, showing an example of DNA focus–Pol II condensate colocalization. Insets show the enlarged boxed region. **(F)** Median intensity maps centered on DNA-FISH foci for the *Trim8*, the positive-control *Hist1*, and the negative-control locus, together with the corresponding Pol II signal. **(G)** Radial intensity quantification from panel F, showing enrichment of Pol II signal near the *Trim8* and *Hist1* DNA-FISH foci, and little enrichment near the negative-control locus. Median signal is shown along with standard error band. **(H)** Distributions of Pol II signal enrichment measured at the center of each FISH focus. \**p* < 0.05, \*\*\*\**p* < 0.0001 from Mann-Whitney U test versus the negative-control locus. **(I)** Smoothed density distributions of distances from FISH foci to the nearest segmented Pol II condensate boundary for all tested loci and the negative-control probe (<3 μm). The dashed curve indicates the pooled distribution of randomly sampled nuclear coordinates across all analyses. **(J)** The fraction of foci within 1 μm of a Pol II condensate boundary normalized to the same fraction for the randomly sampled nuclear coordinates. \**p* < 0.05, \*\*\**p* < 0.001 and 95% CI calculated using nucleus-wise bootstrap resampling.

We next sought direct cell-based validation that candidate TCI loci identified biochemically can localize at or near endogenous transcriptional condensates *in situ*. We selected representative loci with strong TCI DNA signal for DNA-FISH accompanied by imaging of tagged Pol II and/or Med1. Example tested TCI regions include *Trim8* promoter, a long non-coding RNA locus (**Figure 2C-D**), and enhancer-like elements (**Figure S2E**), all of which displayed robust active chromatin marks. Imaging of these loci revealed examples of spatial proximity or colocalization between the DNA-FISH focus and Pol II puncta (**Figure 2E, S2F**).

Average intensity maps centered on FISH foci showed proximal enrichment of Pol II signal (**Figure 2F, S2G**), whereas a negative-control locus (**Figure S2E**) did not show such enrichment (**Figure 2F**). By contrast, the positive-control HLB-associated gene *Hist1* (**Figure S2E**) displayed clear enrichment, as expected (**Figure 2F**). Radial intensity quantification confirmed these differences (**Figure 2G-H, S2H-I**). To quantify this spatial relationship in an object-based manner, we segmented Pol II or Med1 condensates and computed distance to nearest condensate boundary for each DNA-FISH probe. Within 1 μm, a relevant distance for condensate-controlled gene bursting,^23^ we observed greater proximity of the TCI loci (although one did not reach significance) to Pol II or Med1 condensates compared to the negative-control locus and randomly sampled nuclear coordinates (**Figure 2I-J**). Together, these imaging analyses provide orthogonal validation that loci identified by POCEMON can localize in close spatial proximity to endogenous Pol II- or Med1-marked transcriptional condensates and in Pol II-rich regions of the nucleus.

Notably, only ∼4% of cells showed Pol II–FISH or Med1–FISH colocalization for candidate condensate-associated regions. Given that POCEMON identifies thousands of Pol II-and Med1-TCI regions genome-wide (**Figure 2B**) and there are at most tens of large transcriptional condensates present per nucleus,^10^ this low per-locus colocalization frequency is not unexpected. Instead, this suggests that association with transcriptional condensates is heterogeneous across cells and/or dynamic, with only a small subset of candidate loci engaging transcriptional condensates at a given time in a single nucleus. Recovery of both enhancer- and promoter-proximal loci suggests that endogenous transcriptional condensates may engage multiple classes of regulatory DNA, providing a framework for dissecting which genomic elements preferentially associate with these structures.

### Transcriptional condensate-associated genomic regions are enriched at promoter-proximal regulatory elements

Although early models proposed condensate formation at super-enhancers and their target promoters,^7,8,10,26,28^ recent live-cell imaging has demonstrated a more dynamic view in which condensates transiently contact enhancer-promoter DNA.^23^ Conversely, the transcription factory literature proposes continued immobilization of a polymerase throughout the transcription cycle.^57^ Given these distinct predictions of prior studies, we envisioned that a genome-wide analysis of DNA elements which stably engage larger transcriptional condensates across a population of cells could potentially distinguish these models, afford insight into their function, and interrogate the generality of regulatory principles. If POCEMON is capturing biologically meaningful condensate-associated DNA rather than a nonspecific subset of active chromatin, then the recovered loci would be expected to display enrichment of particular candidate *cis*-regulatory elements (cCREs). To assess enrichment of specific classes of regulatory DNA, we compared the overlap of TCI regions with the ENCODE catalog of cCREs across annotation classes^58^ to a genome-shuffled permutation background. This analysis revealed selective enrichment of TCI regions (merged from the Pol II and Med1 pulldowns) at sequences with promoter-like and proximal enhancer-like signatures (**Figure 3A**). To our surprise, other cCRE classes, including distal enhancer-like sequences, were comparatively underrepresented in the combined TCI regions. Similar analyses hold for the Pol II and Med1 captured regions individually (**Figure S3A**), although Med1 accounts for more of the effect size.

**Figure 3.**
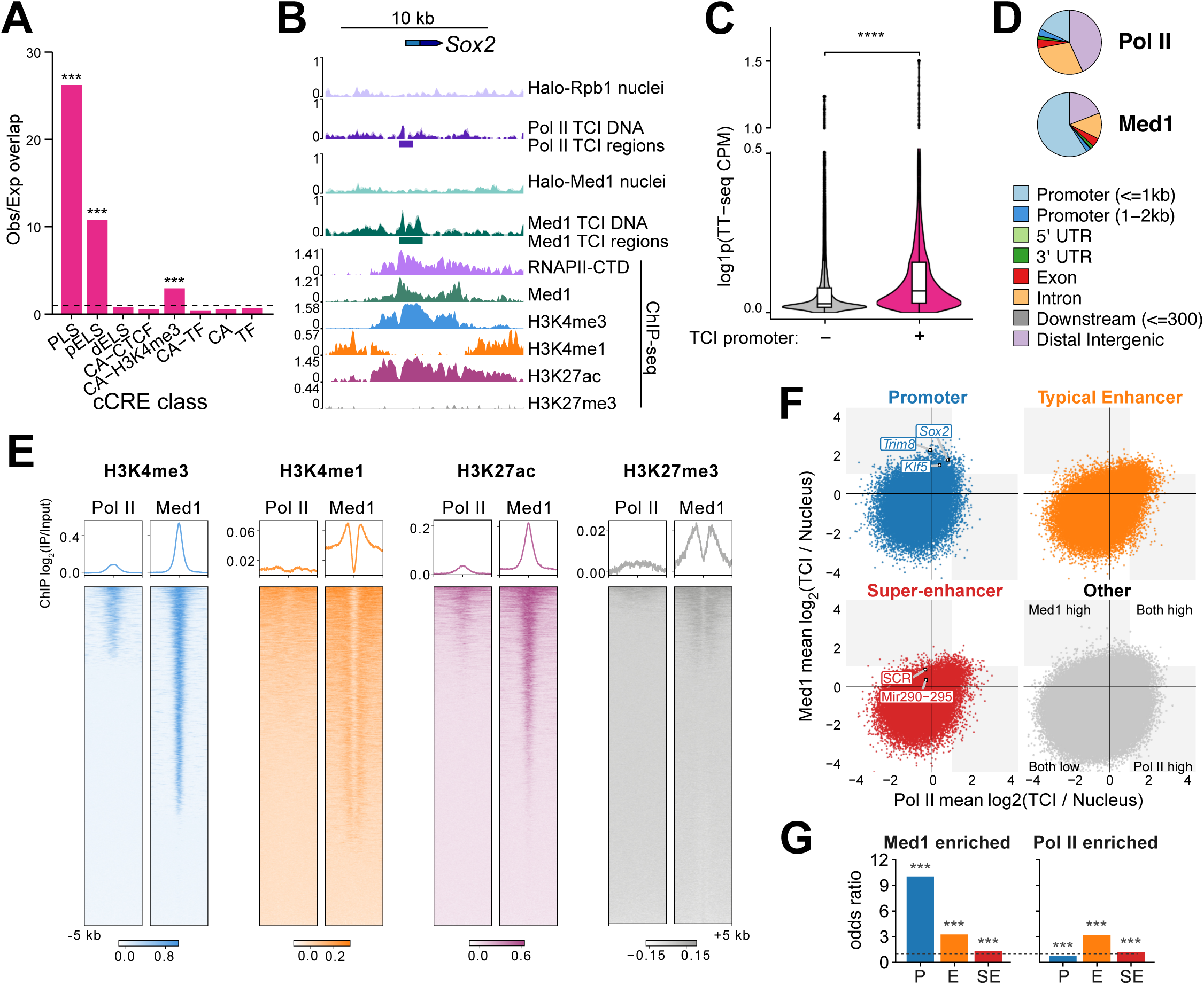
Transcriptional condensate isolates associate with promoters and promoter-proximal enhancer DNA. **(A)** Enrichment of Pol II and Med1 merged TCI DNA regions across ENCODE cCRE classes^58^. Bars show observed/expected overlap of the merged TCI DNA regions with: PLS – promoter-like signatures, pELS – proximal enhancer-like signatures (within 2 kb of TSS), dELS – distal enhancer-like signature, CA-CTCF – chromatin accessibility + CTCF, CA-H3K4me3 – chromatin accessibility + H3K4me3, CA-TF – chromatin accessibility + transcription factor, CA – chromatin accessibility, TF – transcription factor. \*\*\**p* < 0.001 based on permutation-derived empirical upper-tail *p*-values, Benjamini-Hochberg corrected. **(B)** Representative promoter-proximal TCI DNA region at the *Sox2* locus showing Pol II and Med1 TCI-associated DNA and ChIP-seq tracks as in Figure 2C.^104–106^ **(C)** Nascent transcription levels from Shao *et al.*^107^ integrated over gene bodies with non-TCI and TCI promoters, defined by promoter overlap with Pol II and Med1-merged TCI DNA regions. TT-seq signal was visualized as log1p-transformed mean CPM truncated at the 99.5th percentile of all data for clarity of display. \*\*\*\**p* < 0.0001 based on two-sided Wilcoxon rank-sum tests with Benjamini-Hochberg correction. **(D)** Genomic annotation of UCSC Known Gene elements^108,109^ for Med1 or Pol II TCI regions. **(E)** ChIP-seq metaprofiles and heatmaps of promoter- and enhancer-associated chromatin marks^104–106^ presented as log_2_(IP/Input) centered on Med1 or Pol II TCI regions, spanning 5 kb upstream and downstream. **(F)** Scatterplots of genome-wide 2-kb bins showing mean Pol II and Med1 TCI DNA enrichment relative to nuclear input, with bins stratified by overlap with promoters, typical enhancers, super-enhancers, or neither of those elements (other) from early mouse development.^68^ To highlight representative loci featured in other figures, labels were placed on the highest-signal overlapping 2-kb bin. **(G)** Odds-ratio analysis of region classes stratified by Med1-high or Pol II-high condensate enrichment. Med1 and Pol II enriched defined as bins where log_2_(TCI/nucleus) > 1. P – promoter, E – typical enhancer, SE – super-enhancer. \*\*\**p* < 0.001, computed using Fisher’s exact tests comparing bins with mean log2FC > 1 against all other bins.

We next examined a paradigmatic example of condensate-driven transcription in mESCs,^23^ the *Sox2* locus. The promoter-proximal active regulatory region displays robust TCI signal (**Figure 3B**), yet there is comparatively little TCI enrichment of DNA from the well-documented super-enhancer that neighbors it (**Figure S3B**). This locus exemplifies the broader genome-wide trend that condensate-associated regions frequently map to promoter-centered regulatory DNA rather than to gene-distal regulatory regions. Inspection of other known super-enhancer elements^59^ reveals similarly minimal enrichment (**Figure S3B**), consistent with the lack of distal-enhancer enrichment seen in the cCRE analysis. The genes whose promoters overlap merged TCI DNA elements are more highly transcribed than genes without TCI promoter peaks (**Figure 3C, S3C**), consistent with substantial prior evidence that larger transcriptional condensates promote transcription^7,10,21–24^. From this merged TCI analysis, it appears that condensate-associated DNA is not distributed uniformly across regulatory chromatin but is concentrated within promoters and proximal enhancer-like regions of actively transcribed genes, with unexpectedly limited capture of distal enhancer elements.

While this merged analysis of Pol II and Med1 can be justified by substantial apparent overlap of large condensates in higher resolution imaging,^10,23^ there are clear differences in the genomic elements captured by the two pulldowns. Foremost, in gene-centric categorization, there is a stronger promoter enrichment in Med1 TCI DNA and a bias of intronic and gene-distal regions in Pol II TCI DNA (**Figure 3D**). Nevertheless, there remains modest promoter enrichment when the size and number of elements are taken into account (**Figure S3A**). Further supporting our promoter-centric model, promoter Med1 TCI enrichment tracks with Med1 enrichment by ChIP at promoters (**Figure S3D**), consistent with the sufficiency of the Med1 IDR for large transcriptional condensate formation^7,17,34^.

As an orthogonal view of the apparent promoter-like features of Med1 TCI regions, we analyzed ChIP-seq for histone marks associated with active promoters and enhancers. H3K4me3, an exclusive mark of active promoters^60–62^, and H3K27ac, found at both active promoters and enhancers^63–65^, display enrichment at TCI region centers with markedly higher signal for Med1 captured material (**Figure 3E**). Interestingly, H3K4me1, which is also present at active promoters and enhancers^62,64,65^ exhibits a distinct, center-depleted pattern particularly at Med1-TCI regions with length-scales more consistent with wider promoter distributions that flank the higher methylforms^62^. The H3K27me3 mark of facultative heterochromatin^66,67^, displayed very low overall signal at Med1 and Pol II regions, arguing against substantial Polycomb-chromatin association^65^. These results indicate that within TCIs, Med1 is more strongly associated with promoter-like chromatin than Pol II. Intriguingly, upon transcription inhibition by TPL treatment, we uncover a *de novo* association of promoters with TCIs (**Figure S3E**), and accompanying increase in active chromatin marks (**Figure S3F**). Under identical TPL treatment conditions, the accompanying study (Pandey *et al.*) also notes dramatic increases in the same marks within condensates using fixed-cell microscopy. Moreover, they observe an overall increase in histone density, indicating that under initiation inhibition and Pol II degradation, condensates remodel to include more active chromatin, potentially accommodating for the Pol II loss.

To directly compare the capture for Med1 to Pol II genome-wide, without the potential bias of peak calling and to explicitly delineate enhancers and super-enhancers within intergenic elements, we analyzed condensate enrichment as log_2_(TCI/nucleus) across 2-kb bins, stratifying bins by overlap with known promoter, enhancer, or super-enhancer elements (**Figure 3F**).^68^ Promoter bins were ∼10-fold enriched in Med1-high bins relative to all bins, but slightly underrepresented in the independently defined Pol II-high set, reinforcing the Med1 TCI promoter bias interpretation (**Figure 3G**). Early developmental enhancers showed moderate enrichment in both groups (odds ratio ∼3), indicating comparable enhancer representation in Pol II- and Med1-enriched bins, with no evidence for a Pol II-specific enhancer bias. Given the prior connections of large condensates to super-enhancer driven transcription,^7,10,22,23,26,28^ it is surprising that early developmental super-enhancer elements displayed weak, but nevertheless statistically significant enrichment in both groups (**Figure 3F-G**). Together, these data indicate that Med1 occupancy within transcriptional condensates is highest at Med1-rich promoters, while enhancer enrichment is moderate and comparable between Pol II and Med1, yet the divergence between these two POCEMON captures remained puzzling.

### Transcriptional condensate isolates reveal distinct Med1 and Pol II occupancy suggesting a role in promoter escape and promoter-proximal pause release

To examine how the depletion of Pol II at promoters manifested at the gene structure level, we compared gene profiles of TCI DNA to corresponding Pol II and Med1 ChIP-seq occupancy. Med1 occupancy within TCIs closely resembled Med1 ChIP, whereas Pol II TCI differed strikingly from bulk chromatin-engaged Pol II (**Figure 4A, S4A**). Remarkably, TCI Pol II displays pronounced depletion at the transcription start site (TSS) and relative enrichment on both sides of it, quite distinct from Pol II ChIP, which displays the canonical promoter-proximal pause-centered peak (**Figure 4B**).^61,69,70^ As the Pol II TCI DNA signal peaks downstream of the mean pause site, we interpret this pattern to reflect both elevated promoter clearance and increased pause-release^70–72^ within transcriptional condensates. In support, mean Pol II and Med1 TCI RNA both exhibit a subtle shift downstream into the gene body relative to nascent sequencing (**Figure 4B, S4B**), which could be interpreted as representing a pool of transcripts emanating from polymerase released from promoter-proximal pausing.

**Figure 4.**
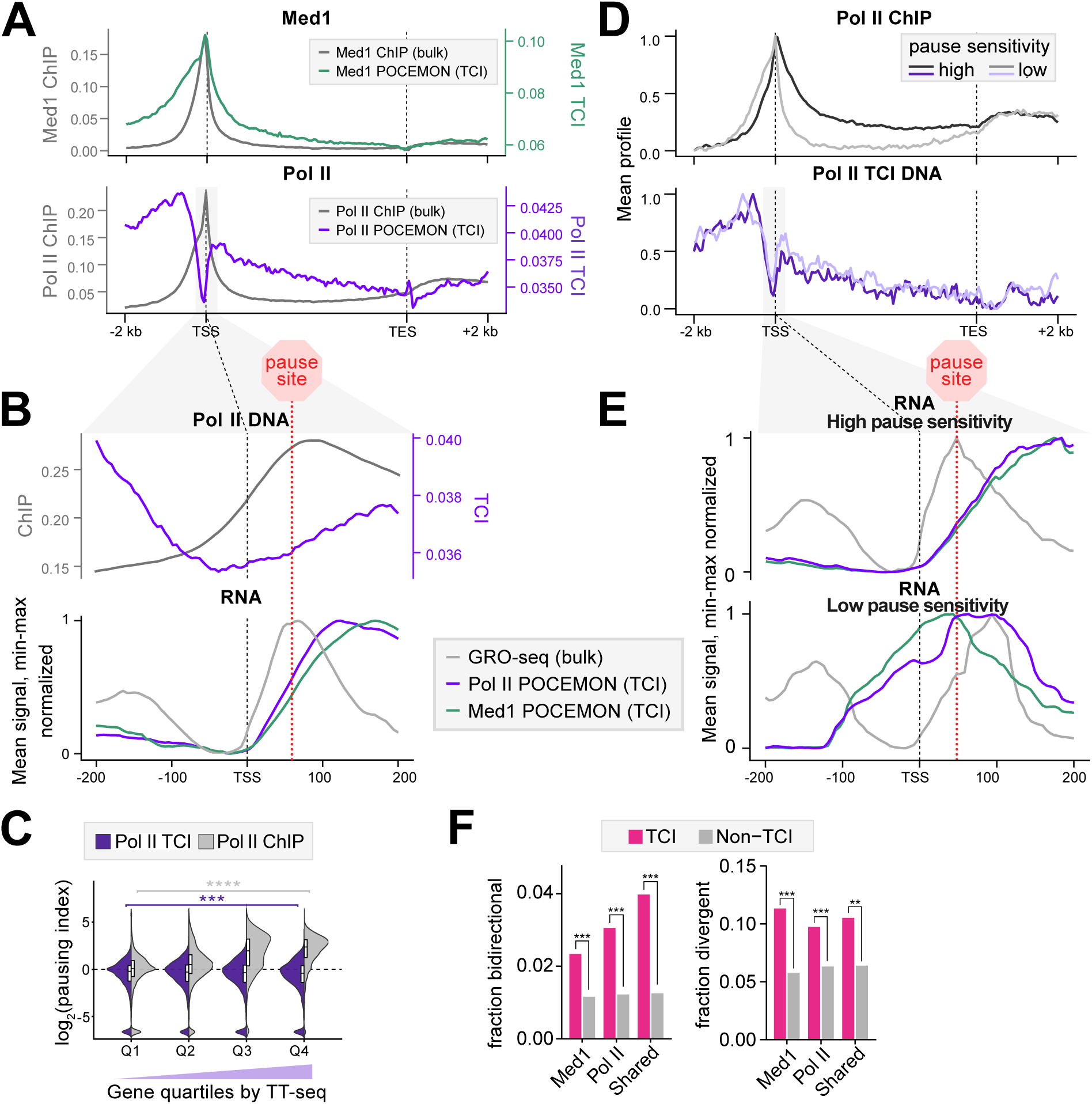
Transcriptional condensate isolates capture promoter-centered Med1 and pause-released Pol II. (**A**-**B**) Metagene profiles across all genes longer than 1 kb. Bulk Pol II and Med1 ChIP-seq are shown as log_2_(IP/input),^104^ and TCI DNA is shown as mean CPM-normalized signal. **(B)** Zoomed meta profiles across 400 bp windows centered on TSSes, shown for Pol II ChIP,^104^ Pol II TCI DNA, nascent RNA (GRO-seq),^77^ and Pol II and Med1 TCI RNA. The approximate pause site was manually annotated based on the GRO-seq promoter-proximal signal. **(C)** Violin plots of pausing index stratified by promoter quartiles ranked by nascent transcription levels.^107^ \*\*\**p* < 0.001, \*\*\*\**p* < 0.0001 from two-sided Wilcoxon rank-sum test with Benjamini-Hochberg adjustment; log_2_-transformed pausing indices were winsorized at the 4th and 99.9th percentiles for clear visualization of the bulk of the distribution. **(D)** Metagene profiles as in panel (A), shown for genes in the top and bottom 10% of pause sensitivity to 5 min flavopiridol (FP) treatment, calculated based on GRO-seq data from Jonkers *et al.*^77^ as log_2_(FP pausing index / untreated pausing index). Each track was min-max normalized. **(E)** Zoomed meta profiles across 400 bp windows centered on TSSes, shown for nascent RNA (GRO-seq^77^), and Pol II and Med1 TCI RNA. Gene pause sensitivity groups are shown as in panel (D). Pause site is manually annotated as in panel (B). **(F)** Fraction of promoter regions classified as bidirectional, defined as head-to-head promoters within 1 kb,^110^ or divergent, defined from mESC FANTOM5 CAGE signal^111,112^ as opposite-strand TSSes within 300 bp for TCI-associated and non-TCI promoter loci. Bars show the fraction of promoters in each category (Med1, Pol II, shared) classified as bidirectional or divergent. \*\*\**p* < 0.001 Fisher’s exact test with Benjamini-Hochberg-adjustment.

To more directly assess pausing, we computed the pausing index for bulk Pol II^69,73^ and a similar metric for TCI Pol II, then plotted both across gene quartiles ranked by nascent transcription (**Figure 4C**). Whereas the bulk Pol II ChIP pausing index increased with transcriptional output, the Pol II TCI-derived pausing index remained comparatively stable and was modestly reduced in the most highly transcribed genes. This argues that even genes that normally undergo substantial promoter-proximal pausing may be more rapidly released into elongation when associated with large transcriptional condensates. This observation is consistent with the recent detection of P-TEFb, the complex responsible for pause release, within Brd4/Med1 transcriptional condensates by *in situ* proteomics,^20^ and coacervation of large Med1 condensates with Spt6 and the Paf complex,^17^ both of which can only productively engage Pol II upon pause release.^71,74–76^ To test this idea, we contoured Pol II TCI DNA over the top and bottom decile of genes ordered by acute sensitivity to flavopiridol.^77^ These genes represent the limiting cases of genes that are very sensitive to P-TEFb-mediated pause release or largely independent (**Figure 4D, S4C-D**). The mean bulk Pol II ChIP peak shifts from TSS-centered distribution in the low pause sensitivity set to a position deeper into the gene body in the high pause sensitivity gene set, reflecting promoter-proximal pausing ∼40 bp downstream from the TSS.^69,70,72,77^ In contrast, TCI-associated Pol II DNA showed little detectable difference between these two gene sets. That the most and least pause-prone genes display similar profiles in TCI Pol II capture, further supports the model whereby genes in transcriptional condensates escape the promoter-proximal pause into productive elongation more readily.

Furthermore, the RNA recovered from Pol II and Med1 TCIs for the least pause sensitive genes is similar to the bulk nascent RNA profile at the promoter, whereas the most pause sensitive promoters display an RNA distribution that peaks past the pause site (**Figure 4E, S4B-E**), recapitulating the global pattern observed (**Figure 4B**). Collectively, these analyses are consistent with condensates accelerating promoter escape and early pause release relative to the same events in the bulk population.

The substantial enrichment of condensate-associated Pol II upstream of TSSs prompted us to test whether TCI regions preferentially associate with bidirectional and/or divergent promoters. We found that condensate-associated promoter loci were indeed enriched for structurally bidirectional promoters and divergent transcription (**Figure 4F and S4F-G**).^70,77–79^ While bidirectional and divergent promoters comprise only a minority of all and TCI-associated promoters (**Figure S4F-G**), both classes were significantly enriched (**Figure 4F**). Pol II-Med1 TCI co-occupancy was highest for bidirectional promoters (**Figure S4G**). These results indicate that TCI promoter regions are preferentially associated with promoter architectures that support opposing-direction transcription initiation.

### Transcriptional condensates host pervasive nascent transcription and are shaped by transcriptional activity

Having defined the DNA composition of TCIs, we next performed a more thorough analysis of the captured RNA. Both Med1 and Pol II TCI transcriptomes were largely genic in origin with high intron content, more closely resembling the nascent transcriptome than exon-rich mature nuclear RNA (**Figure 5A**). Despite less RNA recovered from Pol II TCIs treated with triptolide, the remaining RNA displayed higher exon and lower intron composition, consistent with changes in the TCIs reflecting a global shift to more mature RNA profiles upon transcriptional inhibition and the RNA remaining associated with the transcriptional condensates during processing (**Figure 5A**). In support, we observe TCI capture across the whole gene body (**Figure 5B, S5A**), and apart from the promoter-proximal pause differences (**Figures 4B,E**), this is in rough proportion to its nascent abundance. While we interpret this nascent RNA association as continued adhesion to its site of production, it could alternatively represent adventitious binding of local RNA unrelated to condensate biogenesis. Interestingly, extract-reconstituted transcriptionally active condensates also harbor the transcripts they produce, and do not substantially bind bulk RNA.^34^

**Figure 5.**
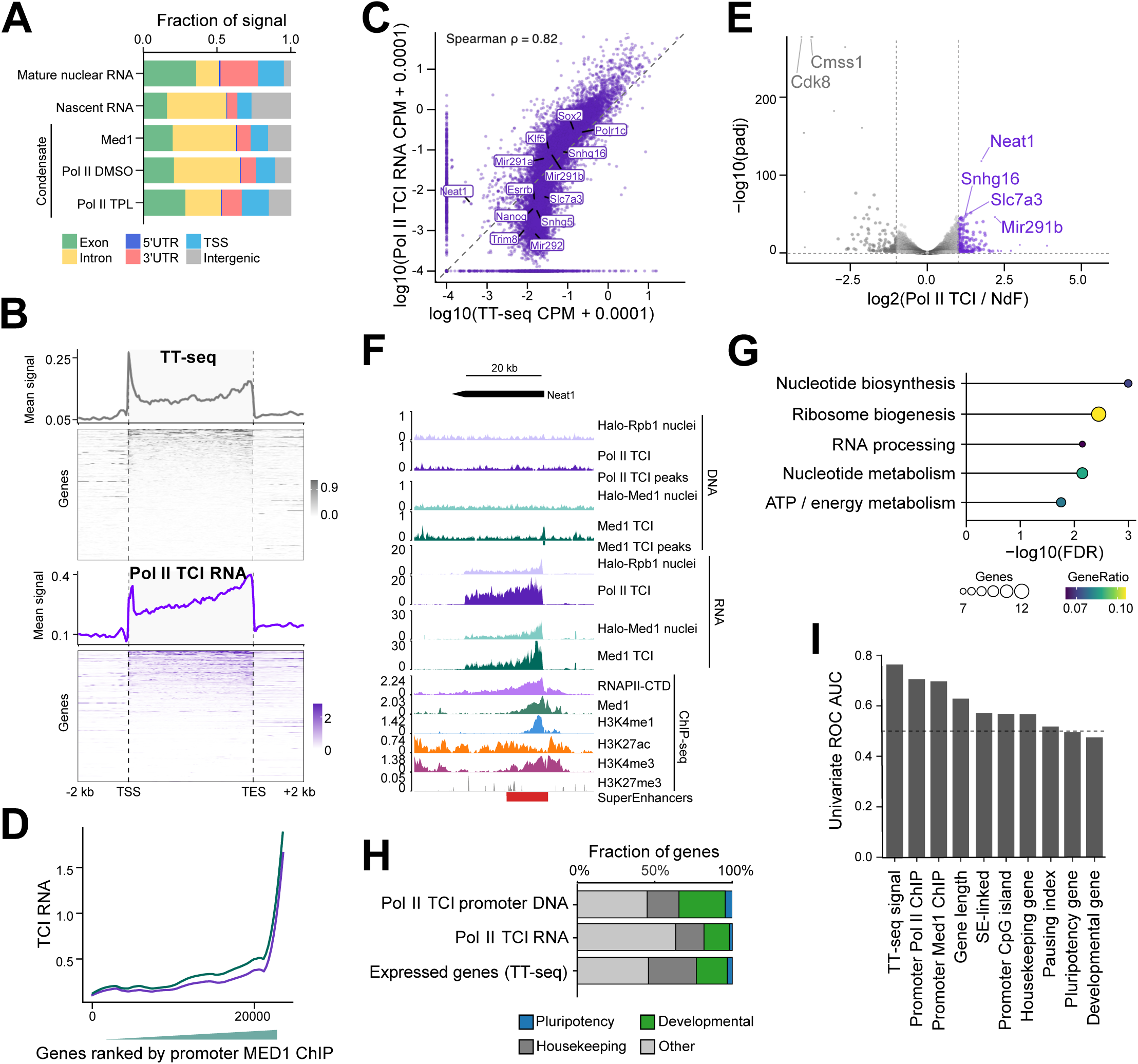
Transcriptional condensates host pervasive nascent transcription. **(A)** Distribution of signal across genomic features in mature nuclear RNA from a polyA selection,^113^ nascent RNA from TT-seq,^107^ Med1 TCI RNA, and Pol II TCI RNA under DMSO and TPL conditions. **(B)** Metaprofile and heatmap of CPM-normalized nascent RNA from TT-seq^107^ and Pol II TCI RNA from the DMSO condition. Genes in heatmap were arranged by decreasing gene body TT-seq signal. **(C)** Scatterplot comparing mean Pol II TCI RNA and TT-seq^107^ signals across genes, with selected representative loci labeled. Signals were plotted as log_10_(CPM + 0.0001). Axes were truncated from −4 to the larger of the 99.95th-percentile x- or y-axis values for visualization only; Spearman correlation was calculated using untransformed gene-level mean signal values. **(D)** Pol II and Med1 TCI RNA levels across genes ranked by promoter Med1 ChIP signal. The line represents a LOESS-smoothed fit. **(E)** Differential expression analysis of Pol II TCI RNA relative to NdF, showing genes enriched or depleted in TCIs. Genes are plotted by apeglm-shrunken log_2_ fold change and −log_10_ adjusted p-value. Significant enriched or depleted genes were defined as adjusted p-value < 0.05 and absolute shrunken log2 fold change ≥ 1; selected genes are labeled. **(F)** Genome-browser views at the *Neat1* locus comparing nuclear input, Pol II TCI DNA, Pol II TCI peaks, Med1 TCI DNA, Med1 TCI peaks, corresponding TCI RNA tracks, ChIP-seq as in Figure 2C,^104–106^ and super-enhancers.^68^ **(G)** Gene Ontology analysis of genes enriched in Pol II TCI RNA relative to the NdF RNA. Significant GO terms were manually collapsed into broad functional categories by grouping related terms and taking the union of genes within each category. Point size indicates the number of genes in each grouped category, and color indicates the gene ratio. **(H)** Fraction of genes classified as pluripotency, developmental, housekeeping, or other among mESC expressed genes derived from TT-seq,^107^ genes enriched in Pol II TCI RNA relative to the NdF RNA, or in genes whose promoter overlapped Pol II TCI DNA regions. **(I)** Univariate receiver operating characteristic (ROC) analysis evaluating predictors of Pol II condensate association. The dashed line denotes an AUC of 0.5, corresponding to random classification performance.

Although earlier TCI DNA element analysis suggested that TCI promoters represent more actively transcribed genes, we sought to directly measure this effect from the captured RNA. Congruent with the prior analyses **(Figures 3C, S3C**), gene-level TCI RNA signal was highly correlated with corresponding nascent RNA levels (**Figure 5C, S5B**). We note that low expression transcripts were underrepresented in the condensate. Similar comparisons of TCI gene body RNA versus TCI promoter DNA (**Figure S5C**) lend further support for a model in which condensates associate with active promoter-centered chromatin and with nascent RNA they produce. Both Pol II and Med1 TCI RNA abundance displayed progressive increase across genes ranked by promoter Med1-ChIP occupancy, with the most drastic increase observed among promoters with the highest Med occupancy (**Figure 5D**). Collectively, these data further support the inference that TCIs do not represent a privileged class of transcriptional events.

As an additional test for preferential condensate association of transcripts, we performed differential expression analysis between TCI and nuclear or NdF RNA. This analysis identified a subset of RNAs modestly enriched in TCIs (**Figure 5E**), including microRNAs from the pluripotency-linked miR-290-295 cluster, which lies adjacent to a potent mESC super-enhancer,^59^ and several histone cluster genes, consistent with enrichment of HLBs (**Figure 5E, S5D).** Inspection of representative enriched loci showed concordant enrichment of condensate-associated RNA and Med1 promoter DNA within active chromatin landscapes (**Figure 5F**).

Genes enriched within the condensate transcriptome did not appear restricted to canonical lineage-specific or super-enhancer regulated genes. Instead, gene ontology analysis of Pol II TCI-enriched RNAs revealed overrepresentation of genes involved in housekeeping functions, not typically regulated by distal enhancers,^29^ including nucleotide biosynthesis, ribosome biogenesis, RNA processing, and energy metabolism (**Figure 5G).** Comparison of broad gene classes revealed that condensate-enriched genes correspond well to those of all expressed genes (**Figure 5H, S5E-F**). These results argue that, rather than being limited to a small set of lineage-specific loci, transcriptional condensates in mESCs broadly engage active genes.

To better understand which factors predict gene localization to transcriptional condensates, we used a univariate AUC model and ranked individual gene features by their ability to distinguish condensate-associated transcripts (**Figure 5I, S5G**). TT-seq gene-body signal and promoter Pol II and Med1 occupancy were the strongest predictors. In contrast, developmental and pluripotency gene annotations were weak discriminators, indicating that condensate association is not primarily explained by canonical cell-identity gene classes. Other features, including gene length, super-enhancer linkage, CpG island linkage, housekeeping status, pausing index, and gene-body Pol II condensate DNA levels, showed intermediate predictive value.

### Transcriptional condensate isolate elements occupy highly interconnected 3D chromatin neighborhoods

We next examined whether condensate-associated genomic regions also exhibit distinctive 3D interaction properties, as might be anticipated for stereotyped hubs of gene expression that are large enough to accommodate many promoter-like elements. To query 3D genome organization, we compared TCI signal to region capture Micro-C across two 1-Mb loci encompassing *Klf1* and *Ppm1g* (**Figure 6A, S6A**).^80^ The *Klf1* gene-rich locus contained several Pol II and numerous Med1 TCI regions, many of which seemed to correspond to highly H3K4me3 decorated active promoters. Node graphs of these two loci revealed a structured local interaction network in which promoters overlapped loop anchors more frequently than enhancers (**Figure 6B-C** top panels, and **S6B**). Loop anchors overlapping merged Pol II and Med1 TCI regions occupied more highly connected positions within this network than non-TCI anchors, exhibiting higher interaction degree overall (**Figure 6B-C**, bottom panels).

**Figure 6.**
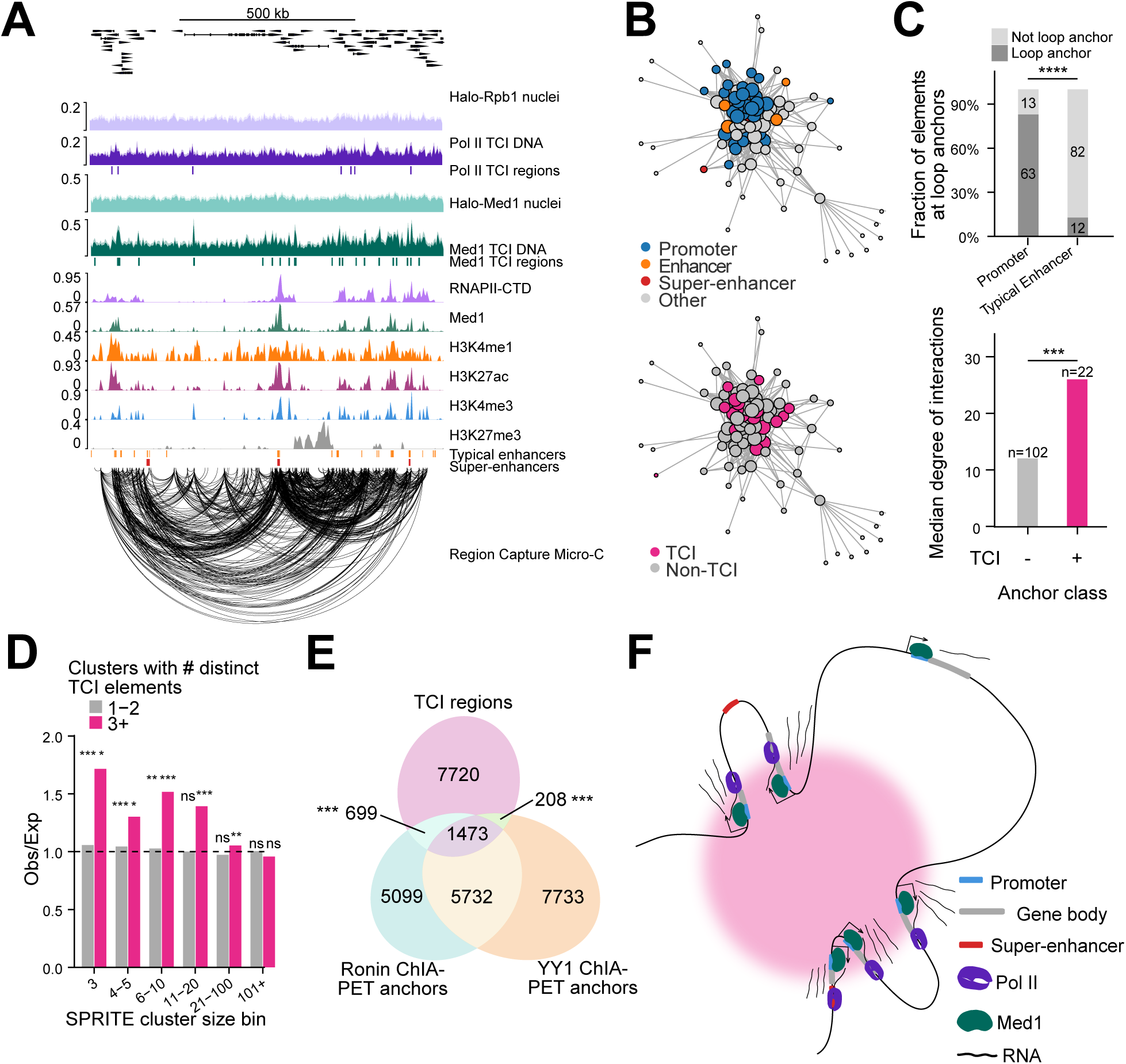
Transcriptional condensate isolates capture promoter-rich local chromatin networks. **(A)** Genome browser track view across a 1 Mb locus centered on *Klf1* showing Pol II and Med1 TCI DNA, corresponding TCI regions, ChIP-seq tracks,^104^ enhancer annotations,^68^ and region capture micro-C loops from Goel *et al.*^80^ **(B)** Network representation of loop anchors from the captured locus, representing loop anchors as nodes, with node size proportional to interaction degree, and looping interactions as edges. Top, anchors colored by annotation class (promoter, enhancer, super-enhancer,^68^ or other). Bottom, anchors colored by overlap with merged TCI regions. **(C)** Quantification of anchor properties within the capture Micro-C networks from panel B and Figure S6B. Top, fraction of promoter and typical early mouse enhancer elements^68^ that overlap loop anchors. Bottom, median degree of interaction for TCI and non-TCI anchors. \*\*\**p* < 0.001, \*\*\*\**p* < 0.0001 from Fisher’s exact test (top) and Wilcoxon rank-sum test (bottom). **(D)** Enrichment of TCI DNA elements in multiway SPRITE clusters from Quinodoz *et al*.^81^ SPRITE clusters were binned by cluster size, and clusters containing either 1-2 or ≥3 distinct TCI DNA elements are shown. TCI DNA elements were defined from merged Pol II and Med1 TCI DNA peaks. Bars show observed/expected enrichment relative to 5,000 label-shuffle permutations. The dashed line indicates observed/expected = 1. \**p* < 0.05, \*\**p* < 0.01, \*\*\**p* < 0.001 from empirical upper-tail permutation p-values with Benjamini-Hochberg correction. **(E)** Euler diagram showing overlap between the merged TCI DNA peak set and Ronin and YY1 ChIA-PET anchors. Observed overlap between TCI peaks and Ronin^29^ or YY1 loop anchors^82^ was compared to a ENCODE^58^ class-, chromosome-, and width-matched regulatory-element null. Significance was assessed by 5,000 permutations. \*\*\**p* < 0.001 based on permutation-derived empirical upper-tail *p*-values with Benjamini-Hochberg correction. **(F)** Model summarizing the promoter-centered and locally interconnected chromatin organization of transcriptional condensates.

To address the possibility that multiple promoters might colocalize within one condensate, we analyzed multiway, single-particle chromatin interaction data generated by SPRITE.^81^ To our surprise, TCI DNA regions were enriched above expectation within smaller SPRITE clusters, particularly clusters containing up to ∼ 20 elements (**Figure 6D, S6C-D**) thought to preferentially capture local chromatin contacts.^81^ This suggests that condensate-associated DNA preferentially participates in compact, local interaction neighborhoods, rather than in large-scale nuclear assemblies. Moreover, this enrichment was driven specifically by clusters containing multiple distinct TCI elements, indicating that condensate-associated loci are not merely present in pairwise interactions, but can co-occur within multiway local interaction structures.

Because POCEMON preferentially enriches 3D-connected promoters and proximal enhancers, we tested whether TCI regions are likewise associated with anchors of architectural-factor-mediated loops using ChIA-PET datasets for Ronin and YY1.^29,82^ We assessed TCI overlap with Ronin- or YY1 loop anchors against cCRE class-matched background elements, to control for the regulatory composition of TCI regions. We found significant enrichment for anchors from both factors, with a stronger overlap observed for Ronin (**Figure 6E**), consistent with the promoter-centered character of the TCI landscape, since Ronin bridges promoter-promoter interactions, while YY1 bridges enhancer-promoter and enhancer-enhancer interactions.^29,82^ Notably, Ronin regulates metabolic and housekeeping genes, which are well-represented within TCI-enriched gene sets (**Figure 5G**). The significant overlap with both factors suggests that condensate-associated regions participate in promoter-promoter and enhancer-promoter looping.

Taken together, these analyses suggest that the genomic DNA recovered by POCEMON is not an arbitrary subset of accessible or transcriptionally active chromatin. Instead, these data support a model in which transcriptional condensates embed promoter-proximal regulatory elements and are preferentially localized within chromatin neighborhoods of high interactivity (**Figure 6F**).

## DISCUSSION

We developed POCEMON, a biochemical approach which enables access to larger transcriptional condensates and affords a systematic view of their nucleic acid components. Many lines of evidence support the idea that POCEMON preserves the structure of transcriptional condensates in living cells. Sequencing of TCI-associated nucleic acids reveals unexpectedly broad heterogeneity of genic composition, with thousands of gene promoters and nascent transcripts embedded within across the population. Mediator and RNA polymerase II occupy distinct regions within TCIs, suggesting that condensates might enable efficient entry into transcription elongation. Together, our findings establish a promoter-centric model of transcriptional condensate organization at steady-state and suggest that condensate associations are widespread across active genes.

### Biochemical isolation appears to preserve endogenous condensate structure

We reasoned that the integrity of large transcriptional condensates outside the nuclear milieu is plausible because they are unusually stable^10,35^ and may become even less dynamic under the limited NTP conditions during the preparation. Cell rupture and nuclear isolation in large volumes relative to intact cellular context drives NTP nuclear egress, with further orders of magnitude dilution upon sonication and sedimentation gradient stage. Thus, effective NTP depletion during this preparation as well as the lowering of temperature may inhibit transcriptional condensate dynamics or disassembly. Based on these dilution estimates, the NTP concentration available by the time of sonication is expected to fall well below the reported *K_m_* for the yeast Pol II.^83^ Beyond limiting the activity of the transcriptional apparatus, ATP depletion can directly lead to solidification of biomolecular condensates^84–86^ or indirectly by limiting ATP-dependent chromatin remodeling enzymes,^87^ and other biomolecular condensates such as stress granules require ATP for their disassembly.^88^

Multiple orthogonal lines of evidence show that TCIs are representative of endogenous transcriptional condensates. First, confocal imaging of Pol II shows that punctate structures are retained throughout the POCEMON protocol (**Figure 1D**). Second, isolated material displays molecular identity corresponding to transcriptional condensates. Pol II, particularly its initiating phosphoisoform, is progressively enriched throughout the protocol, while markers of other nuclear bodies are depleted during the preparation, with residua notably remaining in the supernatant upon streptavidin pulldown from the F10 (**Figure 1E-F, S1E,G**). Third, cryoelectron tomography of F10 reveals discrete mesoscale assemblies whose sizes roughly correspond to large transcriptional condensates in cells,^10,23^ and are reminiscent of recently characterized nucleosome array condensates/mesoscale chromatin domains.^89,90^ Fourth, the density of transcriptional condensate isolates is sensitive to *in cellulo* transcriptional inhibition and Pol II degradation via triptolide,^55,56^ or to 1,6-hexanediol relative to 2,5-hexanediol treatment, a widely used metric for biomolecular condensates (**Figure 1I-J, S1A-B**).^7,9,10,53,54^ Fifth, TCIs are associated with a restricted subset of transcriptionally active chromatin, distinct from bulk nuclear Med1 and Pol II regions (**Figure 2A, B**), arguing against enrichment of a nonspecific set of Pol II and Med1 occupied regions. Moreover, this chromatin subset is not uniformly distributed across regulatory DNA (**Figure 3A,F,G**) but specifically enriched for 3D-interconnected promoters (**Figure 6**), in line with expectation for higher order chromatin hubs.

Sixth, TCI regions colocalize with endogenous transcriptional condensates more frequently and are generally more condensate-proximal in cells than negative controls (**Figure 2I-J**), despite the low probability expectation of such an association occurring in any given cell inferred from their cell-to-cell compositional heterogeneity. Collectively, these results demonstrate that POCEMON successfully enriches for transcriptional condensates while preserving their integrity by all available measures.

### The genic heterogeneity and permissive nature of transcriptional condensates

Initial studies postulated that transcriptional condensates preferentially regulate lineage-specific genes;^7,8,22,23,25–28^ a later study extended condensate regulation to housekeeping genes.^29^ Our data point to an even broader view, in agreement with early work which found pervasive transcription within transcription factories.^30^ Though transcription factories are more analogous to short-lived, smaller transcriptional condensates observed in living cells in both their dimensions and number per nuclei,^10,57^ this apparent similarity raises the question of whether there is a functional distinction beyond the apparent dynamics and size. More direct comparisons are needed to address this point. TCI nucleic acids show substantial genic diversity, with no strong bias to any particular gene class (**Figure 5G, H, S5F**). TCIs contain thousands of DNA elements genome-wide (**Figure 2B**), which are more condensate-proximal compared to negative controls (**Figure 2I-J**) but colocalize with condensates relatively rarely (∼4%). This low colocalization frequency agrees with previous work showing that condensate-gene associations are dynamic and transient.^23^ It is also consistent with the large numbers of condensate-associated regions identified by POCEMON, since only a small subset could plausibly concurrently occupy the limited volume of large condensates in the nucleus, so colocalization of any given element with a condensate is therefore rarely captured. Consistent with the broad condensate-promoter engagement over a population of cells, TCI RNA for both Pol II and Med1 preparations are highly correlated to the nascent transcriptome (**Figure 5C, S5B**). Together, these data argue against privileged regulation of a small set of target genes by large transcriptional condensates and instead support a model in which transcriptional condensates globally engage active genes.

### Promoter-centric organization of transcriptional condensates

Transcriptional condensates are thought to form on and associate with super-enhancer DNA elements.^7,8,22,23,25–28^ We find that promoters are highly enriched within TCIs, while distal enhancers and super-enhancers surprisingly lack enrichment (**Figure 3A, S3A**). This was unexpected as Mediator enrichment is how super-enhancers have previously been defined,^59^ and we note a strong correlation between Med1 ChIP occupancy and TCI enrichment. A plausible explanation for the promoter-super-enhancer disparity in TCI representation is a core-shell organization of these elements within transcriptional condensates, where promoters are integrated into the core, and super-enhancers positioned peripherally. Evidence supporting this hypothesis is presented in the accompanying manuscript, Pandey et al. Given a peripheral position of super-enhancers within condensates, the short DNA fragment size generated by sonication (∼1500 bp, **Figure S1D**) could lead to preferential loss of super-enhancer DNA during the POCEMON preparation. Alternatively, super-enhancers might play a role in condensate nucleation, which POCEMON may not capture if these early assemblies are smaller, less dense, or very short lived.

In accordance with our findings, previous studies have pointed at promoter-centric 3D chromatin organization. Pol II has been shown to spatially bridge promoters of active genes.^91^ Ronin-bridged promoter clusters of housekeeping genes, generally lacking distal enhancer elements, have been demonstrated to form transcriptional condensates.^29^ More recently, Mediator has been found to be dispensable for enhancer-promoter contacts, but important for fine-scale promoter structure,^92,93^ in accord with structural analysis of the PIC which support a role for Mediator promoting Cdk7-Ser5p activity and staging promoter firing.^94–96^ In yeast, where regulation is dominated by promoter-proximal upstream regulatory sequences in the absence of distal enhancers,^97^ Mediator plays a central coactivator role.^98^ This view differs from Mediator models in which Mediator functions as a through-space bridging factor to distal enhancers,^99,100^ although it does not preclude a role for transcriptional condensates in bridging promoters through space (**Figure 6**). Given the extensive Mediator decoration of super-enhancers by ChIP,^59^ formaldehyde crosslinking of these elements to the TCI periphery may best explain this disconnect. In agreement with a central role for promoter DNA in condensate formation, active transcriptional condensates of similar size were shown to self-assemble on promoter DNA in a cell-free system.^34^ Our findings corroborate this promoter-centricity and suggest that Mediator or Pol II recruits gene promoters to condensates.

### Insights into transcriptional condensate function

A number of studies found a causal connection between condensates and transcription or transcriptional bursting.^4,21,23,24,32–34,101^ Precisely how condensates regulate transcription remained unclear. We find that Mediator and Pol II occupancy within TCIs is consistent with a model in which condensates facilitate rapid entry into elongation. Unlike Mediator, which is highly enriched at promoters, TCI Pol II is strikingly depleted from a region that spans from just upstream of the TSS past the mean promoter-proximal pause site,^69,71,72,77,102^ but does occupy flanking regions (**Figure 4A-B**). The absence of capture from this region raises the possibility that TCIs may promote both PIC firing and release from promoter proximal pausing into early elongation. Although transcription from the handful of large condensates per nucleus is heterogeneous and rare in the population of cells, we propose such events appear to drive much more rapid initiation and promoter-proximal pause release than transcription of the same gene outside of the condensate. This model is in line with increased gene bursting as a function of its proximity to a transcriptional condensate.^23^

Since TCIs contain many promoters, it is plausible that they coordinate transcription across gene loci, as has been suggested before for Pol II bridged promoters.^91^ Utilizing SPRITE multiway interaction data^81^ which map higher order chromatin interactions from single cells, we identify concurrent condensate-promoter interactions (**Figure S6D**). We find that multiple TCI promoters colocalize in single SPRITE clusters, but this co-localization over background tapers off for clusters much larger than 20-DNA elements. In the accompanying work, we find a similar number of promoter-like nucleosome nanodomains within the larger Pol II clusters by super-resolution microscopy (Pandey *et al.*). These observations imply that transcriptional condensates are not merely concentrating transcriptional machinery, but potentially also coordinating transcriptional bursting from multiple genes which simultaneously contact the condensate, perhaps accounting for the orchestrated bursting of multiple distal genes which are spatially juxtaposed.^103^

Together, our findings support a model in which large transcriptional condensates are promoter-centered assemblages which engage a broad set of genes in proportion to their bulk transcription, rather than a stereotyped gene class. We propose that these nuclear bodies might function as sites that couple efficient promoter loading of the transcriptional machinery, Pol II promoter escape, and release from promoter proximal pausing, by concentrating the factors regulating these transcriptional steps. By providing biochemical access to larger transcriptional condensates, POCEMON opens a path toward defining how condensate architecture, promoter organization, and nascent transcription are coupled in endogenous nuclear environments.

### Limitations of the Study

POCEMON enriches biochemically recoverable transcriptional condensate material and therefore biases for retention of larger, more stable particles. Although multiple orthogonal methods support structural preservation of TCIs, there could be aspects of condensate organization which are changing during the preparation but are not being captured by the analytical methods we used, such as the loss of super-enhancer DNA. The recovered isolates should not be assumed to retain their endogenous material properties such as liquidity and therefore are unlikely to be useful for purposes such as single condensate biophysics assays. The enriched material is partially purified, and in addition to transcriptional condensates, it still retains some other co-purifying nuclear bodies. POCEMON is capturing a population average, and therefore unable to inform on coordinated bursting or other single-condensate events. Any inference of transcription dynamics from the captured material may reflect the state in the cell but formally could represent changes that occurred during the preparation. We favor the former interpretation, as the nuclear NTP concentration drops precipitously during nuclear isolation, and sonication conditions instantly dilute the extract by an order of magnitude, though this remains an important caveat. TCI DNA signal spans a relatively narrow dynamic range, and quantitative differences between individual loci are difficult to interpret. Finally, this is a steady-state readout and therefore may miss condensate nucleation features if they are rapid.

## RESOURCE AVAILABILITY

### Lead contact for reagent and resource sharing

Further information and requests for resources and reagents should be directed to and will be fulfilled by the lead contact, Dr. Alexander Ruthenburg.

### Materials availability

Reagents generated in this study are available from the lead contact upon request.

### Data and code availability

Any additional information required to reanalyze the data reported in this paper is available from the lead contact upon request.

## ACKNOWLEDGEMENTS

This study was supported by the Cancer Research Foundation (CRF) Fletcher Scholar, NSF (21-509) and NIH (R35-GM145373) grants to A.J.R. This project was supported by the National Center for Advancing Translational Sciences (NCATS) of the National Institutes of Health (NIH) through Grant Number UL1TR002389 that funds the Institute for Translational Medicine (ITM). The content is solely the responsibility of the authors and does not necessarily represent the official views of the NIH. J.V.B. was supported by the Nancy Mollin Michael Endowment Fellowship and the Dositeja Fund for Young Talents of Serbia. A.K. was supported by the Moore Foundation Postdoctoral Fellowship (310001245). J.H.S. acknowledges support from the Research Corporation for Scientific Advancement (CMC 28407), the NIH (R35GM150560), the NSF (MCB-2306187), and the University of Illinois Chicago Startup Fund. This research was supported in part by grants from the NSF (DMS-2235451) and Simons Foundation (MP-TMPS-00005320) to the NSF-Simons National Institute for Theory and Mathematics in Biology (NITMB). We are grateful to Dr. Primal de Lanorolle of UIC for his helpful insight in the development of this project. We thank Luke D. Lavis, Jonathan B. Grimm, and the Open Chemistry team (Janelia Research Campus) for the Janelia Fluor dyes. We thank Christine Labno and the University of Chicago Integrated Light Microscopy Core Facility, (RRID:SCR_019197), Cancer Center Support Grant (3P30CA014599). We are grateful for the instrumentation and expertise of the University of Chicago Advanced Electron Microscopy Core Facility (RRID:SCR_019198) for the Cryo-ET. We wish to thank P. Faber and H. Whitehurst in the University of Chicago Functional Genomics Core Facility for Illumina NovaSEQ-X sequencing.

## AUTHOR CONTRIBUTIONS

J.V.B. and A.J.R conceived the study. J.V.B. performed fractionation experiments, E14 cell line generation, immunoblots, immunofluorescence and confocal microscopy. J.V.B. analyzed and visualized sequencing data. J.G.S. and J.V.B. performed DNA-FISH experiments. J.G.S. performed all DNA-FISH analyses and cryo-ET segmentation. V.P. and J.V.B. performed cryo-ET grid preparation and cryo-ET data acquisition. A.S.K. helped with immunoblotting and provided valuable insights towards analyses and writing. F.M. and A.B. generated V6.5 cell lines. S.F.B., J.G.S., and J.H.S. contributed valuable ideas regarding experimental analyses and conceptualization. J.V.B. wrote the original draft of the manuscript. J.V.B. and A.J.R. revised the manuscript, with input from all coauthors. J.H.S. and A.J.R. obtained funding for this study.

## Materials and Methods

### Experimental model and study participant details

#### Cell culture

V6.5^49^ (gift from Jan Hendrik Spille lab, male cells) and E14Tg2a (MMRRC 015890-UCD-CELL, male cells) mESCs were cultured on 0.1% gelatin (Sigma ES-006-B) coated dishes in high glucose, pyruvate DMEM media (Gibco 11995) supplemented with 15% premium fetal bovine serum (Gibco A5670701), 1x non-essential amino acids (Gibco 11140050), 1x Penicillin/Streptomycin (Gibco 15070063), 2 mM L-glutamine (Gibco 25030081), 110 µM 2-Mercaptoethanol (Gibco 21985023), 2i (CHIR99021 and PD0325901, MCE HY-10182 and HY-10254, to 3 µM and 1 µM, respectively), and 1000 U/mL LIF (BioTechne 8878-LF). Cells were cultured at 37°C, with 5% CO_2_, in a humidified chamber. Cells were passaged every 1-2 days at ∼40,000/cm^2^ and media was changed daily.

### Method details

#### Halo-Rpb1 monoclonal cell line construction

E14Tg2a mouse ES cells were expanded for several days prior to electroporation. During expansion, conditioned medium was collected from healthy cultures and filter-sterilized for use during recovery. Post-electroporation and during early clonal stage, cells were grown in 20% conditioned media.

sgRNA (Synthego, sequence GUGCUCAGAACCCGUCCGGG) and SpCas9 protein (Synthego R20SPCAS9-SM) were prepared immediately prior to electroporation. Ribonucleoprotein complexes were assembled at a 3:1 sgRNA:Cas9 molar ratio by combining 60 pmol sgRNA with 20 pmol Cas9 per electroporation reaction, followed by addition of Resuspension Buffer R (Thermo) to a final RNP volume of 7 µL. RNP mixtures were incubated for 10 min at room temperature to allow complex formation. For each reaction, 100 pg HDR HaloTag donor DNA PCR product, amplified from a gene block fragment (gene block and primer sequences in Table S2), was added to the pre-formed RNP, followed by addition of 5 µL of the cell suspension (corresponding to 3.0 × 10^5^ cells) to generate a 12 µL electroporation mixture.

Electroporation was performed using the Neon electroporation system (Thermo) fitted with a 10 µL tip with the following parameters: 1400 V, 10 ms pulse width, 3 pulses. Immediately after electroporation, cells were transferred into individual wells of a pre-warmed gelatinized 24-well plate containing antibiotic-free medium supplemented with 20% conditioned medium. Medium was replaced the following day. Two weeks after electroporation, cells were incubated with JF646 ligand (Promega GA1120, 100 nM, 20 min) to label HaloTag-positive knock-in cells, washed, and fluorescent cells were sorted by flow cytometry. Single cells were amplified and genotyped using PCR amplification of whole insertion (primer sequences in Table S2), as well as immunoblotting.

#### POCEMON gradient fractionation and transcriptional condensate affinity purification

##### Cell preparation, treatment, and fluorescent labeling

Halo-Med1/SNAP-Rpb1 V6.5 or Halo-Rpb1 E14Tg2a ES cells, as indicated, were expanded in the days preceding fractionation or DNA-FISH experiments. Where indicated, prior to the preparation, cells were treated with DMSO (1:1000) or 10 µM triptolide (TPL, Sigma T3652-1MG) for 60 minutes (for dual gradient preparations) or 90 minutes (for POCEMON preparations); or 10% (w/v) 1,6-hexanediol (1,6-HD, Sigma 240117), or 10% (w/v) 2,5-hexanediol (2,5-HD, Sigma H11904) for 12 minutes. Where imaging of SNAP-tagged proteins was required, cells were labeled in culture with 50-100 nM SNAP-JFX-650 (gift from Luke Lavis lab)^114^ or 1 µM Halo-TMR (Promega), respectively, for 30 min at 37°C prior to harvest. For fractionation, four T75 flasks of cells per sample were detached with TrypLE (Gibco), collected by centrifugation, and the packed cell volume (PCV) was estimated (∼500 µL total per sample). All subsequent steps were performed on ice or at 4°C unless otherwise noted.

##### Isolation and sucrose cushion purification of nuclei

Cell pellets were resuspended in 3x packed cell volume of nuclei isolation buffer 1 (10 mM Na-HEPES pH 7.8, 340 mM sucrose, 10 mM KCl, 1.5 mM MgCl_2_, and 10% glycerol) as in Ruthenburg *et al.,*^115^ supplemented immediately prior to use with 0.5 mM PMSF, 0.5 mM DTT, and 1x protease inhibitor cocktail [1 x PIC (from 200x frozen stocks in DMSO): 1 mM ABESF, 0.8 μM aprotinin, 20 μM leupeptin, 15 μM pepstatin A, 40 μM bestatin, 15 μM E-64, 600 μM Benzamidine]. An equal volume of nuclei isolation buffer 2 (buffer 1 supplemented with 0.2% Triton X-100 (v/v)) was added slowly with gentle mixing to lyse the plasma membrane while maintaining nuclear integrity. Samples were incubated on ice for 12 min with occasional mixing by tube flicking, and nuclei were collected by centrifugation (1500 x g, 10 min, 4°C, Thermo Legend XTR, TX-750 swing bucket rotor). The supernatant (cytoplasmic fraction) was removed, and nuclei were resuspended in nuclei isolation buffer 1. Next, the nuclei were purified by layering over a 10 mL sucrose cushion in a 50 mL conical tube, gently disturbing the interface with a pipette, and centrifuging (1300 x g, 12 min, 4°C, Thermo Legend XTR, TX-750 swing bucket rotor). The supernatant/cushion was removed, and the purified nuclear pellet was processed immediately. For non-native preparations, purified nuclei were resuspended in crosslinked in 15 mL PBS, and 15 mL 0.2% formaldehyde (w/v, Pierce 28908) in PBS was added to each sample, for a final concentration of 0.1% formaldehyde (w/v). Samples were incubated at room temperature for 2 minutes, with mixing by manual inversion. The crosslinking reaction was quenched by addition of 2 M Tris-HCl pH 7.5 to a final 0.5 M concentration. Quenching was allowed to occur for 5 minutes at room temperature, and crosslinked nuclei were collected by centrifugation (3800 x g, 20 min, 4°C, Thermo Legend XTR, TX-750 swing bucket rotor).

##### HaloTag biotinylation, sonication, and nucleoli depletion

Purified nuclei were resuspended in 1 mL sonication buffer (0.28 M sucrose, 200 mM KOAc, 9 mM Tris pH 7.5, 0.9 mM MgCl_2_), freshly supplemented with 0.01% NP-40 and 1x PIC; total resuspension volume was ∼1.2 mL per sample including pellet contribution. Endogenous Halo-tagged proteins were biotinylated by adding HaloTag PEG-biotin (Promega G8591) to 5 µM final (from a 5 mM DMSO stock; 1:1000). Nuclei were sonicated on ice for 120 s total (10 s on / 30 s off; 37% amplitude for native and 45% for crosslinked samples, using a QSonica Q125 sonicator). For RNA-preserving preparations, RNase inhibitor (NEB M0314) was added to 1 U/µL final (from a 40 U/µL stock, 1:40). Sonicated nuclei were then incubated at room-temperature for 30 min to allow completion of the HaloTag biotinylation reaction.

Aliquots of the biotinylated sonicate were reserved for quality control and downstream assays (western blot input; DNA or RNA extraction; and imaging by crosslinking small volumes onto poly-L-lysine-coated slides, as applicable). The following steps of nucleoli depletion and the first ultracentrifugation step were adapted from a native Cajal body isolation (Lam *et al.*, 2002),^36^ with modifications. To deplete nucleoli, sucrose was adjusted to 1.0 M by adding 0.88x volume of 1.8 M sucrose buffer, then nucleoli/other very large nuclear debris were pelleted (3000 x g, 5 min, 4°C, 2 mL microcentrifuge tube, fixed angle), and the supernatant was recovered as the first nucleolus-depleted nuclear fraction (NdF). The nucleolar pellet was washed once in sucrose wash buffer (0.35 M sucrose, 10 mM Tris-HCl pH 7.5, 1 mM MgCl_2_, 1x PIC), spun (1400 x g, 2 min, 4°C), and the wash supernatant was pooled back with the initial supernatant to generate the total NdF.

##### Percoll gradient 1 to separate nuclear particles by size and density

NdF was brought to a uniform 1.0 M sucrose concentration (using 1.8 M sucrose as needed) and sucrose/Tris/MgCl_2_ buffer (1.0 M sucrose, 10 mM Tris-HCl pH 7.5, 1 mM MgCl_2_) was added to 6.175 mL. The sample was then adjusted to a final loading volume compatible with SW41 tubes by addition of 4.94 mL SP1 buffer (1.0 M sucrose, 34.2 % Percoll stock solution (23% (w/w), Cytiva 17089101), 22.2 mM Tris-HCl pH7.5, 1.11 mM MgCl_2_), 0.585 mL 20% Triton X-100 (v/v; final concentration 1%), 55 µL protease inhibitors, and (for RNA samples) 10 µL RNase inhibitor. Samples were loaded into pre-chilled SW41 inserts (Beckman Coulter 344059) and centrifuged at 37000 rpm (234116.4 rcf) for 2 h (4°C, SW41 rotor, Beckman Coulter Optima L-90K Ultracentrifuge). Ten fractions were collected from top to bottom as 1 mL aliquots (F1–F10), with fraction 10 being of variable remaining volume and representing the most sedimented material (“pellet fraction”). Defined fractions (e.g., F2/F5/F10) were optionally crosslinked on poly-L-lysine coated slides for imaging. Aliquots from each fraction were reserved for western blotting and for nucleic acid extraction (ethanol precipitation-based DNA or Trizol-chloroform-based RNA extraction, depending on the nucleic acid species being prepared for sequencing).

##### Streptavidin capture of Halo-biotinylated condensate material

For affinity enrichment, fraction 10 from the first Percoll gradient was normalized to a sample-consistent working volume (e.g., 1275 µL) using sucrose wash buffer. NaCl was added to 150 mM final (1:33 of 5 M NaCl), and samples were further supplemented with a Tris/MgCl_2_/NaCl buffer (10 mM Tris-HCl pH 7.5, 1 mM MgCl_2_, 150 mM NaCl). Dynabeads M-280 Streptavidin (Invitrogen) were equilibrated by washing once in supplemented sonication buffer, then added to samples (75 µL beads per sample). Binding was performed on ice for 45 min with gentle periodic slow mixing by hand, followed by magnetic separation and careful removal of the unbound supernatant.

Beads were washed once for 20 min in 0.14 M sucrose, 75 mM NaCl, 10 mM Tris-HCl pH 7.5, 0.915 mM MgCl_2_, 1x PIC, 0.005 % NP-40, a low-detergent, moderate-salt wash condition. Captured material was eluted either in 1x SDS-PAGE loading buffer (62.5 mM Tris-HCl pH 6.8, 2% SDS, 10% glycerol, 0.1 mg/mL bromophenol blue) and heated at 95°C for 5 min for DNA or protein analysis, or for RNA isolation, directly into Trizol (Invitrogen), followed by magnetic separation to recover the eluate. Fractions and pulldown eluates were flash-frozen in liquid nitrogen for storage prior to downstream western blotting, DNA/RNA extraction, and sequencing library preparation.

#### Percoll gradient 2 to assess perturbation effects on TCI density

F10 from gradient 1 was prewarmed to 37°C for 2 min. CaCl_2_ was added to 3.75 mM and F10 was subjected to enzymatic digestion by addition of 6 U micrococcal nuclease (Worthington Biochemical Corporation LS004798) and incubation at 37°C for 5 min for crosslinked or 2 min for native preparations. Fractions were transferred to ice, and reaction was quenched by addition of 1 mM EGTA. 300 µL dilution buffer (10 mM Tris-HCl pH7.5, 1 mM MgCl_2_) was added to the fractions prior to layering of the diluted fractions onto the gradient 2. 9 mL SP2 buffer (60% Percoll stock solution (23% (w/w), Cytiva), 0.25 M sucrose, 75 mM NaCl, 10 mM Tris-HCl pH7.5) was added to pre-chilled tubes (Beckman Coulter 355603). Samples were gently layered on top of SP2 buffer and centrifuged (25600 rpm = 60000 rcf), 15 min for crosslinked or 20 min for native preparations, 4°C, Ti 70.1 rotor, Beckman Coulter Optima L-90K Ultracentrifuge).

Eleven or twelve fractions were collected, with volumes of 0.5 mL from top to below band fraction, and 1-1.25 mL for the bottom five fractions. Volumes were kept constant as much as possible between samples from the same preparation and were documented for each preparation and sample. It was also noted which fraction or fractions contained the band fraction in different preparations. Samples were flash frozen for downstream molecular analysis.

#### DNA purification

DNA was recovered from sonicated nuclear material, NdF and gradient fractions by overnight −20°C ethanol precipitation and resuspended in 25 µL 1x TE. DNA concentration was determined using Quant-iT PicoGreen (Invitrogen P11496), according to the manufacturer’s instructions.

Lambda ladder (Thermo SM0101) spike-in was added to all DNA samples other than the fraction 10 pulldown samples according to original sample (fraction) volume, so that 0.4 ng spike-in was added per 1 mL sample. To ensure minimal length bias to Illumina PE short read sequencing, we fragmented the isolated sample and spike-in DNA by Dpn II digestion (20 U, 37°C, 10 min, NEB), followed by heat inactivation (65°C, 20 min) prior to library construction. This furnished libraries centered on 300-700 bp, within the range of efficient paired-end sequencing. For any given condition, three independent preparations from separate cultures processed all the way through the gradient 1 and pull-down were performed and subjected to sequencing.

#### RNA purification

RNA samples preserved in Trizol were subjected to Trizol-chloroform extraction, and the aqueous phase was purified using RNA Clean & Concentrator (Zymo) columns with on-column TURBO Dnase (Thermo AM2238) treatment. RNA was eluted in 25 µL warm nuclease-free water and quantified by NanoDrop. For any given condition, three independent preparations from cells were performed and subjected to sequencing.

#### Library construction and sequencing

For RNA sequencing, directional libraries were generated with the NEBNext Ultra II Directional RNA Library Prep workflow, following section 4 manufacturer’s recommendations, using fragmentation times of 15 min for sonicated nuclei samples and 8 min for Percoll-gradient fractions and fraction pulldowns. For low-input pull-down RNA, three parallel library preparations were performed per sample and pooled after second-strand synthesis at the bead-cleanup step.

For DNA libraries, end repair/A-tailing and adapter ligation were performed using NEBNext reagents and following manufacturer’s recommendations, with adapters diluted 1:10 prior to ligation. Post-ligation cleanup was carried out using AMPure XP beads (0.9x, Beckman), and libraries were eluted in 0.1x TE.

To minimize amplification bias, PCR cycle numbers were determined by qPCR test amplification using a universal primer and the appropriate indexed primer, and final library PCR was performed using a cycle number chosen to remain below plateau (empirically selected per sample). Unique NEBNext multiplex oligos for Illumina (NEB E6440S or E7335, E7500, E7710) were added to each library during amplification. Amplified libraries were purified using AMPure XP beads (0.85x), eluted in nuclease-free water, and assessed by TapeStation (Agilent). Final size selection, QC, pooling, and sequencing were performed by the institutional core facility. Libraries were sequenced on an Illumina NovaSeq-X instrument using paired-end 100 bp reads (PE100).

#### POCEMON data processing

Paired-end sequencing reads from POCEMON DNA libraries were aligned to a combined mouse mm10 and lambda phage reference genome using Bowtie2 with otherwise default parameters. Alignments were converted to BAM format, sorted, deduplicated, and indexed using SAMtools v1.15.^116^ Paired-end sequencing reads from POCEMON RNA libraries were aligned to a combined mouse mm10 and ERCC92 reference genome using HISAT2 v2.1.0^117^ with otherwise default parameters. Alignments were converted to BAM format, sorted, deduplicated, and indexed using SAMtools.^116^ Deduplicated BAM files were converted to CPM-normalized bigWig tracks using deepTools bamCoverage^118^ with 50 bp or 5 bp bins and exclusion of mm10 blacklist regions.^119^ 50 bp-binned bigWig files were used for high resolution metagene plots at 400 bp windows centered at TSSes. All other analyses used 50 bp bigWig files. For visualization and signal-based analyses, replicate summary tracks were generated by calculating the per-bin median signal across biological replicates; 75th percentile summary tracks were also generated for locus-level visualization of replicate signal variability.

#### Public ChIP-seq data processing

Public sequencing data were downloaded from the SRA and converted to FASTQ format using fasterq-dump. Single-end sequencing reads from public ChIP-seq datasets were aligned to the mouse mm10 reference genome using Bowtie2 with otherwise default parameters. Alignments were converted to BAM format, sorted, and indexed using SAMtools v1.15.^116^ ChIP and matched input BAM files were compared using deepTools bamCompare^118^ to generate log_2_(IP/Input) bigWig tracks at 1 bp binning, with CPM normalization, duplicate reads ignored, and mm10 blacklist regions^119^ excluded. ChIP-seq peaks were called from individual Pol II and Med1 immunoprecipitation (IP) replicates relative to matched input controls using MACS3 v3.0.3^120^ in broad-peak mode with –nomodel, --broad, --broad-cutoff 0.1, --extsize 300, --shift 0, --keep-dup all, and -B –SPMR. Peak files were imported into R as genomic intervals,^121^ restricted to canonical mouse chromosomes, and filtered to remove any regions overlapping the mm10 blacklist.^119^ To further reduce artifactual calls in poorly mappable sequence, average genomic mappability^122^ was calculated across each peak after extension by ±200 bp, and peaks with mean mappability below 0.8 were excluded.

#### Public TT-seq data processing

Public sequencing data reads from Shao *et al.*^107^ were downloaded from the SRA (SRR13866846) and converted to FASTQ format using fasterq-dump. Reads were aligned to the mouse mm10 reference genome using HISAT2 v2.1.0,^117^ and alignments were converted to BAM format, coordinate-sorted, and indexed with SAMtools.^116^ Alignment quality was assessed using samtools flagstat.^116^ For downstream analyses, reads with mapping quality <10 and duplicate reads were excluded, and CPM-normalized bigWig files were generated from the filtered BAM files using deepTools (v3.5.6) bamCoverage^118^ with 50 bp bins.

#### Public nuclear polyA RNA data processing

Paired-end reads from Yeom et al.^113^ (SRR17278369) were aligned to the mm10 genome using HISAT2 v2.1.0.^117^ For each sample, reads from the R1 and R2 FASTQ files were mapped as mate pairs. The resulting SAM output was converted to BAM, sorted, duplicate reads were removed, and final deduplicated BAM files were indexed using SAMtools v1.15.^116^

#### Public GRO-seq data processing

GRO-seq data from Jonkers *et al.*^77^ were downloaded from the NCBI SRA. Untreated samples consisted of SRR935093 and SRR935094, and the 5-min flavopiridol-treated samples consisted of SRR935101 and SRR935102. Adapter sequences were trimmed with cutadapt using the adapter sequence TGGAATTCTCGGGTGCCAAGG, with a minimum retained read length of 15 nt. Trimmed reads were aligned to the mm10 genome using HISAT2 v2.1.0^117^ with spliced alignment disabled (--no-spliced-alignment) to reflect the nascent, largely unspliced nature of GRO-seq reads. Alignments were sorted and indexed with SAMtools v1.15,^116^ and reads with mapping quality <10 were removed. Individual runs were processed separately for quality control and then merged by condition using SAMtools.^116^ Strand-specific genome coverage tracks were generated from the merged BAM files with deepTools bamCoverage,^118^ using CPM normalization and 5 bp bins. Separate forward- and reverse-strand bigWig files were generated for each condition, and strand orientation was verified by visual inspection of signal over annotated genes before assigning biological plus and minus tracks.

#### TCI peak calling and ChIP-TCI Venn diagrams

We deployed MACS3 peak calling^120^ over the three independent preparation replicates of the Pol II and Med1 streptavidin capture from F10 (TCI) as compared to the sonicated nuclei starting material. Comparisons to other later fractions furnished somewhat similar results, but we chose the earliest stage as the main comparison point, since intermediate stages represent partial enrichment (e.g, NdF or F10), and earlier fractions in the gradient were more limited in their coverage.

TCI DNA peaks were called from three F10PD (TCI) replicates relative to the nucleus background using MACS3 v3.0.3^120^ in broad-peak mode with –nomodel, --broad, --broad-cutoff 0.3, --extsize 300, --shift 0, --keep-dup all, and -B –SPMR. Peak files were imported into R as genomic intervals,^121^ restricted to canonical mouse chromosomes, and filtered to remove any regions overlapping the mm10 blacklist.^119^ To further reduce artifactual calls in poorly mappable sequence, average genomic mappability^122^ was calculated across each peak after extension by ±200 bp, and peaks with mean mappability below 0.8 were excluded. Filtered replicate peak sets were combined into consensus intervals by reducing the union of replicate peaks^121^ and retaining regions present in at least two of three replicates. This furnished 4733 Med1 TCI peaks and 6176 Pol II TCI peaks. Throughout this study, we refer to these high mappability MACS3 replicate consensus peaks, from POCEMON preparations from untreated cells, as Pol II or Med1 TCI regions. We use a reduced set of the union of Pol II and Med1 peaks in several analyses and refer to them as merged peaks. We also use shared Pol II and Med1 peaks in several analysis, and these are defined as overlapping peaks (by at least 1 bp).

Overlap between public ChIP-seq peaks and TCI DNA peak sets was assessed separately for Pol II and Med1 using pairwise Euler diagrams.^123,124^ In addition to the full ChIP peak set, ChIP peaks were ranked by the mean log_2_(IP/Input) signal across each called peak using replicate-specific bigWig tracks, and the top 10% highest-signal peaks were analyzed separately. For each comparison, overlap was quantified on the merged interval universe derived from the union of the two peak sets, and the observed number of shared intervals was compared with a permutation-based null distribution generated by randomizing the ChIP peak set 5,000 times across the mm10 genome with regioneR::randomizeRegions,^125^ while preserving peak widths and chromosome assignment (per.chromosome=TRUE, allow.overlaps=TRUE) and excluding blacklisted regions^119^ through masking. Empirical upper-tail p-values were calculated as !#perm ≥ obs + 1)/(0_!“#$_ + 11, and observed/expected ratios and z-scores were also reported. Observed overlaps were visualized using pairwise Euler diagrams^123,124^ parameterized from the counts of query-only, target-only, and shared intervals.

#### Metagene analysis of ChIP-seq signal at real and shuffled TCI DNA regions

To compare bulk ChIP-seq enrichment at TCI DNA genomic regions with matched genomic background regions, metagene profiles were generated using deepTools.^118^ Pol II and Med1 TCI DNA consensus peak sets were first converted to BED3 intervals, restricted to canonical mouse chromosomes, sorted, merged, and filtered to remove mm10-blacklisted regions.^119^ For each TCI DNA peak set, chromosome- and width-matched shuffled control regions were generated 50 times using bedtools shuffle^126^ with -chrom, excluding blacklisted regions^119^ and the union of real Pol II and Med1 TCI DNA intervals. Shuffled intervals from all iterations were pooled to generate a stable average background set. Pol II and Med1 ChIP-seq log_2_(IP/input) bigWig tracks were then profiled across ±5 kb windows centered on either real or shuffled TCI DNA regions using computeMatrix reference-point,^118^ with missing data treated as zero. Mean profiles were plotted with plotProfile,^118^ with real TCI DNA profiles overlaid against pooled shuffled background profiles. This analysis was performed separately for Pol II ChIP and Med1 ChIP signal at both Pol II TCI DNA and Med1 TCI DNA regions.

#### Reproducibility analysis

Reproducibility across POCEMON fractionation samples and conditions was assessed using CPM-normalized bigWig tracks. For Pol II and Med1 POCEMON DNA-seq, signal was quantified over corresponding consensus TCI DNA peak regions using deepTools multiBigwigSummary;^118^ for POCEMON RNA-seq, signal was quantified over GENCODE mm10 gene annotations. Biological replicates from nuclei, intermediate gradient fractions, F10, and final TCI samples were compared by Pearson correlation using plotCorrelation, with heatmaps generated from the resulting signal matrices. Principal component analysis was performed with deepTools plotPCA^118^ using the top 5,000 most variable regions or genes, and PCA coordinates were replotted in R for visualization. Blacklisted mm10 regions^119^ were excluded during signal quantification.

#### Genomic annotation of TCI DNA peaks

Genomic annotation of TCI DNA peak sets was performed using ChIPseeker in R^108,109^. Consensus Med1 TCI DNA, Pol II TCI DNA, and shared Pol II-Med1 TCI DNA peak intervals were annotated against the TxDb.Mmusculus.UCSC.mm10.knownGene^127^ transcript annotation, with gene identifiers mapped using org.Mm.eg.db.^128^ Peaks were assigned to genomic features using annotatePeak(), with promoter regions defined as ±2 kb from annotated transcription start sites. ChIPseeker annotation categories included promoter intervals, 5′ UTR, 3′ UTR, exon, intron, downstream regions, and distal intergenic regions. Fraction of peaks assigned to each genomic feature class was visualized as pie charts using the feature counts reported in the ChIPseeker annoStat summary.

#### cCRE enrichment analysis

Replicate consensus Pol II and Med1 peak BED files were merged into a nonredundant union peak set and analyzed against the mm10 ENCODE v4 cCRE catalog obtained from SCREEN^58^. Genomic intervals were imported as GRanges,^121^ converted from BED to 1-based coordinates, and restricted to canonical chromosomes (chr1-19, chrX, chrY). cCRE class annotations were taken from column 10 of the cCRE BED file, and unclassified entries were excluded. For each cCRE class, overlap was quantified as the number of union peaks intersecting at least one element of that class. Enrichment was assessed by comparison to a genome-wide permutation background generated by randomizing the union peak set 5,000 times with chromosome-preserving shuffling and blacklist masking.^119^ Observed/expected ratios, z-scores, and empirical one-sided enrichment p-values were computed from the permutation distribution, and p-values were corrected for multiple testing using the Benjamini–Hochberg method.

#### Nascent expression comparison of TCI promoter vs. non-TCI promoter genes

TT-seq signal was quantified over genes associated with TCI promoter peak regions. Med1 and Pol II TCI DNA consensus peak sets were imported as BED files, restricted to canonical mm10 chromosomes, assigned chromosome lengths, and independently reduced to merge overlapping intervals using GenomicRanges.^121^ A union TCI DNA peak set was then generated by combining and reducing the Med1 and Pol II peak sets. Gene annotations were obtained from TxDb.Mmusculus.UCSC.mm10.knownGene,^127^ restricted to canonical chromosomes, and transcription start sites (TSSs) were defined strand-specifically as 1 bp promoter intervals. For each peak set, genes were assigned as TCI-associated if their TSS overlapped a reduced peak extended by ±1 kb. Non-TCI promoter genes were defined as genes whose TSS did not overlap any ±1 kb-extended union TCI DNA peak. TT-seq^107^ CPM signal was imported from a CPM-normalized bigWig file and quantified as the mean signal across each gene body. Gene-level TT-seq values were log1p-transformed for visualization and clipped at the 99.5th percentile.

Distributions of TT-seq signal across Med1 TCI promoter, Pol II TCI promoter, union TCI promoter, and non-TCI promoter genes were visualized as violin plots with embedded boxplots. Statistical comparisons were performed using two-sided Wilcoxon rank-sum tests comparing each TCI promoter gene set to non-TCI promoter genes, followed by Benjamini-Hochberg correction.

#### Genome wide enrichment analysis

TCI DNA enrichment was compared between Med1 and Pol II using genome-wide tiled scatterplot analysis. Replicate log_2_ fold change (log_2_FC) bigWig tracks comparing TCI DNA to sonicated nuclear DNA were used for Med1 and Pol II POCEMON samples. The mm10 genome was divided into non-overlapping 2 kb bins across canonical chromosomes (chr1-19, chrX, chrY), and for each replicate, the mean log_2_FC signal was calculated within each bin using binnedAverage(). Replicate values were then averaged per bin to generate one mean Med1 TCI DNA log_2_FC value and one mean Pol II TCI DNA log_2_FC value for each genomic bin. Bins were annotated by overlap with promoter windows, typical enhancers, super-enhancers, and TCI DNA peak calls. Promoters were defined as ±1 kb around annotated TxDb.Mmusculus.UCSC.mm10.knownGene^127^ transcription start sites. Typical enhancer and ICM super-enhancer annotations were obtained from the M2eD2 online eRNA resource associated with Yu *et al.*^68^ Med1 and Pol II TCI DNA peak-overlap classes were assigned by overlap with consensus broad peak sets. Scatterplots show mean Pol II TCI DNA log_2_FC on the x-axis and mean Med1 TCI DNA log_2_FC on the y-axis for each 2 kb genomic bin, with Spearman correlation calculated across finite bins. Enrichment of high-signal bins was assessed using Fisher’s exact tests comparing bins with mean log_2_FC > 1 against all other bins for each genomic annotation class, followed by Benjamini–Hochberg correction.

#### Metagene analysis

Metagene profiles were generated in R from bigWig tracks imported with rtracklayer^129^ and summarized over mm10 gene models from TxDb.Mmusculus.UCSC.mm10.knownGene.^127^ After harmonization to UCSC chromosome naming, analyses were restricted to canonical chromosomes (chr1-19, chrX, chrY), and genes shorter than 1 kb were excluded. For each gene, a metagene matrix was constructed spanning 2 kb upstream of the TSS, a scaled gene body, and 2 kb downstream of the TES, using 40 bins for each flanking region and 100 bins for the gene body. Minus-strand genes were reversed to maintain a common transcriptional orientation. Mean signal per bin was extracted from each bigWig track and averaged across genes to obtain metagene profiles for bulk ChIP-seq, TCI DNA, TCI RNA, and TT-seq tracks.

For the heatmaps, rows were ordered from high to low by mean TT-seq signal across the full gene interval from TSS to TES, using a CPM-normalized 50 bp TT-seq bigWig. Heatmap signal values were clipped to the 1st–99th percentile range for visualization.

For pausing-sensitivity analyses, genes were grouped using GRO-seq-derived pausing index sensitivity, defined as log_2_(FP PI / untreated PI). Top and bottom 10% PI-sensitivity gene sets were imported from precomputed tables, matched to TxDb Entrez gene models,^127^ and plotted using the same scaled metagene layout. Mean profile comparisons between high- and low-sensitivity groups were min–max normalized per track for visualization.

High-resolution TSS-centered profiles were generated separately using deepTools computeMatrix^118^ in reference-point mode on ranked TSS BED files, and 5 bp binned bigWig input files for ChIP-seq, TCI DNA, TCI RNA and GRO-seq. For GRO-seq visualization, plus-and minus-strand 5 bp CPM bigWigs were summed to generate unstranded untreated and FP tracks. Profiles were computed over symmetric windows around the TSS using 5 bp bins, with regions kept in the predefined rank order and blacklisted regions^119^ excluded. For these analyses, genes were ranked by the selected metric, including TT-seq gene-body signal or GRO-seq-derived FP pausing-index sensitivity.

#### TCI signal stratification by promoter activity and Med1 occupancy

To examine how TCI DNA and RNA signals vary with promoter activity and Med1 occupancy, genomic regions were stratified into quartiles based on either bulk Med1 ChIP-seq signal or TT-seq signal. Analyses were restricted to canonical chromosomes. Promoters were defined as ±2 kb around annotated TSSs from TxDb.Mmusculus.UCSC.mm10.knownGene.^127^ Super-enhancers were analyzed using previously defined mESC enhancer annotations.^68^ For each region class, mean signal was quantified from bigWig tracks for Pol II TCI DNA and Med1 TCI DNA. Promoter plots were stratified by promoter Med1 ChIP signal or promoter TT-seq signal. Super-enhancer plots were stratified by the corresponding Med1 ChIP or TT-seq signal measured over each element. For visualization, TCI signals were transformed as log_2_(mean signal + 1) and winsorized at the 1st and 99th percentiles; statistical tests were performed on the untransformed mean signal values. Differences between Q1 and Q4 were assessed using two-sided Wilcoxon rank-sum tests separately for each region type and TCI signal, followed by Benjamini–Hochberg correction. Split-violin plots show Pol II TCI signal on the left and Med1 TCI signal on the right, with internal boxplots indicating the median and interquartile range.

#### Pausing index

Pol II pausing index was computed genome-wide from a CPM-normalized bulk Pol II ChIP-seq bigWig and a Pol II TCI DNA bigWig using mm10 gene models from TxDb.Mmusculus.UCSC.mm10.knownGene.^127^ Analyses were restricted to canonical chromosomes (chr1-19, chrX, chrY), and only genes with a valid pause window, a valid gene body, and a gene body length of at least 500 bp were retained. For each gene, the pause window was defined as TSS - 30 bp to TSS + 300 bp and the gene body as TSS + 300 bp to TES, with strand taken into account. Mean signal across pause and gene-body intervals was extracted from each bigWig, and pausing index was calculated as the ratio of mean pause-window signal to mean gene-body signal after addition of a pseudocount of 1e-6. To stratify genes by transcriptional output, TT-seq signal was quantified as the mean gene-body signal from a TT-seq bigWig, and genes were assigned to quartiles based on this value. For visualization, pausing indices were log_2_-transformed after addition of a pseudocount of 0.01 and winsorized at the 4th and 99.9th percentiles. Statistical comparisons between Q1 and Q4 were performed separately for bulk Pol II ChIP-derived and TCI DNA-derived pausing indices using two-sided Wilcoxon rank-sum tests, with Benjamini-Hochberg correction applied across the two tests within each panel.

#### Bidirectional promoter analysis

Promoter-class enrichment analyses were performed using mouse EPD promoter annotations (Mm_EPDnew_003, mm10).^130^ EPD transcription start sites (TSSs) were imported, converted to UCSC chromosome nomenclature, and restricted to canonical chromosomes (chr1-19, chrX, chrY). A promoter universe was defined as fixed windows spanning TSS ±1 kb around each EPD TSS. For the bidirectional promoter analysis, opposite-strand, head-to-head TSS pairs separated by ≤1 kb were identified, and the genomic intervals spanning these pairs were merged to generate a nonredundant set of bidirectional promoter regions. For the annotation-defined bidirectional promoter analysis, opposite-strand, head-to-head EPD TSS pairs separated by at most 1 kb were identified, requiring the plus-strand TSS to lie upstream of the minus-strand TSS. Genomic intervals spanning these TSS pairs were merged to generate a nonredundant set of bidirectional promoter regions, and each promoter window was classified as bidirectional or non-bidirectional based on overlap with this merged set.

For the CAGE-defined divergent promoter analysis, mouse FANTOM5 robust CAGE peaks^112^ were lifted from mm9 to mm10, and CAGE expression was quantified using the three untreated OS25 embryonic stem cell replicates from the FANTOM5 expression table.^111^ Active CAGE peaks were defined as peaks with mean OS25 expression ≥ 1 across the three replicates. Strand-specific representative CAGE TSS positions were assigned from the lifted peak coordinates. Promoter windows were classified as divergent if they contained at least one active plus-strand and one active minus-strand CAGE initiation event within 300 bp. To reduce redundancy from densely annotated loci, overlapping promoter windows were additionally collapsed into nonredundant promoter blocks using reduce() from the GenomicRanges package.^121^ Promoter blocks were classified as bidirectional or divergent if any constituent promoter window belonged to the corresponding class. Med1, Pol II, and shared Med1/Pol II peak sets were intersected with promoter windows and promoter blocks, and each promoter unit was scored as overlapped or not overlapped by each peak set. Enrichment of bidirectional or divergent promoter classes among peak-overlapped versus non-overlapped promoter units was assessed by Fisher’s exact test, with Benjamini–Hochberg correction applied across peak sets. Overlap relationships between Med1- and Pol II-associated promoter units were visualized using two-set Euler diagrams.^123,124^

#### RNA signal distribution across genic features

RNA feature distribution analysis was performed using CPM-normalized 50 bp binned bigWig tracks. Genomic annotations were derived from mm10 gene models in TxDb.Mmusculus.UCSC.mm10.knownGene^127^ and restricted to canonical chromosomes. Transcription start site regions were defined as TSS ±250 bp, and 5′ UTR, 3′ UTR, exon, intron, and intergenic regions were generated from transcript annotations. To avoid double-counting overlapping bases, feature classes were made mutually exclusive using the priority order TSS, 5′ UTR, 3′ UTR, exon, intron, and intergenic. For each RNA bigWig, signal was summed across all bases assigned to each feature class and normalized to the total signal across all classes in that sample. Feature fractions were averaged across replicates within each condition and visualized as stacked bar plots for mature nuclear RNA, nascent TT-seq RNA, Med1 TCI RNA, Pol II DMSO TCI RNA, and Pol II triptolide TCI RNA.

#### RNA signal across genes ranked by Med1 ChIP

Promoter rank plots were generated by quantifying mean Med1 ChIP-seq signal over TSS ±2 kb promoter windows for mm10 genes from TxDb.Mmusculus.UCSC.mm10.knownGene.^127^ Genes were restricted to canonical chromosomes and ranked by promoter Med1 ChIP signal. Mean TCI RNA signal was then extracted over the same promoter windows from Med1 and Pol II TCI RNA bigWig tracks. For visualization, genes were plotted in promoter Med1 ChIP rank order and TCI RNA trends were shown as LOESS-smoothed curves; signal values were winsorized prior to smoothing to limit the influence of extreme outliers.

#### Scatter plot RNA comparison

Gene-level comparisons between TCI RNA, TCI DNA, and TT-seq signal were performed over mm10 gene bodies. Gene intervals were obtained from TxDb.Mmusculus.UCSC.mm10.knownGene,^127^ restricted to canonical chromosomes (chr1-19, chrX, chrY), and annotated with gene symbols using org.Mm.eg.db^128^. Mean signal over each gene body was extracted from CPM-normalized bigWig tracks using bigWigAverageOverBed. For TCI RNA analyses, replicate Med1 and Pol II TCI RNA bigWigs were quantified separately and averaged per gene; TT-seq signal was quantified from a CPM-normalized 50 bp TT-seq bigWig. Analogous gene-level comparisons between TCI RNA and TCI DNA were performed by extracting mean gene-body signal from the corresponding TCI RNA and TCI DNA bigWigs and averaging replicate tracks where applicable. Scatter plots show log_10_-transformed CPM signal after addition of a pseudocount of 1e-4, and Spearman correlation coefficients were calculated using untransformed gene-level mean signals. Selected genes were labeled manually for visualization.

#### Differential expression analysis

Differential enrichment of RNA in TCIs was analyzed using gene-body read counts. Gene-body annotations were generated from GENCODE mm10 gene features^131^ and used as SAF intervals spanning the full annotated gene locus, thereby capturing exon and intron signal. Deduplicated RNA-seq BAM files from Pol II and Med1 fractionation experiments were counted over these gene-body intervals using featureCounts^132^ with reverse-stranded counting (-s 2), paired-end filtering, and chimeric/discordant fragments excluded. Counts from all samples were combined into a single gene-by-sample matrix and analyzed with DESeq2.^133^ For Pol II TCI RNA, fraction 10 pulldown samples were compared to the corresponding nucleoli depleted fraction (NdF) within treatment conditions. For Med1 TCI RNA, fraction 10 pulldown samples were compared to sonicated nuclei samples. DESeq2 models used a two-level design for each comparison, and log_2_ fold changes were shrunken using apeglm.^134^ Genes were considered significantly enriched or depleted using an adjusted p-value < 0.05 and absolute shrunken log_2_ fold change ≥ 1. Volcano plots show shrunken log_2_ fold change versus -log_10_ adjusted p-value, with selected top enriched and depleted genes labeled by gene symbol.

#### Gene Ontology enrichment analysis

Gene Ontology enrichment analysis was performed on significantly enriched Pol II TCI RNA genes from the DESeq2 analysis. Genes were mapped to Entrez IDs using org.Mm.eg.db,^128^ and the enrichment background was defined as all genes in the gene-count matrix that could be mapped to Entrez IDs. GO enrichment was performed using clusterProfiler::enrichGO^135^ with org.Mm.eg.db annotations and Benjamini–Hochberg correction, using Gene Ontology annotations.^136^ For visualization, significant GO terms were manually collapsed into broad functional categories, including nucleotide biosynthesis, ribosome biogenesis, RNA processing, nucleotide metabolism, and ATP/energy metabolism. For each broad category, genes were collapsed by union across all GO terms assigned to that category, and the category-level summary was plotted as -log_10_(FDR), with point size indicating the number of genes and color indicating gene ratio.

#### Gene category analysis

Gene category analysis was performed on five gene sets: TT-seq expressed genes, Pol II TCI-enriched RNA, Med1 TCI-enriched RNA, Pol II TCI promoter genes, and Med1 TCI promoter genes. Promoter-associated gene sets were defined by overlapping Pol II or Med1 broad TCI DNA peak sets with ±1 kb windows around TSSs derived from TxDb.Mmusculus.UCSC.mm10.knownGene,^127^ and assigning overlapping peaks to the corresponding genes. TT-seq expressed genes were defined from a CPM-normalized TT-seq bigWig by calculating mean signal across mm10 gene bodies, converting 50 bp bin signal to CPM/kb, and applying a fixed expression threshold of log_10_(CPM/kb + 1e-8) ≥ 0. TCI-enriched transcript sets were defined from DESeq2 significant gene tables. Genes were then classified into four categories: pluripotency, housekeeping, developmental, or other. Pluripotency genes were defined as the union of genes from Mouse MSigDB M2 gene sets with “PLURIPOTEN” in the gene-set name, retrieved using msigdbr v25.1.1.^137–141^ Housekeeping genes were defined from the Eisenberg and Levanon human housekeeping gene list^142^ after orthology-based mapping to mouse. Developmental genes were defined using Gene Ontology biological process term GO:0032502 (“developmental process”) and all descendant terms. Category assignments were summarized as counts and fractions within each gene set and visualized as stacked bar plots.

#### Gene-level predictor table and ROC AUC analysis

A gene-level predictor table was assembled for mm10 genes using TxDb.Mmusculus.UCSC.mm10.knownGene,^127^ restricted to canonical chromosomes. Genes were classified as TCI RNAs if they were significantly enriched in either Pol II or Med1 TCI RNA DESeq2 comparisons using padj < 0.05, log_2_ fold-change > 1, and baseMean ≥ 20. Super-enhancer-regulated genes were annotated by intersecting mESC super-enhancers^68^ with promoter capture Hi-C anchors from Novo *et al.*^143^ and retaining loops in which one anchor overlapped a gene promoter and the other overlapped a super-enhancer.

Candidate predictors were added to the master table, including gene-body TT-seq signal, promoter Pol II and Med1 ChIP-seq signal, promoter and gene-body TCI DNA signal, gene length, promoter CpG-island (obtained from the UCSC genome browser,^144^ mm10) overlap, Pol II pausing index, super-enhancer regulation, number of super-enhancer loops, and binary annotations for housekeeping, developmental, and pluripotency-associated genes.

Continuous count-like predictors were log1p-transformed, and pausing index was log_2_-transformed after applying a pseudocount. For each predictor independently, we calculated the receiver operating characteristic area under the curve (ROC AUC) for distinguishing TCI from non-TCI RNAs. Predictors were ranked by univariate ROC AUC and visualized as a horizontal bar plot, with AUC = 0.5 indicating no discriminative power.

#### SPRITE enrichment analysis

Multiway SPRITE clusters from Quinodoz *et al.*^81^ (mm9; 4D Nucleome 4DNESOJRTZZR) were analyzed using a custom Python pipeline that parsed raw cluster files, converted member coordinates to fixed 2 kb genomic bins, and removed duplicate loci within each cluster.

Replicate consensus Med1 and Pol II TCI DNA BED intervals were first lifted over to mm9, then subjected to the same padding and binning scheme used for SPRITE loci, and the resulting intervals were merged to define TCI DNA elements. Each SPRITE locus was annotated with the set of merged TCI DNA elements it overlapped, and cluster-level TCI DNA content was quantified as the number of distinct TCI DNA elements represented across all loci in that cluster. Summary statistics were calculated within SPRITE cluster-size bins, including the fraction of clusters containing at least 1, 2, or 3 distinct elements. Expected values were estimated from 5,000 permutations in which TCI DNA element identities assigned to SPRITE loci were shuffled while preserving chromosome identity and locus degree-bin structure. Observed values were compared with permutation-derived expectations using observed/expected ratios, z-scores, and empirical upper-tail p-values; p-values displayed on plots were corrected for multiple testing using the Benjamini-Hochberg method. For visualization, clusters were additionally grouped into mutually exclusive classes containing 1-2 or 3+ distinct TCI DNA elements, with cluster sizes above 100 combined into a single 101+ bin.

For pairwise SPRITE analysis, each 2 kb SPRITE locus was labeled according to overlap with Med1 TCI DNA intervals, Pol II TCI DNA intervals, both, or neither. Within each SPRITE cluster containing at least two labeled loci, all unordered pairs of labeled loci were enumerated and classified by pair label, cluster-size bin, and genomic distance class. Distance classes were defined as *trans* pairs or *cis* pairs separated by <10 kb, 10-100 kb, 100 kb-1 Mb, 1-10 Mb, or ≥10 Mb. Expected pair counts were estimated by label-shuffle permutations in which locus labels were randomized only among labeled loci while preserving chromosome and locus degree-bin strata. Observed pair counts were compared with permutation-derived expectations using observed/expected ratios and z-scores.

#### Region capture micro-C network analysis

Region capture Micro-C from Goel *et al.*^80^ analysis was performed in mm10 using published manual loop calls for *Klf1 and Ppm1g* loci and predefined capture windows, both of which were converted from the original mm39 coordinates to mm10 by liftOver.^144^ Analyses were restricted to canonical chromosomes (chr1-19, chrX, chrY). For each selected capture locus, all loops with at least one anchor overlapping the capture window were extracted, and overlapping anchors were merged with GenomicRanges::reduce()^121^ to define a nonredundant set of loop-anchor nodes. Anchors were annotated in two ways. First, anchors were classified by genomic element type using a priority scheme of Promoter > SuperEnhancer > TypicalEnhancer > Other, based on overlap with ±1 kb promoter windows derived from TxDb.Mmusculus.UCSC.mm10.knownGene,^127^ and typical enhancers and super-enhancers from Yu *et al.*^68^ Second, anchors were classified as TCI DNA or non- TCI DNA based on overlap with either Pol II or Med1 TCI DNA consensus peak sets. Capture-region interaction graphs were represented as undirected networks in igraph,^145–147^ and node degree was computed separately within each capture graph. For visualization, representative force-directed network layouts were generated with nodes colored either by genomic element class or by TCI DNA overlap. Summary statistics were then calculated across all selected capture loci combined. The fraction of promoter and typical-enhancer elements^68^ within the capture windows that overlapped loop anchors was compared by Fisher’s exact test. To test whether TCI DNA region-overlapping anchors occupied more connected positions within the local interaction network, node degree was compared between TCI DNA and non-TCI DNA anchors using a Wilcoxon rank-sum test.

#### Architectural factor loop anchor analysis

Ronin and YY1 loop-anchor coordinates were obtained from published loop-annotation tables from Dejosez et al.^29^ (GSM4041606) and Weintraub et al.^82^ (GSM2645440). Med1 and Pol II consensus peak BED files were imported as GRanges, converted from BED to 1-based coordinates, restricted to canonical chromosomes (chr1-19, chrX, chrY), and merged where indicated to generate a nonredundant TCI DNA peak set. To summarize overlap relationships, Ronin, YY1, and the TCI DNA peak set were reduced into a common merged-interval universe, and each interval was classified by membership in the three feature sets for Euler-diagram visualization^123,124^ of shared and unique regions. To test whether TCI DNA regions were preferentially associated with architectural loop anchors relative to other regulatory elements, each merged TCI DNA region was assigned a SCREEN cCRE class^58^ based on overlap with the mm10 ENCODE cCRE catalog using a fixed class-priority scheme, and enrichment at Ronin or YY1 anchors was evaluated against a matched regulatory null composed of non-TCI DNA SCREEN cCREs sampled to preserve cCRE class, chromosome distribution, and approximate element width. Null distributions were generated by 5,000 permutations, and observed overlaps with each anchor set were compared with matched-control expectations to calculate empirical upper-tail p-values, z-scores, and observed-to-expected enrichment ratios.

#### Genome browsers tracks

Representative genomic loci were visualized using pyGenomeTracks from CPM-normalized bigWig tracks for TCI DNA/RNA, nuclear background samples, and other indicated datasets, and log_2_(IP/input) bigWig tracks for ChIP-seq, together with BED annotations for TCI DNA peaks and gene models. For POCEMON samples, the median signal across three biological replicates was plotted as the primary track, and the 75th percentile signal across replicates was overlaid with partial transparency to indicate replicate-level signal variability.

#### Immunoblotting

Fractions (0.5% volume for sonicated nuclei, nucleoli and NdF; and a constant volume across all other fractions [F1-9, 1 mL; F10, 800-850 µL; TCI, 180 µL; supernatant, 1500 µL; wash, 850 µL]) were loaded onto 8% Tris-acetate gels at 2% of the original sample volume in 1x SDS PAGE buffer. Following gel electrophoresis at 80-100 V, proteins were transferred to 0.2 μm pore PVDF membranes (Immobilon-P, Millipore) using wet transfer (Bio-Rad) at 100 V for 1 hour on ice in Towbin’s transfer buffer containing 10% methanol. Membranes were blocked for 1 hour at room temperature in 2% ECL Prime blocking reagent (Cytiva RPN418) or 5% BSA (Dot Scientific 9048-46-8), diluted in TBS-T. After blocking, membranes were cut horizontally at defined molecular weights to enable multiplex probing of distinct targets. Membrane segments were incubated overnight at 4 °C with primary antibodies diluted in 5% BSA/TBS-T. After three 5-minute washes in TBS-T, membranes were incubated with HRP-conjugated secondary antibodies (1:10,000 dilution in 5% BSA/TBS-T) for 1 hour at room temperature. Following another series of three 5-minute washes, signal was detected using Lumigen ECL Ultra (TMA-6, Lumigen) and imaged. A list of antibodies and their working dilutions is provided in Table S1.

#### DNA-FISH probe construction

DNA-FISH probe templates (1 µg PCR product) were fragmented and aminoallyl-dUTP-labeled using DNA polymerase I and DNase I. Reactions (50 µL total) contained 1 µg DNA, 1x DNA polymerase I buffer, 50 µM dNTP mix (dATP/dCTP/dGTP each), 10 µM dTTP, 200 µM aminoallyl-dUTP (Sigma), 10 U DNA polymerase I (Thermo), and 0.04 U DNase I (Thermo).

Reactions were incubated at 15°C for 28 min (or until the desired DNA fragment size distribution peaking around 500-600 bp was reached), then placed on ice. Fragment size distribution was assessed by gel electrophoresis. Reactions were quenched by addition of EDTA to a final concentration of 10 mM. Products were purified using DNA Clean & Concentrator (Zymo) and recovered by overnight ethanol precipitation with glycogen carrier. Pellets were washed twice with 70% ethanol and resuspended in 30 µL nuclease-free water with heating at 55°C.

Fluorescent dye labeling of DNA-FISH probes was performed using NHS-ester dye chemistry. Alexa Fluor 488 NHS Ester (Invitrogen A20000) was prepared as dried aliquots by dissolving 1 mg dye in 520 µL methanol, dividing into 20 aliquots, and desiccating by vacuum centrifugation. Hybridization buffer consisted of 20% (w/v) dextran sulfate in 8x SSC. For labeling, 500-800 ng purified aminoallyl-dUTP incorporated DNA was denatured (95°C, 5 min) and snap-cooled on ice, then 3 µL of 1 M sodium bicarbonate was added to adjust the pH. One dried dye aliquot was resuspended in 2 µL DMSO and immediately mixed with the DNA, vortexed for 15 s, and incubated overnight at room temperature in the dark. The following day, labeled DNA was purified using DNA Clean & Concentrator (Zymo) and labeling efficiency was assessed by UV–Vis spectrophotometry (NanoDrop). For probe preparation, 200 ng labeled DNA was ethanol-precipitated together with Cot-1 DNA (4 µg; Thermo), salmon sperm DNA (10 µg; Thermo), and glycogen carrier, incubated at −80°C for 1 h, pelleted by centrifugation, washed twice with 70% ethanol, and air-dried protected from light. Pellets were resuspended in 50 µL formamide followed by 55°C incubation and addition of 50 µL of hybridization buffer with another incubation at 55°C and intermittent mixing.

#### DNA-FISH

Halo-Med1/SNAP-Rpb1 or Halo-Med1 V6.5 mESCs were plated onto 8-well chambered glass-bottom dishes (Cellvis) coated sequentially with poly-L-ornithine (overnight) and laminin (0.5 mg/mL, 2–4 h). Cells were labeled with either SNAP-JFX-650^114^ (for SNAP-Rpb1) or Halo-TMR ligand (for Halo-Med1; Promega) for 30-60 min at 37°C, rinsed three times in PBS, and incubated in complete medium for 2 h at 37°C to remove unbound dye. Cells were fixed in 3.2% formaldehyde (w/v, Pierce 28906) for 7 min, washed three times in PBS (3 x 5 min; all PBS washes in this protocol), permeabilized in 0.5% Triton X-100 for 10 min, and washed again three times in PBS. Fixed and permeabilized samples were incubated in 20% glycerol (≥20 min) and subjected to three freeze–thaw cycles by briefly immersing the emptied dish in liquid nitrogen (∼5 s), allowing thawing at room temperature, and re-equilibrating in glycerol between cycles, followed by PBS washes. To enhance probe accessibility, samples were treated with 0.1 M HCl (16-22 min) and washed in PBS, then equilibrated in 50% formamide/2x SSC for 5-30 min. In parallel, labeled probe DNA was denatured by incubation at 75°C for 10 minutes, and subsequently flash-cooled on ice. DNA within the cells was denatured by sequential incubation in pre-warmed 70% formamide/2x SSC (2.5 min, 75°C) and 50% formamide/2x SSC (1 min, 75°C). Denatured FISH probes were applied to denatured samples. Chambers were sealed and hybridized overnight at 37°C in a humidified chamber in the dark. The next day, hybridization buffer was removed and samples were washed in pre-warmed 50% formamide/2x SSC (37°C, 1 h), followed by five 10-min washes in 2x SSC at 37°C with gentle agitation prior to imaging. The last wash was left on the samples for confocal imaging.

#### DNA-FISH analysis

##### Nuclear segmentation and quality filtering

Nuclei were segmented in 3D from the Pol II/Med1 channel using micro-SAM^148^. Nuclear segmentations were filtered by area to remove very small or very large labels. Segmentations were revised by hand using *napari-segment-blobs-and-things-with-membranes*^149^.

##### FISH spot detection

FISH foci were detected with *U-FISH,*^150^ a deep learning-based spot detector. For every FOV the probe-channel volume was passed to UFish.predict() with intensity_threshold = 0.5. Spots outside of nuclear segmentations were discarded. Detections were further filtered as described below.

##### Pol II / Mediator puncta detection

Pol II/Mediator puncta were detected with the scikit-image Difference of Gaussians (DoG) blob detector,^151^ applied independently inside each nuclear region. Detection parameters were min_sigma = 0.5, max_sigma = 3.0, sigma_ratio = 1.25, threshold_rel = 0.3. Detections were further filtered as described below.

##### 3D Gaussian foci fitting

All foci detections (FISH and Pol II/Med1) were refined by an anisotropic 3D Gaussian fit via scipy’s optimize.curve_fit.^152^ The fit yields sub-voxel coordinates (z₀, y₀, x₀), standard deviation values for the X-Y and Z axes, an amplitude, and a constant background. From the fit results the signal-to-background ratio (SBR), signal-to-noise ratio (SNR), full-width-at-half-maximum (FWHM), and the coefficient of determination R² were calculated. Detections whose fits failed to converge were excluded from all downstream analysis.

##### Quality scoring and inclusion

For both FISH and condensate detections, a per-focus quality score was computed as the harmonic mean of the SNR and SBR. For each experiment, foci scoring above a given threshold were retained for further analysis. These thresholds were tuned manually such that foci assigned for inclusion were visually assessed to be legitimate. In the rare event that more than two high quality FISH foci were detected per nucleus, only the top two scoring foci were analyzed. To suppress likely false positives, foci above the 99th percentile peak intensity were excluded, as were foci with a FWHM below 0.2 μm or above 0.75 μm. Foci touching segmentation boundaries were also discarded.

##### Distances to Pol II / Med1 features

For each included FISH focus, the 3D Euclidean distance to every Pol II/Med1 puncta in the same nucleus was computed in physical units using the anisotropic voxel size. The X-Y half-width-at-half-maximum of the given Pol II/Med1 puncta was subtracted from this value to yield an estimate of FISH-center-to-puncta-edge distance. Negative distances were clipped to zero. The minimum distance per FISH focus was retained. For the computation of distance distributions, only FISH-puncta pairs with distances less than 5 μm were considered.

##### Random-position null model

To establish a null distribution for foci association, points were selected uniformly at random within each nucleus. For each nucleus and each Monte-Carlo iteration, the number of random points drawn equaled the number of FISH foci detected in that nucleus. Random points were rejected if they fell within two voxels of the nuclear boundary. Distances from random points to Pol II/Med1 features were computed exactly as for real FISH foci. The null was averaged over 10 random draws with deterministic seeds.

##### Radial intensity profiles

For each FISH focus, square 2D cutouts of 3 μm size were extracted from the slice containing the fitted focus center. Each cutout was background-subtracted using the per-nucleus 10th-percentile background and then normalized with the median intensity of pixels in an annulus 0.7X–0.95X of the maximum cutout radius around the focus center. Radial intensity profiles were calculated by binning pixels by their 2D distance from the focus center. Profiles were then averaged across foci by the median and standard error at each radius bin.

##### Per-focus enrichment scores

In addition to the radial profiles, a single scalar enrichment score was computed for each focus and channel. The score was computed as the median intensity in a 2×2 pixel grid around the focus center, divided by the annulus normalizer as described above.

##### Aggregate cutout images

Per-channel 2D cutouts at the fitted focus centers were normalized as above and aggregated across all included foci in an experiment by per-pixel median. Median (rather than mean) aggregation reduces the influence of outlier foci.

#### Immunofluorescence

Cells were seeded onto 0.1% gelatin-coated 35 mm glass-bottom imaging dishes (Cellvis D35-14-1.5-N) one day prior to crosslinking. Cells were crosslinked using 4% formaldehyde (w/v, Pierce 28906) for 10 minutes at room temperature, followed by 3 x 5 min PBS washes.

Crosslinked cell samples were stored at 4°C overnight or immediately subjected to immunofluorescent labeling. To permeabilize cells and block nonspecific labeling, samples were incubated with 0.5% Triton X-100 (v/v) and 10% normal goat serum or 1% BSA (Dot Scientific 9048-46-8) for 30 minutes at room temperature. Samples were washed with PBS 3 x 5 min, followed by primary antibody conjugation for 1 h at room temperature, and another set of PBS washes. Samples were incubated with the fluorescently labeled secondary antibodies matching the primary antibody host. A list of antibodies and their working dilutions is provided in table S1. Final 3 x 5-minute PBS washes were performed, and the last wash was left on the samples for subsequent confocal imaging.

#### Confocal microscopy

DNA-FISH and immunofluorescence images were collected on Stellaris 5 or Stellaris 8 (Leica Microsystems) using a 63x NA 1.40 oil immersion objective with a 2x zoom factor for FISH samples, and 3x zoom for immunofluorescence samples. Acquisition parameters were as follows: pixel size 90.19 nm for FISH data, or 47.31 nm for immunofluorescence data, 1024 × 1024 pixels, speed 200 Hz, line average 2, bit depth 8, pinhole size 1 Airy Unit. For FISH data, z-stacks were collected with step size of 0.25 - 0.3 µm.

#### Cryo-electron tomography

##### Grid Vitrification

Quantifoil grids (R2/1 carbon film on 300 copper mesh; Quantifoil Micro Tools) were prepared by glow discharging (20 watts, 30 seconds). Fraction 10 samples were diluted 1:1 with phosphate buffered saline, after which 3.5 µL were applied to grids and cryopreserved using a Vitrobot Mark IV (Thermo Fisher). The sample chamber was kept at 22 °C and 100% humidity with a 30 second wait time before blotting. Grids were either blotted with both blot pads (standard) or a single pad (back-side). For standard blotting, filter paper was applied to both pads and grids were blotted with blot force 1 and blot time of 3 seconds. For back-side blotting, the front blot pad was covered with parafilm, and filter paper was applied to the back blot pad; grids were back-blotted with a blot force of 10 and blot time of 10 seconds. All samples were plunged into liquid ethane and stored under liquid nitrogen. Frozen grids were clipped into AutoGrid Rings (Thermo Fisher).

##### Cryo-ET Data Collection

Cryo-ET data collection was performed on a Titan Krios G3i (Thermo Fisher) 300 kV transmission electron microscope using a K3 direct-electron detector in CDS mode and BioQuantum energy filter with a 20 eV slit (Gatan). Search maps were collected at magnification of 6,500 x and a physical pixel size of 13.19 Å. Search maps were used to place tilt series acquisition sites and to see an overview of condensate sizes. Tilt series were collected with Tomography 5 (Thermo Fisher) at magnification of 19,500 x and physical pixel size of 4.49 Å. Datasets were collected using dose-symmetric tilt scheme with a 3° tilt increment and angular range of ±54°. The total accumulated dose of each tilt series was 120 e-/Å^2^ with a defocus range of −5 to −8 µm. In total, 84 tilt series were obtained.

##### Tomogram Reconstruction

Tilt-series were motion corrected, aligned, and reconstructed using Tomo Live software (Thermo Fisher).^153^ Briefly, Tomo Live uses a motion correction algorithm based on *MotionCor2.*^154^ Alignment parameters are found by fiducial-free patch tracking. CTF parameters are determined for each tilt image and classical 2D fitting is used to determine defocus and astigmatism (as described in Rohou & Grigoreiff). For the final 3D reconstruction, Tomo Live utilizes simultaneous iterative reconstruction technique (SIRT) algorithm,^155^ and the final output tomogram is 4x binned.

##### Electron tomography segmentation

Each MRC file was loaded into a custom Python script as a 3D array using mrcfile.^156^ Intensities were inverted so that dense material was represented by low values throughout the pipeline.

Volumes were trimmed by 15% axially and by 55 pixels laterally to remove edge artifacts. A maximum intensity projection was then computed along the z-axis, resulting in a 2D blob silhouette. The 2D projection was passed through three sequential filters: a median filter (7 pixels) and a Gaussian filter (20 pixels) to suppress noise, and a large-scale bandpass filter (sigma = 250 pixels) to remove gradient artifacts in the sample matrix.

Otsu’s method (skimage.filters.threshold_otsu) was applied to the preprocessed projection in order to generate an initial segmentation mask. The candidate mask was then cleaned by binary erosion of 55 pixels (scipy.ndimage.binary_erosion). Geometric properties of the resulting masks were computed with skimage.measure.regionprops. Major and minor axes of the best-fit ellipse were used to characterize aggregate size. All masks were inspected by eye to ensure validity.

### Quantification and statistical analysis

Statistical analyses are described in the relevant Method details subsections and figure legends. Unless otherwise indicated, analyses were performed in R. For comparisons of signal distributions between two groups, two-sided Wilcoxon rank-sum tests were used. For categorical overlap analyses, Fisher’s exact tests were used. For genomic interval enrichment analyses, empirical p-values were calculated from permutation tests using shuffled genomic intervals that preserved chromosome identity and interval width where indicated. For differential enrichment analyses of sequencing count data, DESeq2 was used with Benjamini-Hochberg correction for multiple testing. Correlations were assessed using Spearman’s rank correlation. Exact n values, statistical tests, p-value definitions, and multiple-testing correction procedures are provided in the corresponding figure legends and Method details subsections.

## Supplemental Information

**Table S1.** Antibody dilutions.

| **Table S1. Antibody dilutions** |  |  |  |
| --- | --- | --- | --- |
| **Antibody** | **Source** | **Dilution for immunofluorescence** | **Dilution for immunoblot** |
| Rat anti-RNA pol II CTD phospho Ser5 | Active Motif Cat# 61085 | 1:250 |  |
| Mouse anti-RNA polymerase II CTD total | Cosmobio Cat# MCA-MABI0601-100-EX | 1:250 or 1:500 | 1:1000 |
| Mouse anti-RNA polymerase II CTD Ser2p | Cosmobio Cat# MCA-MABI0602-100-EX |  | 1:1000 |
| Mouse anti-RNA polymerase II CTD Ser5p | Cosmobio Cat# MCA-MABI0603-100-EX |  | 1:1000 |
| Rabbit anti-SRRM1 | Proteintech Cat# 12822-1-AP | 1:500 | 1:1000 |
| Mouse anti-NPM1 | Thermo Fisher Scientific Cat# 32-5200 | 1:100 | 1:1000 |
| Rabbit anti-Coilin | Proteintech Cat# 10967-1-AP | 1:200 | 1:1000 |
| Rabbit anti-SF3b155 | Bethyl Cat# A300-996A |  | 1:1000 |
| Rabbit anti-MED7 | Abcam Cat# ab187146 |  | 1:1000 |
| Rabbit anti-LaminA/B1/C | Abcam Cat# ab108922 |  | 1:1000 |
| Anti-HaloTag | Promega Cat# G921A |  | 1:1000 |
| Rabbit anti-H3K4me3 | Thermo Fisher Scientific Cat# PA5-27029 |  | 1:1000 |
| Goat anti-rat IgG (H+L) Alexa Fluor Plus 405 | Thermo Fisher Scientific Cat# A48261 | 1:500 |  |
| Goat anti-mouse IgG (H+L) Alexa Fluor Plus 568 | Thermo Fisher Scientific Cat# A-11004 | 1:500 |  |
| Goat anti-rabbit IgG (H+L) Alexa Fluor Plus 488 | Thermo Fisher Scientific Cat# A-11008 | 1:500 |  |
| Goat anti-mouse IgG (H+L), HRP linked | Thermo Fisher Scientific Cat# 31432 |  | 1:10000 |
| Goat anti-rabbit IgG (H+L), HRP<br>linked | Sigma-Aldrich Cat#<br>A6154 |  | 1:10000 |

**Table S2.**
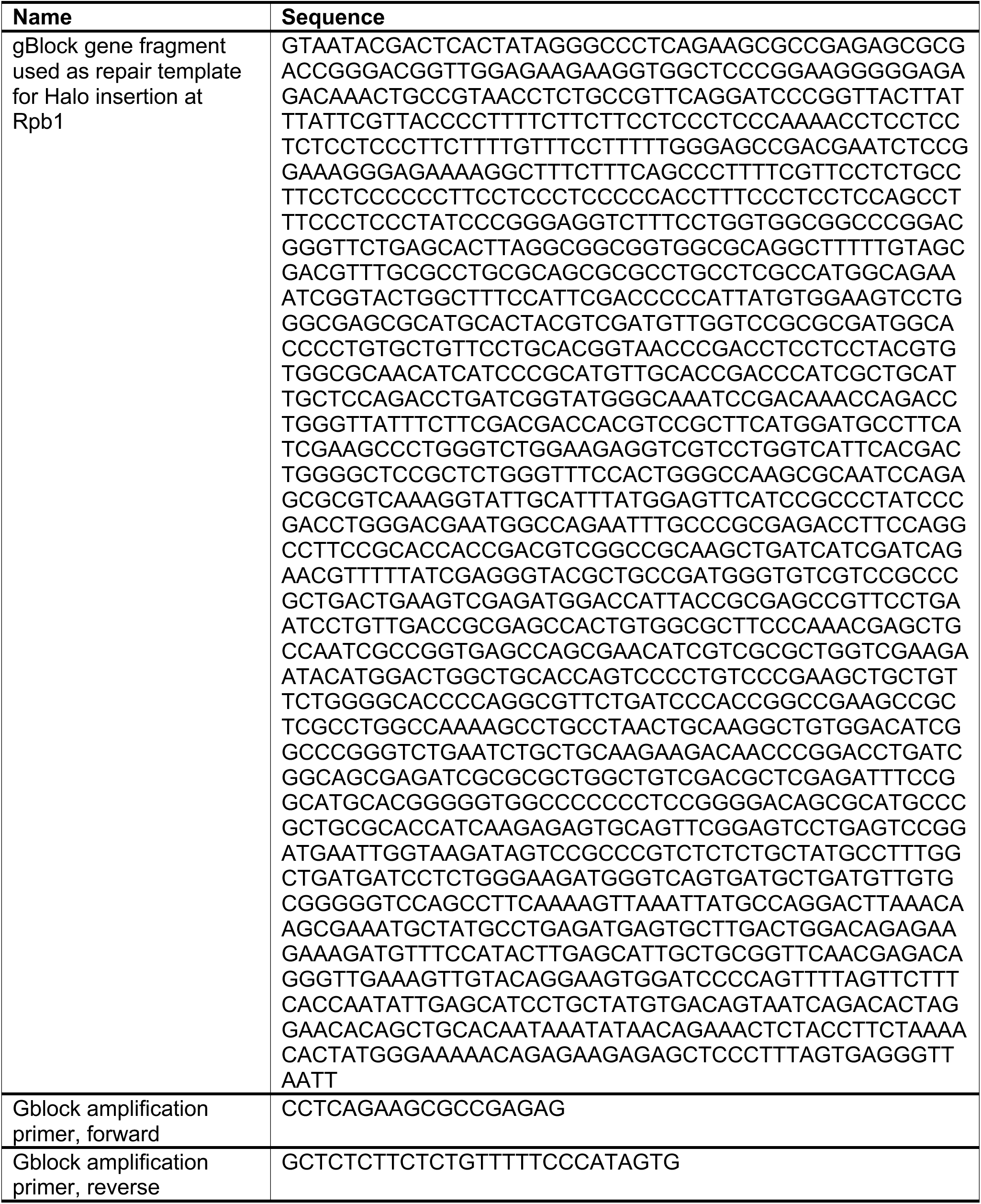
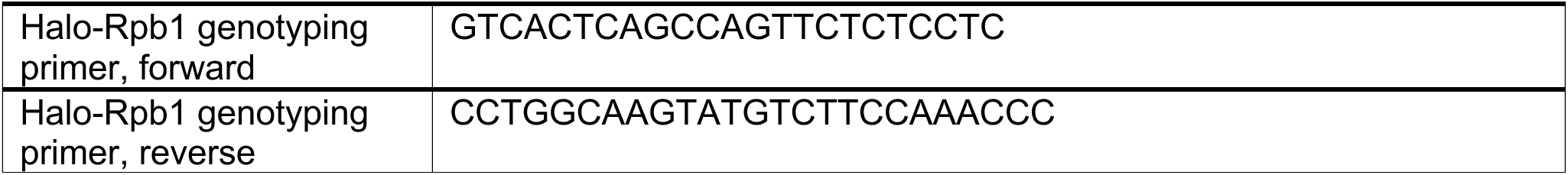
Recombinant DNA and oligonucleotide sequences.

**Figure S1.**
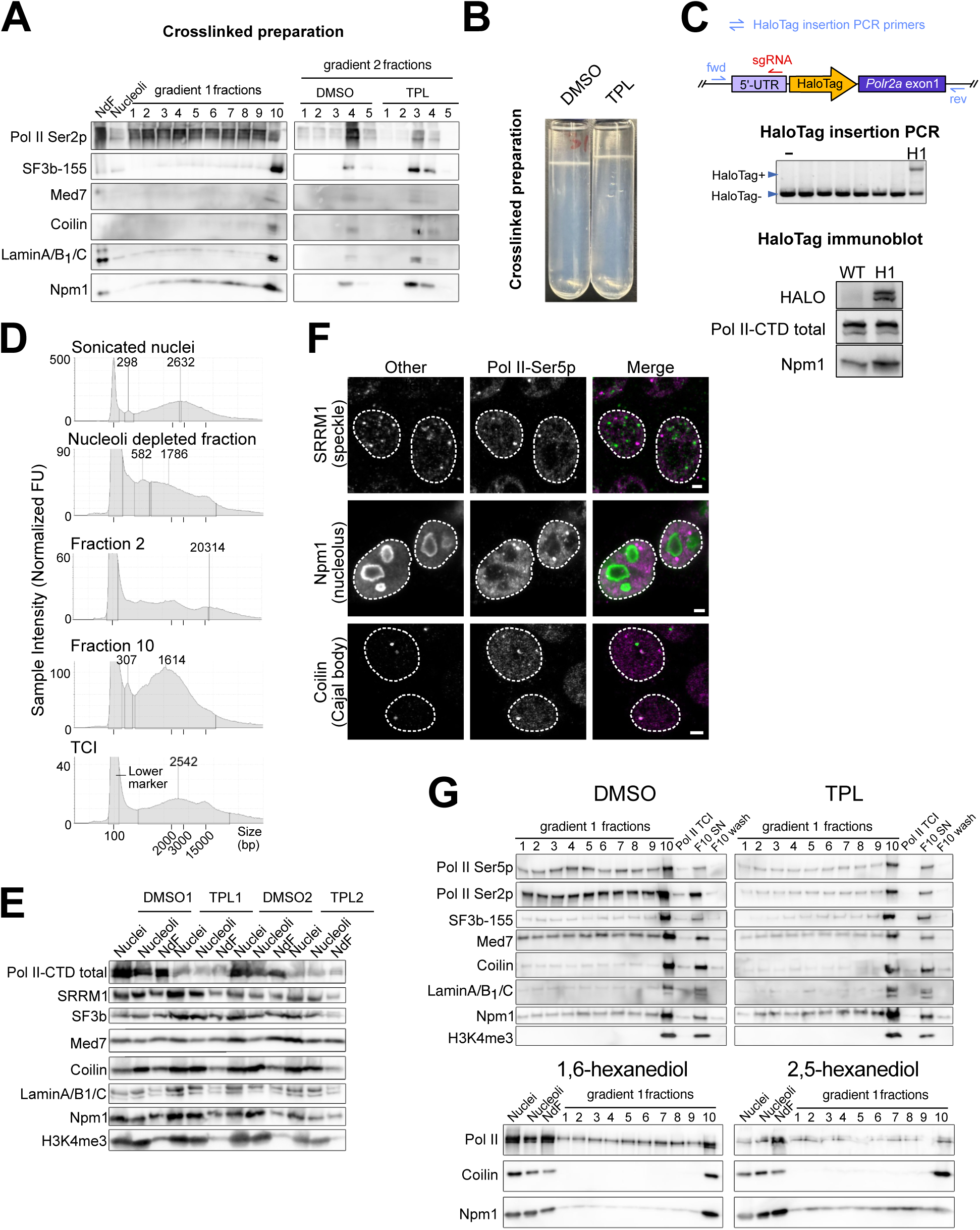
Characterization of the transcriptional condensate isolate preparation, related to Figure 1. **(A)** Immunoblot analysis of crosslinked TCI preparation. NdF, nucleoli, and relevant fractions from primary and secondary gradient are displayed for the DMSO condition, alongside relevant secondary gradient fractions from the TPL condition. **(B)** Images of representative crosslinked secondary gradient preparations following indicated treatments: 10 μM triptolide (TPL) or the equivalent volume of DMSO added to culture medium for 60 minutes prior to nuclear isolation. **(C)** Characterization of Pol II-HaloTag E14TG2a mESC line (Halo-Rpb1). Top, schematic of HaloTag heterozygous insertion at the N-terminus of endogenous Rpb1. Middle, genotyping PCR around the HaloTag insertion, confirming its presence. Bottom, immunoblot validation of Halo-Rpb1 expression relative to wild-type cells, with total Pol II-CTD and Npm1 shown as controls. The HaloTag-Med1/SNAP-Rpb1 V6.5 mESC lines are characterized in Budhathoki *et al.*^49^. **(D)** TapeStation analysis of DNA fragment size distributions across the POCEMON workflow. Representative electropherograms are shown for sonicated nuclei, NdF, F2, F10, and the final TCI sample. Size markers and dominant fragment-size peaks are indicated. **(E)** Immunoblot analysis of nuclei, nucleoli, and NdF from DMSO and TPL treated samples. **(F)** Immunofluorescence assessment of nuclear body marker localization relative to Pol II condensates in mESC nuclei. Representative images show SRRM1-positive nuclear speckles (top), Npm1-positive nucleoli (middle), and coilin-positive Cajal bodies (bottom), alongside with Pol II Ser5p, and merged channels. Images were background subtracted in FIJI with rolling-ball radius of 50 pixels. Dashed outlines indicate nuclei. Scale bar: 2 μm. **(G)** Top, immunoblot analysis of primary gradient fractions and final TCI preparation, supernatant (SN) and wash from the pulldown, from DMSO- and TPL-treated cells. Pol II Ser5p, Pol II Ser2p, SF3b-155, Med7, Coilin, Lamin A/B1/C, Npm1, and H3K4me3 were examined to assess enrichment of transcription-related factors and depletion of markers of other nuclear compartments. Bottom, immunoblot analysis of sonicated nuclei, nucleoli, and NdF, as well as primary gradient fractions, following 1,6-HD or 2,5-HD treatment.

**Figure S2.**
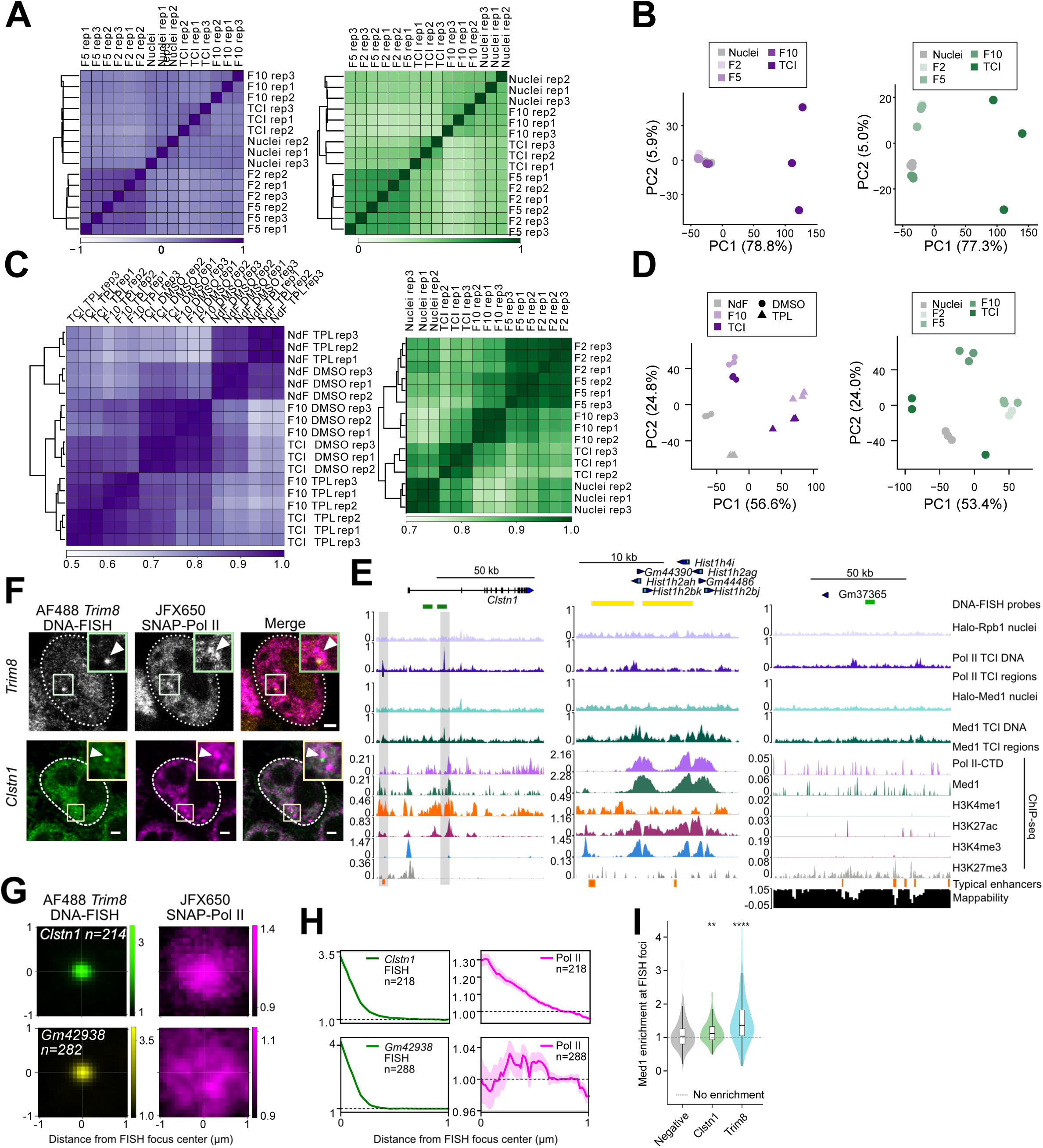
Transcriptional condensate isolate nucleic acid reproducibility and DNA-FISH validation, related to Figure 2. (**A-B**) Correlation and principal component analysis of DNA-seq profiles across fractionation samples. Signal was quantified from CPM-normalized bigWig tracks over consensus TCI DNA peak regions. (**A**) Pearson correlation heatmaps showing clustering of biological replicates and separation of nuclei, intermediate fractions, and transcriptional condensate isolates (TCIs). (**B**) Principal component analysis of the top 5,000 most variable peak regions showing fraction-specific clustering. Left, Pol II DNA; right, Med1 DNA. (**C-D**) Correlation and principal component analysis of RNA-seq profiles across fractionation samples. Signal was quantified from CPM-normalized bigWig tracks over GENCODE mm10 gene annotations. (**C**) Pearson correlation heatmaps showing clustering of biological replicates and separation of nuclei or NdF, intermediate fractions, and TCIs. (**D**) Principal component analysis of the top 5,000 most variable gene annotations showing fraction- and treatment-specific clustering. Left, Pol II RNA; right, Med1 RNA. **(E)** Genome browser tracks for the *Clstn1, Hist1*, and negative-control regions, as in Figure 2C. **(F)** Representative DNA-FISH images at the *Trim8* locus (top) and *Clstn1* locus (bottom), together with tagged Pol II signal, showing examples of DNA focus–Pol II condensate proximity. Insets show the enlarged boxed region. Scale bars: 2 μm. **(G)** Median intensity maps centered on DNA-FISH foci for the *Clstn1* locus, and the *Gm42938* locus, together with the corresponding Pol II signal. **(H)** Radial intensity quantification from panel (G), showing enrichment of Pol II signal near the *Clstn1* and *Gm42938* DNA-FISH foci. **(I)** Distributions of Med1 signal enrichment measured at the center of each FISH focus. \*\**p* < 0.01, \*\*\*\**p* < 0.0001 from Mann-Whitney U test versus the negative-control locus.

**Figure S3.**
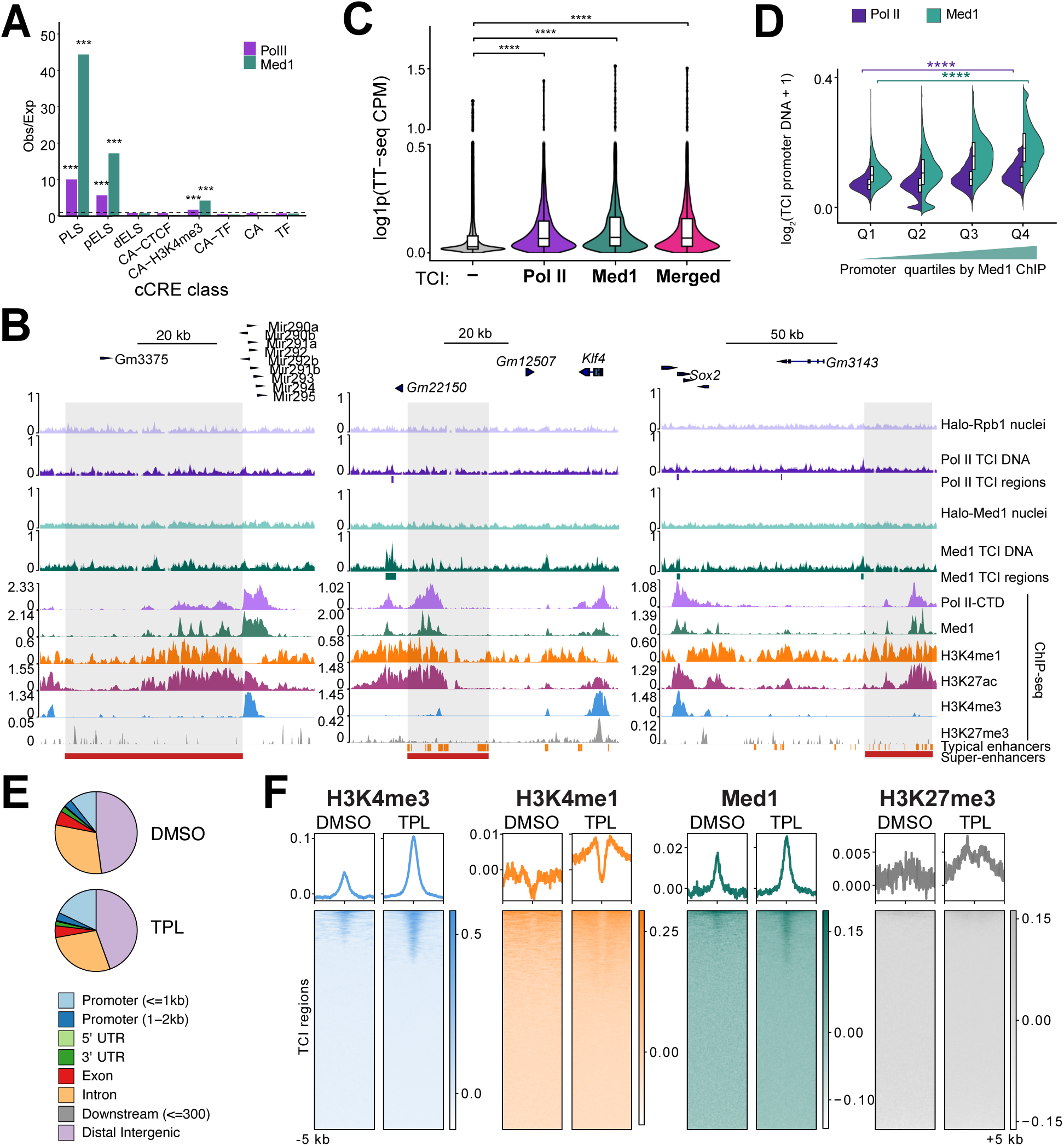
Genomic annotation and chromatin features of transcriptional condensate isolate DNA regions, related to Figure 3. **(A)** Enrichment of Pol II and Med1 TCI DNA separate region sets across ENCODE cCRE classes^58^. Bars present observed/expected overlap of each TCI DNA region set with each cCRE class. PLS – promoter-like signatures, pELS – proximal enhancer-like signatures (within 2kb of TSS), dELS – distal enhancer-like signature, CA-CTCF – chromatin accessibility + CTCF, CA-H3K4me3 – chromatin accessibility + H3K4me3, CA-TF – chromatin accessibility + transcription factor, CA – chromatin accessibility, TF – transcription factor. \*\*\**p* < 0.001 based on permutation-derived empirical upper-tail *p*-values, corrected for multiple testing using the Benjamini-Hochberg method. **(B)** Browser tracks of representative, well studied mESC super-enhancer regions at the Mir290-295 locus (left), *Klf4* locus (middle), and *Sox2* locus (right), as in Figure 2C. **(C)** Nascent transcription levels^107^ of non-TCI and TCI genes, defined by promoter overlap with separate or merged Pol II and Med1 TCI DNA region sets; range is truncated to 99.5% of all data for clarity of display. \*\*\*\**p* < 0.0001 based on two-sided Wilcoxon rank-sum tests with Benjamini-Hochberg correction. **(D)** Violin plots of Pol II and Med1 TCI signal at promoters stratified by promoter quartiles ranked by Med1 ChIP-seq.^104^ \*\*\*\**p* < 0.0001 from two-sided Wilcoxon rank-sum test with Benjamini-Hochberg correction comparing Q1 and Q4; log2-transformed CPM-normalized signal was winsorized at the 1st and 99th percentiles for clear visualization of the bulk of the distribution. **(E)** Genomic annotation of UCSC Known Gene elements for Pol II TCI DNA regions from DMSO or TPL treated cells. **(F)** ChIP-seq metaprofiles and heatmaps of promoter- and enhancer-associated chromatin marks^104–106^ presented as log_2_(IP/Input) centered on Pol II TCI regions from DMSO or TPL treated cells, spanning 5 kb upstream and downstream.

**Figure S4.**
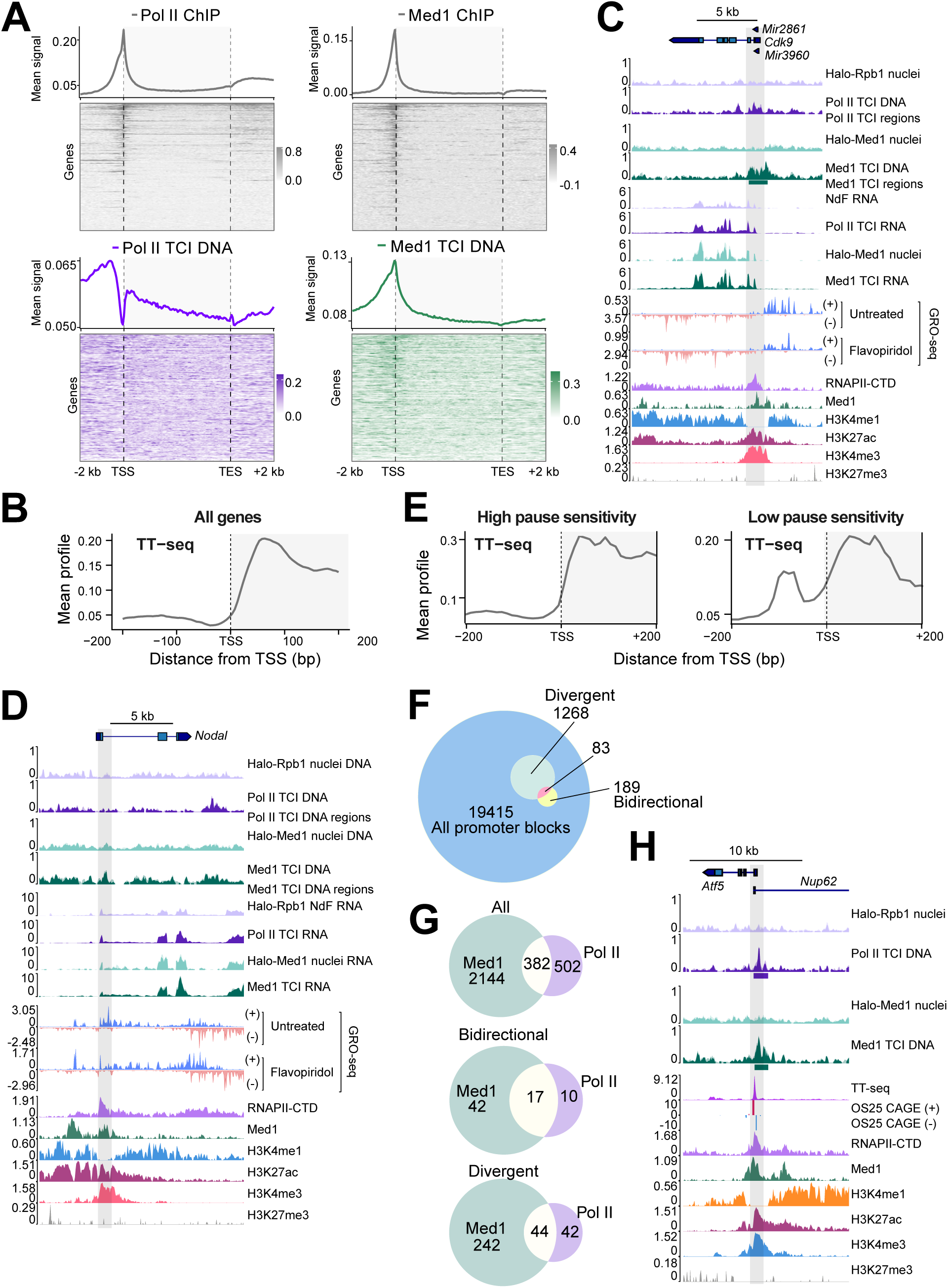
Chromatin and transcriptional features associated with Med1 and Pol II TCI partitioning, related to Figure 4. **(A)** Metagene profiles as in Figure 4A, shown along with heatmaps arranged by decreasing gene body TT-seq signal.^107^ **(B)** Zoomed metagene profiles of all genes across a 400 bp window centered on the TSS, shown for nascent RNA from TT-seq^107^ as mean CPM-normalized signal. (**C-D**) Genome browser tracks comparing nuclear input, Pol II TCI DNA, Pol II TCI peaks, Med1 TCI DNA, Med1 TCI peaks, corresponding TCI RNA tracks, stranded GRO-seq CPM-normalized signal from Jonkers *et al.*^77^ and ChIP-seq as in Figure 2C.^104–106^ (**C**) Representative pause-insensitive *Cdk9* locus, showing no change between untreated and flavopiridol (FP) conditions. (**D**) Representative pause-sensitive *Nodal* locus, displaying a prominent shift in GRO-seq^77^ signal between untreated and FP conditions. **(E)** Metagene profiles as in panel (B), shown separately for high- and low-pause sensitivity genes, as in Figure 4D and E. **(F)** Euler diagram of collapsed promoter regions for all, divergent and bidirectional promoters used in Figure 4F. **(G)** Euler diagrams of Pol II and Med1 TCI promoter regions, defined by TCI DNA region overlap, across all, bidirectional, and divergent promoter groups. Bidirectional promoters show highest overlap between Pol II and Med1 TCI. **(H)** Genome browser track displaying an example divergent promoter. Tracks show nuclear input, Pol II TCI DNA, Pol II TCI peaks, Med1 TCI DNA, Med1 TCI peaks, TT-seq,^107^ OS25 CAGE,^111^ and public ChIP-seq^104–106^ as in Figure 2C.

**Figure S5.**
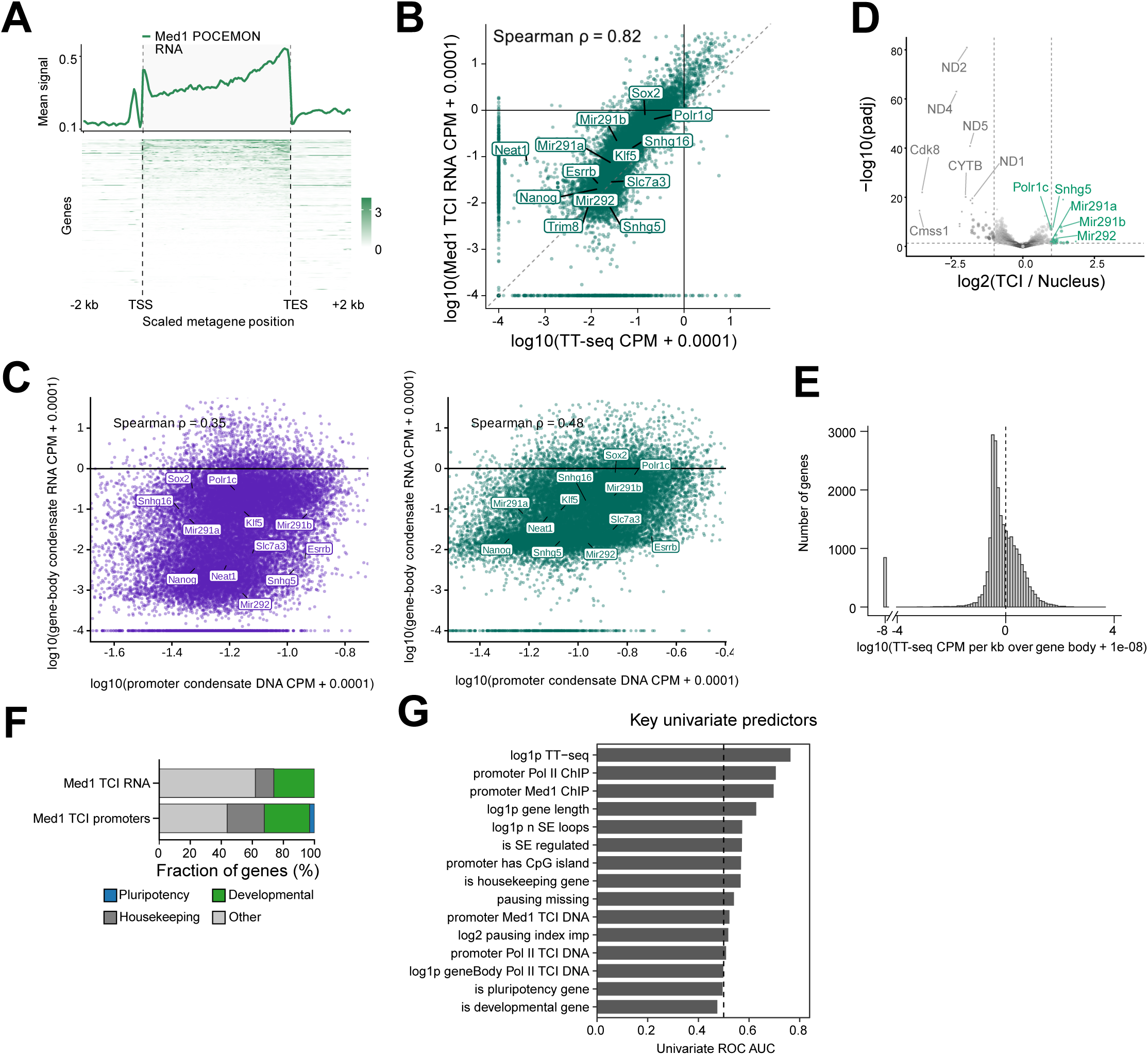
Mediator transcriptional condensate isolate RNA analyses complement Pol II TCI RNA analyses, related to Figure 5. **(A)** Metagene profile and heatmap of CPM-normalized Med1 TCI RNA signal across all genes, ranked by TT-seq^107^ gene-body signal. Genes shorter than 1 kb were excluded from the analysis. **(B)** Scatterplot comparing mean Med1 TCI RNA signal and TT-seq^107^ signal across genes, with selected representative loci labeled. **(C)** Scatterplot comparing mean Pol II (left) or Med1 (right) TCI RNA at gene bodies to the corresponding TCI DNA signal at promoters (defined as TSS ± 2kb), plotted as replicate average log_10_(CPM + 0.0001), with selected representative loci labeled. Axes were truncated at the 3.5th and 99.9th percentiles for visualization only; correlations were calculated before axis truncation. **(D)** Differential expression analysis showing genes enriched or depleted in Med1 TCI RNA relative to RNA from sonicated nuclei. Genes are plotted by apeglm-shrunken log_2_ fold change and −log_10_ adjusted p-value. Significant enriched or depleted genes were defined as adjusted p-value < 0.05 and absolute shrunken log2 fold change ≥ 1; selected genes are labeled. Note that mitochondrial transcripts were markedly depleted in Med1 TCI RNA, further supporting the nuclear specificity of the purification. **(E)** Distribution of TT-seq^107^ gene-body signal used to define expressed genes for the analysis in Figure 5H. The dashed line indicates the fixed expression cutoff. **(F)** Gene category composition of Med1 TCI RNA enriched relative to sonicated nuclear RNA, or genes whose promoter overlapped Med1 TCI DNA regions. **(G)** Full panel of key univariate predictors used in the ROC AUC model displayed in Figure 5I. The dashed line denotes an AUC of 0.5, corresponding to random classification performance.

**Figure S6.**
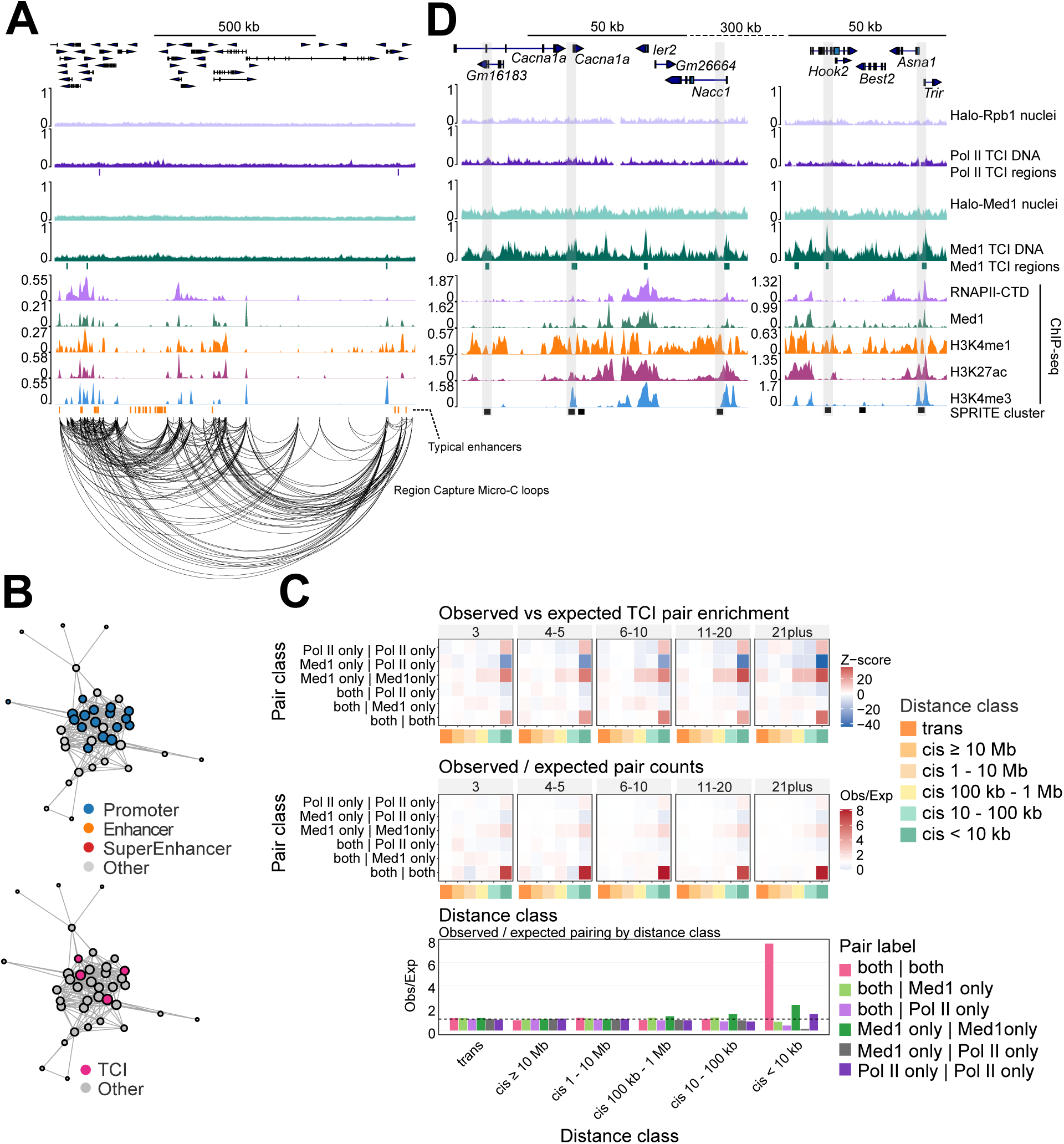
Transcriptional condensate isolate DNA regions participate in local multiway chromatin interactions, related to Figure 6. **(A)** Genome browser track view across a ∼1 Mb locus centered on *Ppm1g* showing Pol II and Med1 TCI DNA, corresponding TCI regions, public ChIP-seq tracks,^104–106^ enhancer annotations^68^, and region capture micro-C chromatin loops from Goel *et al.^80^* **(B)** Network representation of loop anchors from the captured *Ppm1g* locus. For visual clarity, loop anchors were represented as nodes, with node size proportional to interaction degree, and loops as edges. Top, anchors colored by annotation class (promoter, enhancer, super-enhancer^68^, or other). Bottom, anchors colored by overlap with merged TCI regions. **(C)** Pairwise co-occurrence analysis of Med1 and Pol II TCI DNA-overlapping loci within multiway SPRITE clusters. Each 2 kb SPRITE locus was classified by overlap with Med1 and Pol II TCI DNA regions. Pair counts were stratified by SPRITE cluster size and genomic distance class. Heatmaps show Z-scores and observed/expected ratios relative to label-shuffle permutations. Bottom, observed/expected pair enrichment summarized by genomic distance class. **(D)** Genome browser tracks of two connected gene-rich regions displaying Pol II and Med1 TCI DNA, corresponding TCI regions, public ChIP-seq tracks,^104–106^ and a single SPRITE cluster (#6_2p5-2_DPM6.YbotE61.Odd2Bo20.Even2Bo25.Odd2Bo4) from Quinodoz *et al.*^81^ Several TCI region-overlapping promoters establish a multiway chromatin interaction, based on colocalization within the same SPRITE cluster.

## References

1. Iborra, F.J., Pombo, A., Jackson, D.A., and Cook, P.R. (1996). Active RNA polymerases are localized within discrete transcription “factories’ in human nuclei. J Cell Sci 109 (Pt 6), 1427–1436. 10.1242/jcs.109.6.1427.

2. Jackson, D.A., Hassan, A.B., Errington, R.J., and Cook, P.R. (1993). Visualization of focal sites of transcription within human nuclei. EMBO J 12, 1059–1065.

3. Chong, S., Dugast-Darzacq, C., Liu, Z., Dong, P., Dailey, G.M., Cattoglio, C., Heckert, A., Banala, S., Lavis, L., Darzacq, X., and Tjian, R. (2018). Imaging dynamic and selective low-complexity domain interactions that control gene transcription. Science 361. 10.1126/science.aar2555.

4. Mir, M., Stadler, M.R., Ortiz, S.A., Hannon, C.E., Harrison, M.M., Darzacq, X., and Eisen, M.B. (2018). Dynamic multifactor hubs interact transiently with sites of active transcription in. Elife 7. 10.7554/eLife.40497.

5. Esbin, M.N., Cookis, T., Anantakrishnan, S., Abidi, A.A., Karr, J., Cattoglio, C., Darzacq, X., and Tjian, R. (2025). Assembly and Dynamics of Transcription Initiation Complexes. Annu Rev Biochem 94, 305–331. 10.1146/annurev-biochem-072324-035226.

6. Kwon, I., Kato, M., Xiang, S., Wu, L., Theodoropoulos, P., Mirzaei, H., Han, T., Xie, S., Corden, J.L., and McKnight, S.L. (2013). Phosphorylation-regulated binding of RNA polymerase II to fibrous polymers of low-complexity domains. Cell 155, 1049–1060. 10.1016/j.cell.2013.10.033.

7. Sabari, B.R., Dall’Agnese, A., Boija, A., Klein, I.A., Coffey, E.L., Shrinivas, K., Abraham, B.J., Hannett, N.M., Zamudio, A.V., Manteiga, J.C., et al. (2018). Coactivator condensation at super-enhancers links phase separation and gene control. Science 361. 10.1126/science.aar3958.

8. Hnisz, D., Shrinivas, K., Young, R.A., Chakraborty, A.K., and Sharp, P.A. (2017). A Phase Separation Model for Transcriptional Control. Cell 169, 13–23. 10.1016/j.cell.2017.02.007.

9. Boehning, M., Dugast-Darzacq, C., Rankovic, M., Hansen, A.S., Yu, T., Marie-Nelly, H., McSwiggen, D.T., Kokic, G., Dailey, G.M., Cramer, P., et al. (2018). RNA polymerase II clustering through carboxy-terminal domain phase separation. Nat Struct Mol Biol 25, 833–840. 10.1038/s41594-018-0112-y.

10. Cho, W.K., Spille, J.H., Hecht, M., Lee, C., Li, C., Grube, V., and Cisse, I.I. (2018). Mediator and RNA polymerase II clusters associate in transcription-dependent condensates. Science 361, 412–415. 10.1126/science.aar4199.

11. Banani, S.F., Lee, H.O., Hyman, A.A., and Rosen, M.K. (2017). Biomolecular condensates: organizers of cellular biochemistry. Nat Rev Mol Cell Biol 18, 285–298. 10.1038/nrm.2017.7.

12. Mittag, T., and Pappu, R.V. (2022). A conceptual framework for understanding phase separation and addressing open questions and challenges. Mol Cell 82, 2201–2214. 10.1016/j.molcel.2022.05.018.

13. Narlikar, G.J., Myong, S., Larson, D., Maeshima, K., Francis, N., Rippe, K., Sabari, B., Strader, L., and Tjian, R. (2021). Is transcriptional regulation just going through a phase? Mol Cell 81, 1579–1585. 10.1016/j.molcel.2021.03.046.

14. Rippe, K., and Papantonis, A. (2025). RNA polymerase II transcription compartments - from factories to condensates. Nat Rev Genet 26, 775–788. 10.1038/s41576-025-00859-6.

15. Banani, S.F., Afeyan, L.K., Hawken, S.W., Henninger, J.E., Dall’Agnese, A., Clark, V.E., Platt, J.M., Oksuz, O., Hannett, N.M., Sagi, I., et al. (2022). Genetic variation associated with condensate dysregulation in disease. Dev Cell 57, 1776–1788.e1778. 10.1016/j.devcel.2022.06.010.

16. Lyons, H., Pradhan, P., Prakasam, G., Vashishtha, S., Li, X., Eppert, M., Fornero, C., Tcheuyap, V.T., McGlynn, K., Yu, Z., et al. (2025). RNA polymerase II partitioning is a shared feature of diverse oncofusion condensates. Cell 188, 3843–3862.e3828. 10.1016/j.cell.2025.04.002.

17. Lyons, H., Veettil, R.T., Pradhan, P., Fornero, C., De La Cruz, N., Ito, K., Eppert, M., Roeder, R.G., and Sabari, B.R. (2023). Functional partitioning of transcriptional regulators by patterned charge blocks. Cell 186, 327–345.e328. 10.1016/j.cell.2022.12.013.

18. De La Cruz, N., Pradhan, P., Veettil, R.T., Conti, B.A., Oppikofer, M., and Sabari, B.R. (2024). Disorder-mediated interactions target proteins to specific condensates. Mol Cell 84, 3497–3512 e3499. 10.1016/j.molcel.2024.08.017.

19. Trojanowski, J., Frank, L., Rademacher, A., Mucke, N., Grigaitis, P., and Rippe, K. (2022). Transcription activation is enhanced by multivalent interactions independent of phase separation. Mol Cell 82, 1878–1893 e1810. 10.1016/j.molcel.2022.04.017.

20. Lee, C., Quintana, A., Suppanz, I., Gomez-Auli, A., Mittler, G., and Cissé, I.I. (2024). Light-induced targeting enables proteomics on endogenous condensates. Cell 187, 7079–7090.e7017. 10.1016/j.cell.2024.09.040.

21. Wei, M.T., Chang, Y.C., Shimobayashi, S.F., Shin, Y., Strom, A.R., and Brangwynne, C.P. (2020). Nucleated transcriptional condensates amplify gene expression. Nat Cell Biol 22, 1187–1196. 10.1038/s41556-020-00578-6.

22. Boija, A., Klein, I.A., Sabari, B.R., Dall’Agnese, A., Coffey, E.L., Zamudio, A.V., Li, C.H., Shrinivas, K., Manteiga, J.C., Hannett, N.M., et al. (2018). Transcription Factors Activate Genes through the Phase-Separation Capacity of Their Activation Domains. Cell 175, 1842–1855.e1816. 10.1016/j.cell.2018.10.042.

23. Du, M., Stitzinger, S.H., Spille, J.H., Cho, W.K., Lee, C., Hijaz, M., Quintana, A., and Cissé, I.I. (2024). Direct observation of a condensate effect on super-enhancer controlled gene bursting. Cell 187, 2595–2598. 10.1016/j.cell.2024.04.001.

24. Cho, W.K., Jayanth, N., English, B.P., Inoue, T., Andrews, J.O., Conway, W., Grimm, J.B., Spille, J.H., Lavis, L.D., Lionnet, T., and Cisse, I.I. (2016). RNA Polymerase II cluster dynamics predict mRNA output in living cells. Elife 5. 10.7554/eLife.13617.

25. Mehta, S., and Zhang, J. (2022). Liquid-liquid phase separation drives cellular function and dysfunction in cancer. Nat Rev Cancer 22, 239–252. 10.1038/s41568-022-00444-7.

26. Zamudio, A.V., Dall’Agnese, A., Henninger, J.E., Manteiga, J.C., Afeyan, L.K., Hannett, N.M., Coffey, E.L., Li, C.H., Oksuz, O., Sabari, B.R., et al. (2019). Mediator Condensates Localize Signaling Factors to Key Cell Identity Genes. Mol Cell 76, 753–766 e756. 10.1016/j.molcel.2019.08.016.

27. Gan, P., Eppert, M., De La Cruz, N., Lyons, H., Shah, A.M., Veettil, R.T., Chen, K., Pradhan, P., Bezprozvannaya, S., Xu, L., et al. (2024). Coactivator condensation drives cardiovascular cell lineage specification. Sci Adv 10, eadk7160. 10.1126/sciadv.adk7160.

28. Lv, J., Maher, K.A., Dong, L., Valentine, V., Staller, S., Veluchamy, A., Tian, L., Kim, Y., Ju, B., Valentine, M., et al. (2026). 3D-super-enhancers are condensate-associated cis-regulatory communities. Nucleic Acids Res 54. 10.1093/nar/gkag191.

29. Dejosez, M., Dall’Agnese, A., Ramamoorthy, M., Platt, J., Yin, X., Hogan, M., Brosh, R., Weintraub, A.S., Hnisz, D., Abraham, B.J., et al. (2023). Regulatory architecture of housekeeping genes is driven by promoter assemblies. Cell Rep 42, 112505. 10.1016/j.celrep.2023.112505.

30. Caudron-Herger, M., Cook, P.R., Rippe, K., and Papantonis, A. (2015). Dissecting the nascent human transcriptome by analysing the RNA content of transcription factories. Nucleic Acids Res 43, e95. 10.1093/nar/gkv390.

31. Cisse, I.I., Izeddin, I., Causse, S.Z., Boudarene, L., Senecal, A., Muresan, L., Dugast-Darzacq, C., Hajj, B., Dahan, M., and Darzacq, X. (2013). Real-time dynamics of RNA polymerase II clustering in live human cells. Science 341, 664–667. 10.1126/science.1239053.

32. Li, J., Dong, A., Saydaminova, K., Chang, H., Wang, G., Ochiai, H., Yamamoto, T., and Pertsinidis, A. (2019). Single-Molecule Nanoscopy Elucidates RNA Polymerase II Transcription at Single Genes in Live Cells. Cell 178, 491–506.e428. 10.1016/j.cell.2019.05.029.

33. Forero-Quintero, L.S., Raymond, W., Handa, T., Saxton, M.N., Morisaki, T., Kimura, H., Bertrand, E., Munsky, B., and Stasevich, T.J. (2021). Live-cell imaging reveals the spatiotemporal organization of endogenous RNA polymerase II phosphorylation at a single gene. Nat Commun 12, 3158. 10.1038/s41467-021-23417-0.

34. Lewis, B.A., Das, S.K., Jha, R.K., and Levens, D. (2023). Self-assembly of promoter DNA and RNA Pol II machinery into transcriptionally active biomolecular condensates. Sci Adv 9, eadi4565. 10.1126/sciadv.adi4565.

35. Imada, T., Shimi, T., Kaiho, A., Saeki, Y., and Kimura, H. (2021). RNA polymerase II condensate formation and association with Cajal and histone locus bodies in living human cells. Genes Cells 26, 298–312. 10.1111/gtc.12840.

36. Lam, Y.W., Lyon, C.E., and Lamond, A.I. (2002). Large-scale isolation of Cajal bodies from HeLa cells. Mol Biol Cell 13, 2461–2473. 10.1091/mbc.02-03-0034.

37. Muramatsu, M., Smetana, K., and Busch, H. (1963). Quantitative Aspects of Isolation of Nucleoli of the Walker Carcinosarcoma and Liver of the Rat. Cancer Res 23, 510–518.

38. Mintz, P.J., Patterson, S.D., Neuwald, A.F., Spahr, C.S., and Spector, D.L. (1999). Purification and biochemical characterization of interchromatin granule clusters. EMBO J 18, 4308–4320. 10.1093/emboj/18.15.4308.

39. Khong, A., Matheny, T., Jain, S., Mitchell, S.F., Wheeler, J.R., and Parker, R. (2017). The Stress Granule Transcriptome Reveals Principles of mRNA Accumulation in Stress Granules. Mol Cell 68, 808–820 e805. 10.1016/j.molcel.2017.10.015.

40. Melnik, S., Deng, B., Papantonis, A., Baboo, S., Carr, I.M., and Cook, P.R. (2011). The proteomes of transcription factories containing RNA polymerases I, II or III. Nat Methods 8, 963–968. 10.1038/nmeth.1705.

41. Egloff, S., Dienstbier, M., and Murphy, S. (2012). Updating the RNA polymerase CTD code: adding gene-specific layers. Trends Genet 28, 333–341. 10.1016/j.tig.2012.03.007.

42. Mitrea, D.M., Cika, J.A., Guy, C.S., Ban, D., Banerjee, P.R., Stanley, C.B., Nourse, A., Deniz, A.A., and Kriwacki, R.W. (2016). Nucleophosmin integrates within the nucleolus via multi-modal interactions with proteins displaying R-rich linear motifs and rRNA. Elife 5. 10.7554/eLife.13571.

43. Shandilya, J., Swaminathan, V., Gadad, S.S., Choudhari, R., Kodaganur, G.S., and Kundu, T.K. (2009). Acetylated NPM1 localizes in the nucleoplasm and regulates transcriptional activation of genes implicated in oral cancer manifestation. Mol Cell Biol 29, 5115–5127. 10.1128/MCB.01969-08.

44. Andrade, L.E., Chan, E.K., Raska, I., Peebles, C.L., Roos, G., and Tan, E.M. (1991). Human autoantibody to a novel protein of the nuclear coiled body: immunological characterization and cDNA cloning of p80-coilin. J Exp Med 173, 1407–1419. 10.1084/jem.173.6.1407.

45. Machyna, M., Neugebauer, K.M., and Staněk, D. (2015). Coilin: The first 25 years. RNA Biol 12, 590–596. 10.1080/15476286.2015.1034923.

46. Courchaine, E., Gelles-Watnick, S., Machyna, M., Straube, K., Sauyet, S., Enright, J., and Neugebauer, K.M. (2022). The coilin N-terminus mediates multivalent interactions between coilin and Nopp140 to form and maintain Cajal bodies. Nat Commun 13, 6005. 10.1038/s41467-022-33434-2.

47. Blencowe, B.J., Issner, R., Nickerson, J.A., and Sharp, P.A. (1998). A coactivator of pre-mRNA splicing. Genes Dev 12, 996–1009. 10.1101/gad.12.7.996.

48. Ilik, İ., Malszycki, M., Lübke, A.K., Schade, C., Meierhofer, D., and Aktaş, T. (2020). SON and SRRM2 are essential for nuclear speckle formation. Elife 9. 10.7554/eLife.60579.

49. Spille, J.-H., Budhathoki, A., Pandey, G., Medhanie, F., Ngo, G.-H., and Mahmood, S. (2025). Chromatin Binding Enriches RNA Polymerase II in Transcription Condensates. 10.21203/rs.3.rs-7302634/v1.

50. White, A.E., Leslie, M.E., Calvi, B.R., Marzluff, W.F., and Duronio, R.J. (2007). Developmental and cell cycle regulation of the Drosophila histone locus body. Mol Biol Cell 18, 2491–2502. 10.1091/mbc.e06-11-1033.

51. Marmolejo, C.O., Sanchez, C., Helms, E., McEvoy, M.J., Lee, J., Werner, M., Roberts, P., Hamperl, S., and Saldivar, J.C. (2026). Precise control of transcription condensates across S phase balances linker histone expression with DNA replication, ensuring genome stability. Mol Cell 86, 640–655 e646. 10.1016/j.molcel.2026.01.005.

52. Suzuki, H., Abe, R., Shimada, M., Hirose, T., Hirose, H., Noguchi, K., Ike, Y., Yasui, N., Furugori, K., Yamaguchi, Y., et al. (2022). The 3’ Pol II pausing at replication-dependent histone genes is regulated by Mediator through Cajal bodies’ association with histone locus bodies. Nat Commun 13, 2905. 10.1038/s41467-022-30632-w.

53. Gu, J., Zhou, X., Sutherland, L., Kato, M., Jaczynska, K., Rizo, J., and McKnight, S.L. (2023). Oxidative regulation of TDP-43 self-association by a β-to-α conformational switch. Proc Natl Acad Sci U S A 120, e2311416120. 10.1073/pnas.2311416120.

54. Lu, H., Yu, D., Hansen, A.S., Ganguly, S., Liu, R., Heckert, A., Darzacq, X., and Zhou, Q. (2018). Phase-separation mechanism for C-terminal hyperphosphorylation of RNA polymerase II. Nature 558, 318–323. 10.1038/s41586-018-0174-3.

55. Titov, D.V., Gilman, B., He, Q.L., Bhat, S., Low, W.K., Dang, Y., Smeaton, M., Demain, A.L., Miller, P.S., Kugel, J.F., et al. (2011). XPB, a subunit of TFIIH, is a target of the natural product triptolide. Nat Chem Biol 7, 182–188. 10.1038/nchembio.522.

56. Wang, Y., Lu, J.J., He, L., and Yu, Q. (2011). Triptolide (TPL) inhibits global transcription by inducing proteasome-dependent degradation of RNA polymerase II (Pol II). PLoS One 6, e23993. 10.1371/journal.pone.0023993.

57. Papantonis, A., and Cook, P.R. (2013). Transcription factories: genome organization and gene regulation. Chem Rev 113, 8683–8705. 10.1021/cr300513p.

58. Moore, J.E., Pratt, H.E., Fan, K., Phalke, N., Fisher, J., Elhajjajy, S.I., Andrews, G., Gao, M., Shedd, N., Fu, Y., et al. (2026). An expanded registry of candidate cis-regulatory elements. Nature. 10.1038/s41586-025-09909-9.

59. Whyte, W.A., Orlando, D.A., Hnisz, D., Abraham, B.J., Lin, C.Y., Kagey, M.H., Rahl, P.B., Lee, T.I., and Young, R.A. (2013). Master transcription factors and mediator establish super-enhancers at key cell identity genes. Cell 153, 307–319. 10.1016/j.cell.2013.03.035.

60. Santos-Rosa, H., Schneider, R., Bannister, A.J., Sherriff, J., Bernstein, B.E., Emre, N.C., Schreiber, S.L., Mellor, J., and Kouzarides, T. (2002). Active genes are tri-methylated at K4 of histone H3. Nature 419, 407–411. 10.1038/nature01080.

61. Guenther, M.G., Levine, S.S., Boyer, L.A., Jaenisch, R., and Young, R.A. (2007). A chromatin landmark and transcription initiation at most promoters in human cells. Cell 130, 77–88. 10.1016/j.cell.2007.05.042.

62. Shah, R.N., Grzybowski, A.T., Cornett, E.M., Johnstone, A.L., Dickson, B.M., Boone, B.A., Cheek, M.A., Cowles, M.W., Maryanski, D., Meiners, M.J., et al. (2018). Examining the Roles of H3K4 Methylation States with Systematically Characterized Antibodies. Mol Cell 72, 162–177.e167. 10.1016/j.molcel.2018.08.015.

63. Creyghton, M.P., Cheng, A.W., Welstead, G.G., Kooistra, T., Carey, B.W., Steine, E.J., Hanna, J., Lodato, M.A., Frampton, G.M., Sharp, P.A., et al. (2010). Histone H3K27ac separates active from poised enhancers and predicts developmental state. Proc Natl Acad Sci U S A 107, 21931–21936. 10.1073/pnas.1016071107.

64. Rada-Iglesias, A., Bajpai, R., Swigut, T., Brugmann, S.A., Flynn, R.A., and Wysocka, J. (2011). A unique chromatin signature uncovers early developmental enhancers in humans. Nature 470, 279–283. 10.1038/nature09692.

65. Heintzman, N.D., Hon, G.C., Hawkins, R.D., Kheradpour, P., Stark, A., Harp, L.F., Ye, Z., Lee, L.K., Stuart, R.K., Ching, C.W., et al. (2009). Histone modifications at human enhancers reflect global cell-type-specific gene expression. Nature 459, 108–112. 10.1038/nature07829.

66. Cao, R., Wang, L., Wang, H., Xia, L., Erdjument-Bromage, H., Tempst, P., Jones, R.S., and Zhang, Y. (2002). Role of histone H3 lysine 27 methylation in Polycomb-group silencing. Science 298, 1039–1043. 10.1126/science.1076997.

67. Müller, J., Hart, C.M., Francis, N.J., Vargas, M.L., Sengupta, A., Wild, B., Miller, E.L., O’Connor, M.B., Kingston, R.E., and Simon, J.A. (2002). Histone methyltransferase activity of a Drosophila Polycomb group repressor complex. Cell 111, 197–208. 10.1016/s0092-8674(02)00976-5.

68. Yu, H., Zhao, J., Shen, Y., Qiao, L., Liu, Y., Xie, G., Chang, S., Ge, T., Li, N., Chen, M., et al. (2024). The dynamic landscape of enhancer-derived RNA during mouse early embryo development. Cell Rep 43, 114832. 10.1016/j.celrep.2024.114832.

69. Zeitlinger, J., Stark, A., Kellis, M., Hong, J.W., Nechaev, S., Adelman, K., Levine, M., and Young, R.A. (2007). RNA polymerase stalling at developmental control genes in the Drosophila melanogaster embryo. Nat Genet 39, 1512–1516. 10.1038/ng.2007.26.

70. Core, L.J., Waterfall, J.J., and Lis, J.T. (2008). Nascent RNA sequencing reveals widespread pausing and divergent initiation at human promoters. Science 322, 1845–1848. 10.1126/science.1162228.

71. Gilchrist, D.A., Dos Santos, G., Fargo, D.C., Xie, B., Gao, Y., Li, L., and Adelman, K. (2010). Pausing of RNA polymerase II disrupts DNA-specified nucleosome organization to enable precise gene regulation. Cell 143, 540–551. 10.1016/j.cell.2010.10.004.

72. Kwak, H., Fuda, N.J., Core, L.J., and Lis, J.T. (2013). Precise maps of RNA polymerase reveal how promoters direct initiation and pausing. Science 339, 950–953. 10.1126/science.1229386.

73. Wade, J.T., and Struhl, K. (2008). The transition from transcriptional initiation to elongation. Curr Opin Genet Dev 18, 130–136. 10.1016/j.gde.2007.12.008.

74. Yamaguchi, Y., Takagi, T., Wada, T., Yano, K., Furuya, A., Sugimoto, S., Hasegawa, J., and Handa, H. (1999). NELF, a multisubunit complex containing RD, cooperates with DSIF to repress RNA polymerase II elongation. Cell 97, 41–51. 10.1016/s0092-8674(00)80713-8.

75. Vos, S.M., Farnung, L., Boehning, M., Wigge, C., Linden, A., Urlaub, H., and Cramer, P. (2018). Structure of activated transcription complex Pol II-DSIF-PAF-SPT6. Nature 560, 607–612. 10.1038/s41586-018-0440-4.

76. Vos, S.M., Farnung, L., Urlaub, H., and Cramer, P. (2018). Structure of paused transcription complex Pol II-DSIF-NELF. Nature 560, 601–606. 10.1038/s41586-018-0442-2.

77. Jonkers, I., Kwak, H., and Lis, J.T. (2014). Genome-wide dynamics of Pol II elongation and its interplay with promoter proximal pausing, chromatin, and exons. Elife 3, e02407. 10.7554/eLife.02407.

78. Andersson, R., Gebhard, C., Miguel-Escalada, I., Hoof, I., Bornholdt, J., Boyd, M., Chen, Y., Zhao, X., Schmidl, C., Suzuki, T., et al. (2014). An atlas of active enhancers across human cell types and tissues. Nature 507, 455–461. 10.1038/nature12787.

79. Seila, A.C., Calabrese, J.M., Levine, S.S., Yeo, G.W., Rahl, P.B., Flynn, R.A., Young, R.A., and Sharp, P.A. (2008). Divergent transcription from active promoters. Science 322, 1849–1851. 10.1126/science.1162253.

80. Goel, V.Y., Huseyin, M.K., and Hansen, A.S. (2023). Region Capture Micro-C reveals coalescence of enhancers and promoters into nested microcompartments. Nat Genet 55, 1048–1056. 10.1038/s41588-023-01391-1.

81. Quinodoz, S.A., Ollikainen, N., Tabak, B., Palla, A., Schmidt, J.M., Detmar, E., Lai, M.M., Shishkin, A.A., Bhat, P., Takei, Y., et al. (2018). Higher-Order Inter-chromosomal Hubs Shape 3D Genome Organization in the Nucleus. Cell 174, 744–757.e724. 10.1016/j.cell.2018.05.024.

82. Weintraub, A.S., Li, C.H., Zamudio, A.V., Sigova, A.A., Hannett, N.M., Day, D.S., Abraham, B.J., Cohen, M.A., Nabet, B., Buckley, D.L., et al. (2017). YY1 Is a Structural Regulator of Enhancer-Promoter Loops. Cell 171, 1573–1588.e1528. 10.1016/j.cell.2017.11.008.

83. Dangkulwanich, M., Ishibashi, T., Liu, S., Kireeva, M.L., Lubkowska, L., Kashlev, M., and Bustamante, C.J. (2013). Complete dissection of transcription elongation reveals slow translocation of RNA polymerase II in a linear ratchet mechanism. Elife 2, e00971. 10.7554/eLife.00971.

84. Patel, A., Malinovska, L., Saha, S., Wang, J., Alberti, S., Krishnan, Y., and Hyman, A.A. (2017). ATP as a biological hydrotrope. Science 356, 753–756. 10.1126/science.aaf6846.

85. Linsenmeier, M., Hondele, M., Grigolato, F., Secchi, E., Weis, K., and Arosio, P. (2022). Dynamic arrest and aging of biomolecular condensates are modulated by low-complexity domains, RNA and biochemical activity. Nat Commun 13, 3030. 10.1038/s41467-022-30521-2.

86. Yewdall, N.A., André, A.A.M., van Haren, M.H.I., Nelissen, F.H.T., Jonker, A., and Spruijt, E. (2022). ATP:Mg. Biophys J 121, 3962–3974. 10.1016/j.bpj.2022.08.025.

87. Moore, C., Wong, E., Kaur, U., Chio, U.S., Zhou, Z., Ostrowski, M., Wu, K., Irkliyenko, I., Wang, S., Ramani, V., and Narlikar, G.J. (2025). ATP-dependent remodeling of chromatin condensates reveals distinct mesoscale outcomes. Science 390, eadr0018. 10.1126/science.adr0018.

88. Gwon, Y., Maxwell, B.A., Kolaitis, R.M., Zhang, P., Kim, H.J., and Taylor, J.P. (2021). Ubiquitination of G3BP1 mediates stress granule disassembly in a context-specific manner. Science 372, eabf6548. 10.1126/science.abf6548.

89. Zhang, M., Díaz-Celis, C., Onoa, B., Cañari-Chumpitaz, C., Requejo, K.I., Liu, J., Vien, M., Nogales, E., Ren, G., and Bustamante, C. (2022). Molecular organization of the early stages of nucleosome phase separation visualized by cryo-electron tomography. Mol Cell 82, 3000–3014.e3009. 10.1016/j.molcel.2022.06.032.

90. Zhou, H., Huertas, J., Maristany, M.J., Russell, K., Hwang, J.H., Yao, R.W., Samanta, N., Hutchings, J., Billur, R., Shiozaki, M., et al. (2025). Multiscale structure of chromatin condensates explains phase separation and material properties. Science 390, eadv6588. 10.1126/science.adv6588.

91. Li, G., Ruan, X., Auerbach, R.K., Sandhu, K.S., Zheng, M., Wang, P., Poh, H.M., Goh, Y., Lim, J., Zhang, J., et al. (2012). Extensive promoter-centered chromatin interactions provide a topological basis for transcription regulation. Cell 148, 84–98. 10.1016/j.cell.2011.12.014.

92. Jaeger, M.G., Schwalb, B., Mackowiak, S.D., Velychko, T., Hanzl, A., Imrichova, H., Brand, M., Agerer, B., Chorn, S., Nabet, B., et al. (2020). Selective Mediator dependence of cell-type-specifying transcription. Nat Genet 52, 719–727. 10.1038/s41588-020-0635-0.

93. Li, H., Dalgleish, J.L.T., Lister, G., Maristany, M.J., Huertas, J., Dopico-Fernandez, A.M., Hamley, J.C., Denny, N., Bloye, G., Zhang, W., et al. (2025). Mapping chromatin structure at base-pair resolution unveils a unified model of cis-regulatory element interactions. Cell 188, 7175–7193.e7119. 10.1016/j.cell.2025.10.013.

94. Plaschka, C., Larivière, L., Wenzeck, L., Seizl, M., Hemann, M., Tegunov, D., Petrotchenko, E.V., Borchers, C.H., Baumeister, W., Herzog, F., et al. (2015). Architecture of the RNA polymerase II-Mediator core initiation complex. Nature 518, 376–380. 10.1038/nature14229.

95. Abdella, R., Talyzina, A., Chen, S., Inouye, C.J., Tjian, R., and He, Y. (2021). Structure of the human Mediator-bound transcription preinitiation complex. Science 372, 52–56. 10.1126/science.abg3074.

96. Chen, X., Wang, X., Liu, W., Ren, Y., Qu, X., Li, J., Yin, X., and Xu, Y. (2022). Structures of +1 nucleosome-bound PIC-Mediator complex. Science 378, 62–68. 10.1126/science.abn8131.

97. Dobi, K.C., and Winston, F. (2007). Analysis of transcriptional activation at a distance in Saccharomyces cerevisiae. Mol Cell Biol 27, 5575–5586. 10.1128/MCB.00459-07.

98. Myers, L.C., Gustafsson, C.M., Bushnell, D.A., Lui, M., Erdjument-Bromage, H., Tempst, P., and Kornberg, R.D. (1998). The Med proteins of yeast and their function through the RNA polymerase II carboxy-terminal domain. Genes Dev 12, 45–54. 10.1101/gad.12.1.45.

99. Kagey, M.H., Newman, J.J., Bilodeau, S., Zhan, Y., Orlando, D.A., van Berkum, N.L., Ebmeier, C.C., Goossens, J., Rahl, P.B., Levine, S.S., et al. (2010). Mediator and cohesin connect gene expression and chromatin architecture. Nature 467, 430–435. 10.1038/nature09380.

100. Malik, S., and Roeder, R.G. (2010). The metazoan Mediator co-activator complex as an integrative hub for transcriptional regulation. Nat Rev Genet 11, 761–772. 10.1038/nrg2901.

101. Ma, L., Gao, Z., Wu, J., Zhong, B., Xie, Y., Huang, W., and Lin, Y. (2021). Co-condensation between transcription factor and coactivator p300 modulates transcriptional bursting kinetics. Mol Cell 81, 1682–1697.e1687. 10.1016/j.molcel.2021.01.031.

102. Muse, G.W., Gilchrist, D.A., Nechaev, S., Shah, R., Parker, J.S., Grissom, S.F., Zeitlinger, J., and Adelman, K. (2007). RNA polymerase is poised for activation across the genome. Nat Genet 39, 1507–1511. 10.1038/ng.2007.21.

103. Levo, M., Raimundo, J., Bing, X.Y., Sisco, Z., Batut, P.J., Ryabichko, S., Gregor, T., and Levine, M.S. (2022). Transcriptional coupling of distant regulatory genes in living embryos. Nature 605, 754–760. 10.1038/s41586-022-04680-7.

104. Narita, T., Ito, S., Higashijima, Y., Chu, W.K., Neumann, K., Walter, J., Satpathy, S., Liebner, T., Hamilton, W.B., Maskey, E., et al. (2021). Enhancers are activated by p300/CBP activity-dependent PIC assembly, RNAPII recruitment, and pause release. Mol Cell 81, 2166–2182 e2166. 10.1016/j.molcel.2021.03.008.

105. Kubo, N., Ishii, H., Xiong, X., Bianco, S., Meitinger, F., Hu, R., Hocker, J.D., Conte, M., Gorkin, D., Yu, M., et al. (2021). Promoter-proximal CTCF binding promotes distal enhancer-dependent gene activation. Nat Struct Mol Biol 28, 152–161. 10.1038/s41594-020-00539-5.

106. Illingworth, R.S., Moffat, M., Mann, A.R., Read, D., Hunter, C.J., Pradeepa, M.M., Adams, I.R., and Bickmore, W.A. (2015). The E3 ubiquitin ligase activity of RING1B is not essential for early mouse development. Genes Dev 29, 1897–1902. 10.1101/gad.268151.115.

107. Shao, R., Kumar, B., Lidschreiber, K., Lidschreiber, M., Cramer, P., and Elsasser, S.J. (2022). Distinct transcription kinetics of pluripotent cell states. Mol Syst Biol 18, e10407. 10.15252/msb.202110407.

108. Wang, Q., Li, M., Wu, T., Zhan, L., Li, L., Chen, M., Xie, W., Xie, Z., Hu, E., Xu, S., and Yu, G. (2022). Exploring Epigenomic Datasets by ChIPseeker. Curr Protoc 2, e585. 10.1002/cpz1.585.

109. Yu, G., Wang, L.G., and He, Q.Y. (2015). ChIPseeker: an R/Bioconductor package for ChIP peak annotation, comparison and visualization. Bioinformatics 31, 2382–2383. 10.1093/bioinformatics/btv145.

110. Dreos, R., Ambrosini, G., Cavin Perier, R., and Bucher, P. (2013). EPD and EPDnew, high-quality promoter resources in the next-generation sequencing era. Nucleic Acids Res 41, D157–164. 10.1093/nar/gks1233.

111. Noguchi, S., Arakawa, T., Fukuda, S., Furuno, M., Hasegawa, A., Hori, F., Ishikawa-Kato, S., Kaida, K., Kaiho, A., Kanamori-Katayama, M., et al. (2017). FANTOM5 CAGE profiles of human and mouse samples. Sci Data 4, 170112. 10.1038/sdata.2017.112.

112. Consortium, F., the, R.P., Clst, Forrest, A.R., Kawaji, H., Rehli, M., Baillie, J.K., de Hoon, M.J., Haberle, V., Lassmann, T., et al. (2014). A promoter-level mammalian expression atlas. Nature 507, 462–470. 10.1038/nature13182.

113. Yeom, K.H., Pan, Z., Lin, C.H., Lim, H.Y., Xiao, W., Xing, Y., and Black, D.L. (2021). Tracking pre-mRNA maturation across subcellular compartments identifies developmental gene regulation through intron retention and nuclear anchoring. Genome Res 31, 1106–1119. 10.1101/gr.273904.120.

114. Grimm, J.B., Xie, L., Casler, J.C., Patel, R., Tkachuk, A.N., Falco, N., Choi, H., Lippincott-Schwartz, J., Brown, T.A., Glick, B.S., et al. (2021). A General Method to Improve Fluorophores Using Deuterated Auxochromes. JACS Au 1, 690–696. 10.1021/jacsau.1c00006.

115. Ruthenburg, A.J., Li, H., Milne, T.A., Dewell, S., McGinty, R.K., Yuen, M., Ueberheide, B., Dou, Y., Muir, T.W., Patel, D.J., and Allis, C.D. (2011). Recognition of a mononucleosomal histone modification pattern by BPTF via multivalent interactions. Cell 145, 692–706. 10.1016/j.cell.2011.03.053.

116. Danecek, P., Bonfield, J.K., Liddle, J., Marshall, J., Ohan, V., Pollard, M.O., Whitwham, A., Keane, T., McCarthy, S.A., Davies, R.M., and Li, H. (2021). Twelve years of SAMtools and BCFtools. Gigascience 10. 10.1093/gigascience/giab008.

117. Kim, D., Paggi, J.M., Park, C., Bennett, C., and Salzberg, S.L. (2019). Graph-based genome alignment and genotyping with HISAT2 and HISAT-genotype. Nat Biotechnol 37, 907–915. 10.1038/s41587-019-0201-4.

118. Ramirez, F., Ryan, D.P., Gruning, B., Bhardwaj, V., Kilpert, F., Richter, A.S., Heyne, S., Dundar, F., and Manke, T. (2016). deepTools2: a next generation web server for deep-sequencing data analysis. Nucleic Acids Res 44, W160–165. 10.1093/nar/gkw257.

119. Amemiya, H.M., Kundaje, A., and Boyle, A.P. (2019). The ENCODE Blacklist: Identification of Problematic Regions of the Genome. Sci Rep 9, 9354. 10.1038/s41598-019-45839-z.

120. Zhang, Y., Liu, T., Meyer, C.A., Eeckhoute, J., Johnson, D.S., Bernstein, B.E., Nusbaum, C., Myers, R.M., Brown, M., Li, W., and Liu, X.S. (2008). Model-based analysis of ChIP-Seq (MACS). Genome Biol 9, R137. 10.1186/gb-2008-9-9-r137.

121. Lawrence, M., Huber, W., Pagès, H., Aboyoun, P., Carlson, M., Gentleman, R., Morgan, M.T., and Carey, V.J. (2013). Software for computing and annotating genomic ranges. PLoS Comput Biol 9, e1003118. 10.1371/journal.pcbi.1003118.

122. Wilson, T. (2022). mm10.mappability_score. DOI:10.17632/jr36ntmzsh.1.

123. Micallef, L., and Rodgers, P. (2014). eulerAPE: drawing area-proportional 3-Venn diagrams using ellipses. PLoS One 9, e101717. 10.1371/journal.pone.0101717.

124. Wilkinson, L. (2012). Exact and approximate area-proportional circular Venn and Euler diagrams. IEEE Trans Vis Comput Graph 18, 321–331. 10.1109/TVCG.2011.56.

125. Gel, B., Diez-Villanueva, A., Serra, E., Buschbeck, M., Peinado, M.A., and Malinverni, R. (2016). regioneR: an R/Bioconductor package for the association analysis of genomic regions based on permutation tests. Bioinformatics 32, 289–291. 10.1093/bioinformatics/btv562.

126. Quinlan, A.R., and Hall, I.M. (2010). BEDTools: a flexible suite of utilities for comparing genomic features. Bioinformatics 26, 841–842. 10.1093/bioinformatics/btq033.

127. Team BC, M.B. (2019). _TxDb.Mmusculus.UCSC.mm10.knownGene: Annotation package for TxDb object(s)_. R package version 3.10.0.

128. org.Mm.eg.db. DOI: 10.18129/B9.bioc.org.Mm.eg.db.

129. Lawrence, M., Gentleman, R., and Carey, V. (2009). rtracklayer: an R package for interfacing with genome browsers. Bioinformatics 25, 1841–1842. 10.1093/bioinformatics/btp328.

130. Dreos, R., Ambrosini, G., Groux, R., Cavin Perier, R., and Bucher, P. (2017). The eukaryotic promoter database in its 30th year: focus on non-vertebrate organisms. Nucleic Acids Res 45, D51–D55. 10.1093/nar/gkw1069.

131. Frankish, A., Carbonell-Sala, S., Diekhans, M., Jungreis, I., Loveland, J.E., Mudge, J.M., Sisu, C., Wright, J.C., Arnan, C., Barnes, I., et al. (2023). GENCODE: reference annotation for the human and mouse genomes in 2023. Nucleic Acids Res 51, D942–D949. 10.1093/nar/gkac1071.

132. Liao, Y., Smyth, G.K., and Shi, W. (2014). featureCounts: an efficient general purpose program for assigning sequence reads to genomic features. Bioinformatics 30, 923–930. 10.1093/bioinformatics/btt656.

133. Love, M.I., Huber, W., and Anders, S. (2014). Moderated estimation of fold change and dispersion for RNA-seq data with DESeq2. Genome Biol 15, 550. 10.1186/s13059-014-0550-8.

134. Zhu, A., Ibrahim, J.G., and Love, M.I. (2019). Heavy-tailed prior distributions for sequence count data: removing the noise and preserving large differences. Bioinformatics 35, 2084–2092. 10.1093/bioinformatics/bty895.

135. Xu, S., Hu, E., Cai, Y., Xie, Z., Luo, X., Zhan, L., Tang, W., Wang, Q., Liu, B., Wang, R., et al. (2024). Using clusterProfiler to characterize multiomics data. Nat Protoc 19, 3292–3320. 10.1038/s41596-024-01020-z.

136. Gene Ontology, C., Aleksander, S.A., Balhoff, J., Carbon, S., Cherry, J.M., Drabkin, H.J., Ebert, D., Feuermann, M., Gaudet, P., Harris, N.L., et al. (2023). The Gene Ontology knowledgebase in 2023. Genetics 224. 10.1093/genetics/iyad031.

137. Subramanian, A., Tamayo, P., Mootha, V.K., Mukherjee, S., Ebert, B.L., Gillette, M.A., Paulovich, A., Pomeroy, S.L., Golub, T.R., Lander, E.S., and Mesirov, J.P. (2005). Gene set enrichment analysis: a knowledge-based approach for interpreting genome-wide expression profiles. Proc Natl Acad Sci U S A 102, 15545–15550. 10.1073/pnas.0506580102.

138. Castanza, A.S., Recla, J.M., Eby, D., Thorvaldsdottir, H., Bult, C.J., and Mesirov, J.P. (2023). Extending support for mouse data in the Molecular Signatures Database (MSigDB). Nat Methods 20, 1619–1620. 10.1038/s41592-023-02014-7.

139. Dolgalev, I. (2026). msigdbr: MSigDB Gene Sets for Multiple Organisms in a Tidy Data Format. R package version 25.1.1.

140. Liberzon, A., Birger, C., Thorvaldsdottir, H., Ghandi, M., Mesirov, J.P., and Tamayo, P. (2015). The Molecular Signatures Database (MSigDB) hallmark gene set collection. Cell Syst 1, 417–425. 10.1016/j.cels.2015.12.004.

141. Mikkelsen, T.S., Hanna, J., Zhang, X., Ku, M., Wernig, M., Schorderet, P., Bernstein, B.E., Jaenisch, R., Lander, E.S., and Meissner, A. (2008). Dissecting direct reprogramming through integrative genomic analysis. Nature 454, 49–55. 10.1038/nature07056.

142. Eisenberg, E., and Levanon, E.Y. (2013). Human housekeeping genes, revisited. Trends Genet 29, 569–574. 10.1016/j.tig.2013.05.010.

143. Novo, C.L., Javierre, B.M., Cairns, J., Segonds-Pichon, A., Wingett, S.W., Freire-Pritchett, P., Furlan-Magaril, M., Schoenfelder, S., Fraser, P., and Rugg-Gunn, P.J. (2018). Long-Range Enhancer Interactions Are Prevalent in Mouse Embryonic Stem Cells and Are Reorganized upon Pluripotent State Transition. Cell Rep 22, 2615–2627. 10.1016/j.celrep.2018.02.040.

144. Perez, G., Barber, G.P., Benet-Pages, A., Casper, J., Clawson, H., Diekhans, M., Fischer, C., Gonzalez, J.N., Hinrichs, A.S., Lee, C.M., et al. (2025). The UCSC Genome Browser database: 2025 update. Nucleic Acids Res 53, D1243–D1249. 10.1093/nar/gkae974.

145. Csárdi G, N.T. (2006). The igraph software package for complex network research.

146. Csárdi G, N.T., Traag V, Horvát S, Zanini F, Noom D, Müller K, Schoch D, Salmon M (2026). igraph: Network Analysis and Visualization in R.

147. Antonov M, C.G., Horvát S, Müller K, Nepusz T, Noom D, Salmon M, Traag V, Welles BF, Zanini F igraph enables fast and robust network analysis across programming languages. arXiv preprint doi:10.48550/arXiv.2311.10260.

148. Archit, A., Freckmann, L., Nair, S., Khalid, N., Hilt, P., Rajashekar, V., Freitag, M., Teuber, C., Spitzner, M., Tapia Contreras, C., et al. (2025). Segment Anything for Microscopy. Nat Methods 22, 579–591. 10.1038/s41592-024-02580-4.

149. Haase, R. (2026). napari-segment-blobs-and-things-with-membranes [Computer software].

150. Xu, W., Cai, H., Zhang, Q., Wang, Z., Yang, J., Wu, X., Li, C., Cui, C., Liu, C., He, J., et al. (2025). U-FISH: a fluorescent spot detector for imaging-based spatial-omics analysis and AI-assisted FISH diagnosis. Genome Biol 26, 261. 10.1186/s13059-025-03736-x.

151. van der Walt, S., Schonberger, J.L., Nunez-Iglesias, J., Boulogne, F., Warner, J.D., Yager, N., Gouillart, E., Yu, T., and scikit-image, c. (2014). scikit-image: image processing in Python. PeerJ 2, e453. 10.7717/peerj.453.

152. Virtanen, P., Gommers, R., Oliphant, T.E., Haberland, M., Reddy, T., Cournapeau, D., Burovski, E., Peterson, P., Weckesser, W., Bright, J., et al. (2020). SciPy 1.0: fundamental algorithms for scientific computing in Python. Nat Methods 17, 261–272. 10.1038/s41592-019-0686-2.

153. Comet, M., Dijkman, P.M., Boer Iwema, R., Franke, T., Masiulis, S., Schampers, R., Raschdorf, O., Grollios, F., Pryor, E.E., Jr., and Drulyte, I. (2024). Tomo Live: an on-the-fly reconstruction pipeline to judge data quality for cryo-electron tomography workflows. Acta Crystallogr D Struct Biol 80, 247–258. 10.1107/S2059798324001840.

154. Zheng, S.Q., Palovcak, E., Armache, J.P., Verba, K.A., Cheng, Y., and Agard, D.A. (2017). MotionCor2: anisotropic correction of beam-induced motion for improved cryo-electron microscopy. Nat Methods 14, 331–332. 10.1038/nmeth.4193.

155. Gilbert, P. (1972). Iterative methods for the three-dimensional reconstruction of an object from projections. J Theor Biol 36, 105–117. 10.1016/0022-5193(72)90180-4.

156. Burnley, T., Palmer, C.M., and Winn, M. (2017). Recent developments in the CCP-EM software suite. Acta Crystallogr D Struct Biol 73, 469–477. 10.1107/S2059798317007859.

